# Influences of rare copy number variation on human complex traits

**DOI:** 10.1101/2021.10.21.465308

**Authors:** Margaux L.A. Hujoel, Maxwell A. Sherman, Alison R. Barton, Ronen E. Mukamel, Vijay G. Sankaran, Po-Ru Loh

## Abstract

The human genome contains hundreds of thousands of regions exhibiting copy number variation (CNV). However, the phenotypic effects of most such polymorphisms are unknown because only larger CNVs (spanning tens of kilobases) have been ascertainable from the SNP-array data generated by large biobanks. We developed a new computational approach that leverages abundant haplotype-sharing in biobank cohorts to more sensitively detect CNVs co-inherited within extended SNP haplotypes. Applied to UK Biobank, this approach achieved 6-fold increased CNV detection sensitivity compared to previous analyses, accounting for approximately half of all rare gene inactivation events produced by genomic structural variation. This extensive CNV call set enabled the most comprehensive analysis to date of associations between CNVs and 56 quantitative traits, identifying 269 independent associations (*P* < 5 × 10^−8^) – involving 97 loci – that rigorous statistical fine-mapping analyses indicated were likely to be causally driven by CNVs. Putative target genes were identifiable for nearly half of the loci, enabling new insights into dosage-sensitivity of these genes and implicating several novel gene-trait relationships. CNVs at several loci created extended allelic series including deletions or duplications of distal enhancers that associated with much stronger phenotypic effects than SNPs within these regulatory elements. These results demonstrate the ability of haplotype-informed analysis to empower structural variant detection and provide insights into the genetic basis of human complex traits.

## Introduction

Copy number variants (CNVs), which duplicate and delete 50 base pair to megabase-scale genomic segments throughout the human genome^1–3^, are known to contribute to numerous genomic disorders including neuropsychiatric diseases^4–6^ and have been estimated to account for a considerable fraction of all rare loss-of-function (LoF) events affecting protein-coding genes^2^. Beyond disrupting coding sequences of genes, CNVs can also have unique functional consequences not producible by SNPs: for example, duplications can increase gene dosage, and deletions can eliminate regulatory elements. Investigating the broader phenotypic impacts of CNVs thus has the potential to uncover new large-effect variants and further our understanding of the genetic architecture of complex traits.

However, well-powered, phenome-wide CNV association analyses to date have been limited to considering large CNVs (tens of kilobases or longer) detectable from low-cost SNP-array data^7^ available for biobank-scale cohorts. Moreover, CNV association studies have encountered analytical challenges such as how to harmonize imprecise breakpoints of CNV calls, how to group CNVs for association testing, and how to filter associations that only reflect linkage disequilibrium with nearby SNPs. Despite these difficulties, previous studies have made many important discoveries both by investigating the role of known pathogenic CNVs on various phenotypes^8–10^ and by conducting association analysis on all CNVs detected in large cohorts^11–17^, including UK Biobank^18^. Here we developed a more sensitive CNV-detection method leveraging haplotype-sharing within biobank cohorts and applied it to UK Biobank, empowering exploration of the phenotypic effects of CNVs at much higher resolution than previously possible.

## Results

### Haplotype-informed copy-number variant detection

We developed a novel computational approach to CNV detection, called HI-CNV (**<u>H</u>**aplotype-**<u>I</u>**nformed **<u>C</u>**opy-**<u>N</u>**umber-**<u>V</u>**ariation), that substantially increases CNV detection power in large cohorts by pooling information across individuals who share extended SNP haplotypes. The intuition behind this approach is that in large biobank cohorts, population-polymorphic CNVs are usually carried by multiple individuals who co-inherited a CNV on a shared haplotype originating from a common ancestor. As such, power to detect a CNV can be increased by sharing information about its presence (e.g., from genotyping array intensity data) across multiple carriers (Fig. 1a).

**Figure 1:**
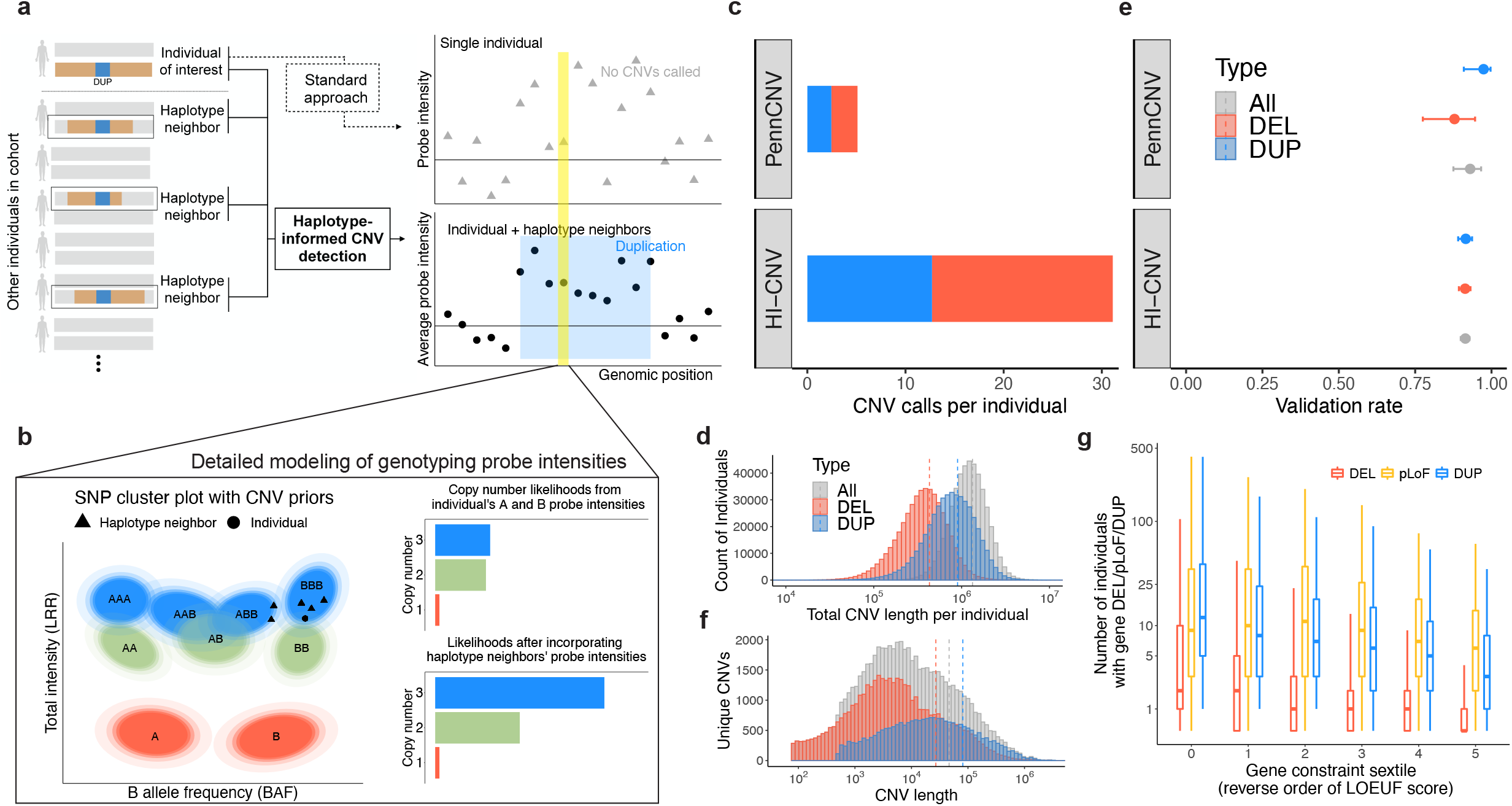
Haplotype-informed CNV detection from SNP-array data in UK Biobank. **a**. The HI-CNV framework improves power to detect CNVs by analyzing SNP-array data from an individual together with corresponding data from individuals with long shared haplotypes (“haplotype neighbors”). In contrast, standard approaches analyze data from the individual alone. **b**. SNP-specific genotype cluster priors map allele-specific (A and B allele) probe intensity measurements to probabilistic information about copy-number likelihoods. **c**. Average number of CNVs called by PennCNV and HI-CNV per UK Biobank participant. **d**. Distribution of total CNV length per individual in the HI-CNV call set. **e**. Validation rate of CNV calls from PennCNV and HI-CNV on 43 UK Biobank participants with independent whole-genome sequencing data. Error bars, 95% CIs. **f**. Distribution of CNV lengths in the HI-CNV call set. **g**. Distributions (across increasingly constrained gene sets) of observed counts of whole-gene deletions and duplications and pLoF CNVs in n=452,500 UK Biobank participants. Centers, medians; box edges, 25^th^ and 75^th^ percentiles; whiskers, 5^th^ and 95^th^ percentiles.

To identify individuals who are likely to share a segment of genome inherited from a recent common ancestor (and therefore likely to have co-inherited any CNVs contained within the shared genomic tract), we adapted recent approaches that use the positional Burrows-Wheeler transform (PBWT)^19^ to rapidly identify identity-by-descent (IBD) segments^20^. Specifically, for each haplotype of each individual in a cohort, we use a PBWT-based algorithm to identify its closest “haplotype neighbors” – i.e., the longest IBD matches with other haplotypes in the cohort – spanning each genomic position (Fig. 1a). Then, given quantitative information about the potential presence of a CNV in genetic data from the individual, as well as corresponding information from “haplotype neighbors,” we use a hidden Markov model (HMM) to detect CNVs co-inherited on shared haplotypes.

To apply our HI-CNV approach to SNP-array genotyping probe intensity data available for the UK Biobank cohort, we further developed methods to learn probabilistic models that map allele-specific probe intensity measurements to probabilistic information about copy-number likelihoods (Fig. 1b). Intuitively, genotyping probes within CNVs produce distinctive intensity measurements compared to probes not within CNVs. While these deviations are difficult to detect given data from one SNP, the signal becomes clearer when consistent deviations are observed across multiple consecutive SNPs^7^ – or, for HI-CNV, across multiple individuals co-inheriting a CNV. To optimize signal available from SNP-array probe intensities, we estimated SNP-specific genotype cluster priors corresponding to nine possible genotypes across copy-number states 1 (deletion), 2, and 3 (duplication) (Fig. 1b), and we also denoised total intensities using principal component analysis. Full methodological details are provided in Methods and the Supplementary Note.

### Modeling haplotype sharing increases CNV detection power in UK Biobank

We applied HI-CNV to detect CNVs across 452,500 UK Biobank participants of European ancestry. HI-CNV detected >6 times as many CNVs per individual as the widely-used PennCNV method (Fig. 1c), producing an average of 31.1 CNV calls per individual (18.4 deletions and 12.7 duplications spanning an average of 430kb and 899kb, respectively; Fig. 1c,d and Supplementary Table 1). In contrast, previous PennCNV-based analyses of UK Biobank SNP-array intensity data produced ∼ 4-6 CNV calls per individual depending on quality-control filters applied^8,12^. Validation analyses using whole-genome sequencing (WGS) pilot data available for 43 participants estimated a validation rate of 91% for HI-CNV, similar to that of PennCNV (Methods, Fig. 1e, and Supplementary Table 2).

HI-CNV’s increased detection sensitivity was driven by improved ability to detect CNVs on the scale of 10kb or shorter (Fig. 1f; Supplementary Table 3), which account for the majority of all CNVs^1–3^ but have traditionally been difficult to detect from SNP-array data. We designed HI-CNV with the goal of sensitively detecting low-frequency and rare CNVs of length >5kb (versus ∼ 50kb for previous SNP-array-based analyses of UK Biobank), focusing on CNVs with minor allele frequency (MAF) < 5% because of their potential to be more deleterious and because SNP-array designs tend to avoid common CNV regions. Among such CNVs called from WGS pilot data and spanning at least two SNP-array probes (the minimum required by our approach), HI-CNV achieved a recall rate of 81% (Supplementary Fig. 1 and Supplementary Table 4). Recall was unsurprisingly much lower (6%) when considering all MAF<5% CNVs called from WGS data (i.e., removing restrictions on size and array-overlap), consistent with most CNVs being shorter than the resolution of SNP-array probe spacing. However, recall of gene-overlapping CNVs was substantially higher (24%) because the UK Biobank SNP-array was designed to prioritize inclusion of coding variants^18^. Moreover, the HI-CNV call set appeared to account for approximately half of the 10.2 genes per genome estimated to be altered by rare structural variants^2^: restricting to rare (MAF < 1%) whole-gene duplications and CNVs predicted to cause loss-of-function (pLoF), a mean of 5.0 genes per individual were altered by such CNVs (2.8 pLoF and 2.2 gene duplications). Across 18,251 genes, whole-gene duplications and pLoF CNVs were called in a median of 6 and 8 individuals, respectively, with observed counts decreasing with increasing gene constraint (Fig. 1g).

### Fine-mapping analyses reveal likely-causal CNV-trait associations

HI-CNV’s detection of many previously-undiscovered CNVs in UK Biobank suggested that CNV-phenotype association analyses might uncover new CNVs impacting human traits. We applied a combination of single-variant and burden-style analyses to test three categories of CNVs (gene-level, CNV-level, and probe-level; Fig. 2a) for association with 56 heritable quantitative traits, including anthropometric traits, blood pressure, measures of lung function, bone mineral density, blood cell indices, and serum biomarkers (Supplementary Data 1). We performed association analyses on up to 452,500 UK Biobank participants of European ancestry using linear mixed models implemented in BOLT-LMM^21,22^. We then removed associations that could potentially be explained by linkage disequilibrium with other variants by requiring each association to remain significant (P < 5 × 10^−8^) after conditioning on any other more-strongly-associated SNP, indel, or CNV within 3 megabases (Methods). We previously observed that when fine-mapping associations involving rare variants (which comprised nearly all CNVs we detected; Supplementary Fig. 2 and Supplementary Table 5), this pairwise LD filter effectively identifies variants likely to causally drive associations^23^. This analysis pipeline resulted in 269 fine-mapped CNV-trait associations at 97 loci involving 252 likely-causal CNVs (Supplementary Data 2 and 3).

**Figure 2:**
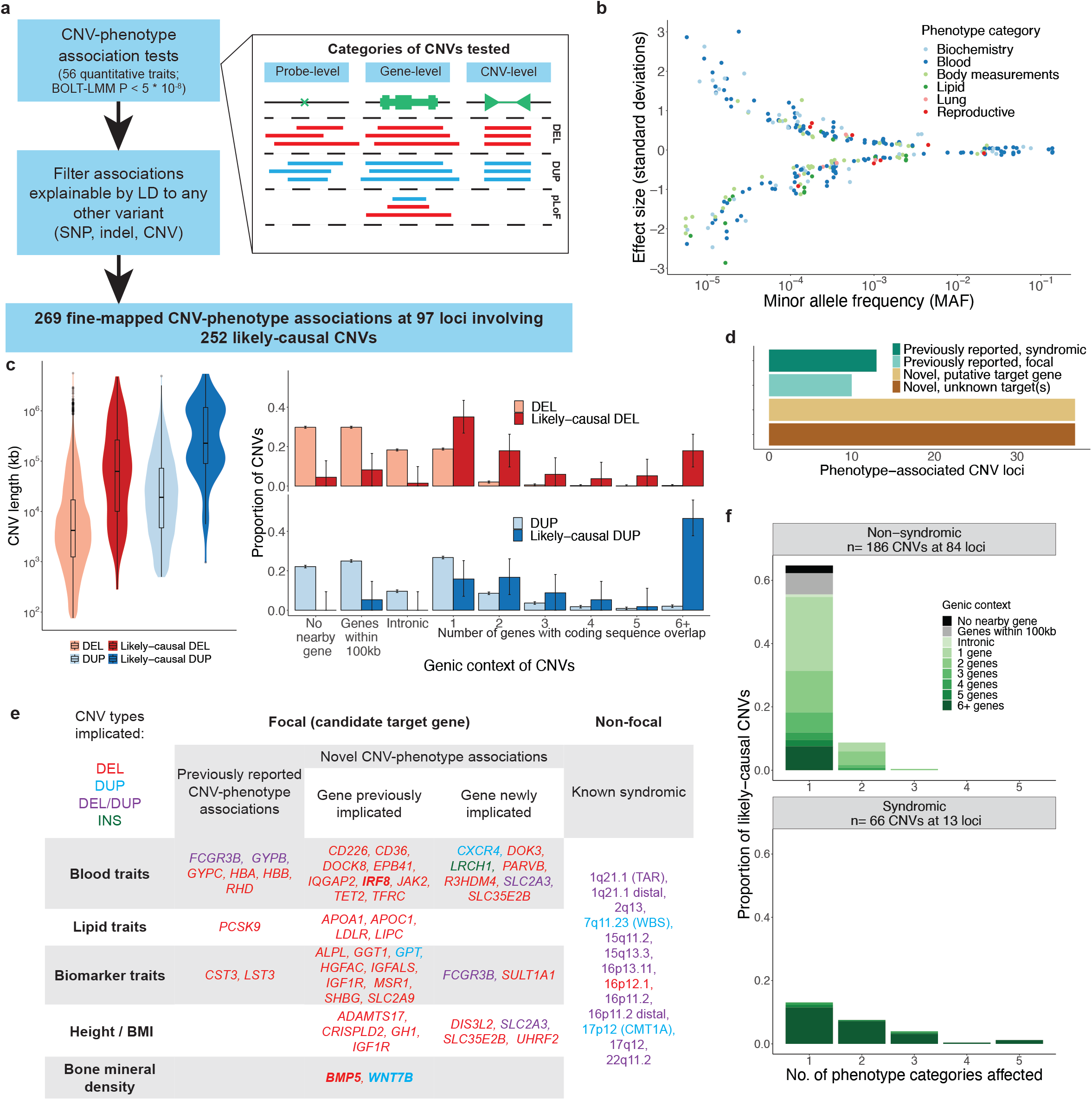
Fine-mapping analyses reveal likely-causal CNV-trait associations. **a**. Association and fine-mapping pipeline; inset depicts the three categories of CNVs tested. **b**. Effect size versus minor allele frequency for 269 likely-causal CNV-phenotype associations, colored by phenotype category. **c**. Distributions of CNV length (left) and genic context (right) across all CNVs and across likely-causal CNVs. **d**. Breakdown of 97 CNV loci according to prior literature status and (for novel loci) whether a putative target gene was identified. **e**. Candidate target genes, categorized according to whether (i) the CNV-phenotype association was previously reported, (ii) the target gene was previously implicated (either by a previously-reported coding variant association or by previous experimental work), or (iii) the gene is newly implicated for the phenotype. The rightmost column lists syndromic CNVs re-identified here. Colors indicate CNV type; bold font indicates noncoding CNVs potentially regulating the target gene. **f**. Genic context of syndromic CNVs (bottom) and non-syndromic CNVs (top) stratified by the number of phenotype categories associated with the CNV.

Many of the 269 likely-causal CNV-phenotype associations had large effect sizes – including 59 associations with an absolute effect size greater than 1 standard deviation (s.d.) – and effect sizes generally increased with decreasing minor allele frequency (MAF) (Fig. 2b). Only 10 of the 269 associations involved common (MAF > 5%) CNVs, whereas 186 associations involved CNVs with MAF < 0.1%. The associations affected most categories of phenotypes we considered, with blood cell phenotypes accounting for the majority of likely-causal associations (137 of 269 associations).

The likely-causal CNV-phenotype associations involved at least 252 unique CNVs (138 deletions, 114 duplications; Methods and Supplementary Data 3) which were enriched for multiple attributes correlated with functional impact (Fig. 2c). Likely-causal CNVs tended to be longer than average^13^ and were much more likely to overlap coding sequences of genes (85.8% coding-overlapping vs. 22.1% expected for deletions; 94.7% vs. 43.4% expected for duplications; Fig. 2c and Supplementary Table 6). For the small fraction of likely-causal deletions that did not overlap coding sequence (14.2%), roughly half overlapped enhancer annotations (42.1% vs. 8.4% expected; P = 7.76 × 10^−5^) (Supplementary Table 7). The majority of likely-causal deletions affected either one gene (35%) or two genes (18%), facilitating further investigation of potential targets of trait-modifying CNVs.

### CNV loci corroborate SNP associations and implicate new genes

Of the 97 loci involved in the 269 fine-mapped CNV-trait associations, 74 loci represented novel (to our knowledge) CNV-trait associations (Fig. 2d). In assessing novelty, we considered all previously published large-scale CNV association studies of which we were aware^6,10–15,17^, including previous analyses of UK Biobank in which CNVs were genotyped using PennCNV^10,12^ (which did not detect most likely-causal CNVs smaller than 100 kb; Supplementary Fig. 3). For half of the novel CNV loci (37 of 74 loci), we could identify a putative target gene (Fig. 2d,e and Supplementary Data 3). Among the 23 previously reported loci, roughly half (13 loci) corresponded to syndromic CNVs known to cause genetic disorders (Methods). These CNVs generally affected more phenotype categories and overlapped more genes than CNVs at non-syndromic loci (Fig. 2f), as expected.

Many CNV associations corroborated target genes recently implicated by coding variant association studies^14,23,24^, including rare height-reducing deletions in *CRISPLD2* and *ADAMTS17*, a rare sex hormone binding globulin (SHBG)-increasing deletion in *HGFAC*, and a rare IGF-1-decreasing partial deletion of *MSR1* (Fig. 2e). Other CNVs altered genes with known function but for which effects of population-polymorphic variants have not previously been described, such as *TFRC* (encoding transferrin receptor protein 1)^25,26^. Ultra-rare CNVs predicted to result in *TFRC* loss-of-function (pLoF) were found in 15 individuals and associated with 2.24 (s.e. 0.22) s.d. lower mean corpuscular hemoglobin and increased risk of iron deficiency anemia (OR = 4.9 (95% CI, 1.2-18.1); P = 0.034, Fisher’s exact test among unrelated participants). Several other CNV associations newly implicated genes contributing to the architecture of complex traits (Fig. 2e). Given the large number of novel CNV loci identified here, we focus below on describing three classes of particularly interesting loci: (1) CNV associations stronger than any nearby SNP, (2) loci at which CNVs, together with nearby SNPs, created long allelic series, and (3) additional loci newly implicating putative target genes.

### New CNV associations stronger than nearby SNPs

Among 169 associations involving non-syndromic CNVs, a subset of 37 associations (22%) were stronger than associations of all SNPs within 500kb. Several of these associations implicated novel gene-trait relationships; here we highlight two loci with such associations. First, ultra-rare *UHRF2* pLoF CNVs (carried by 19 UK Biobank participants) associated with a 1.11 (0.17) s.d. decrease in height (corresponding to 7.2 (1.1) cm shorter stature; P = 8.2 × 10^−11^; Fig. 3a and Supplementary Table 8). This association between *UHRF2* and height was not visible from SNPs at the locus, none of which reached genome-wide significance (Fig. 3a).

**Figure 3:**
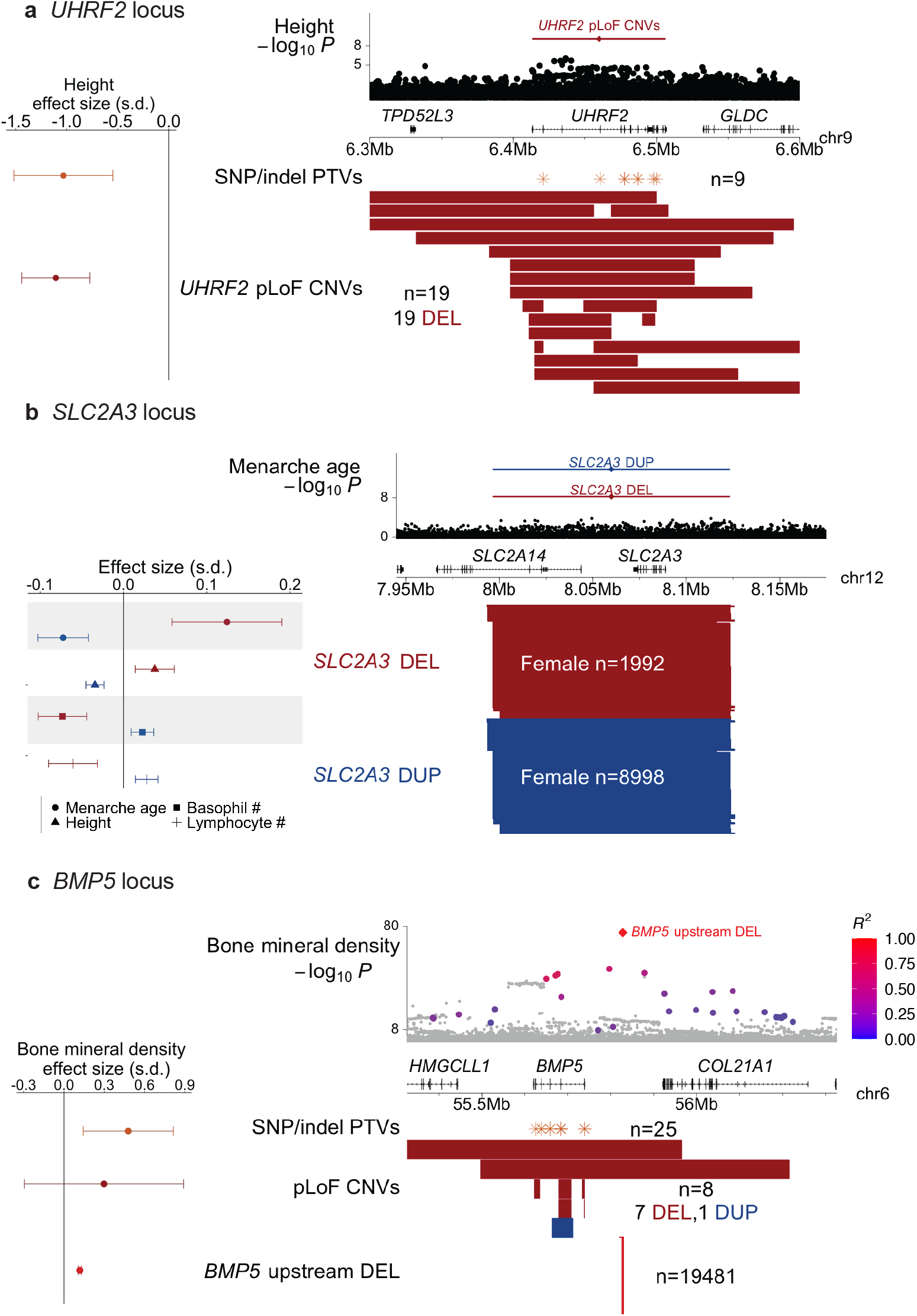
New CNV-phenotype associations stronger than nearby SNPs. **a. *UHRF2* locus**. Top: height associations for *UHRF2* pLoF CNVs and nearby SNPs. Bottom: locations of *UHRF2* pLoF CNVs and SNP and indel PTVs; left: effect sizes for height. **b. *SLC2A3* locus**. Top: menarche age associations for *SLC2A3* duplications and deletions and nearby SNPs. Bottom: locations of *SLC2A3* deletions and duplications; left: effect sizes for menarche age, height, and basophil and lymphocyte counts. **c. *BMP5* locus**. Top: bone mineral density associations for a deletion upstream of *BMP5* and nearby SNPs (colored according to linkage disequilibrium with the deletion, for SNPs with *R*^2^>0.1 to the deletion). Bottom: locations of the upstream deletion, *BMP5* pLoF CNVs, and SNP and indel PTVs; left: effect sizes for bone mineral density. In all panels, deletions are colored red and duplications are colored blue. Error bars on effect sizes, 95% CIs. Numerical results are available in Supplementary Tables 8-10.

However, among 185,365 exome-sequenced UK Biobank participants^27^, nine carriers of *UHRF2* protein-truncating SNP or indel variants (PTVs) exhibited 1.03 (0.25) s.d. decreased height (P = 3 × 10^−5^), corroborating the CNV association (Fig. 3a, Methods, and Supplementary Information). *UHRF2* has not previously been implicated in large genome-wide association studies of height, demonstrating the utility of CNV association studies and motivating further study of how loss of one functional copy of *UHRF2* (which encodes an E3 ubiquitin-protein ligase) impairs growth.

Another set of associations implicated copy-number variation of *SLC2A3* as a modifier of age at menarche (P = 1.6 × 10^−17^), height (P = 7.7 × 10^−12^), and blood count phenotypes (Fig. 3b and Supplementary Data 2). *SLC2A3* encodes GLUT3, a glucose transporter expressed in multiple tissues, and is prone to non-allelic homologous recombination that produces gene dosage-modifying ∼ 130kb duplications and deletions (MAF = 1.9% and 0.4%, respectively, in our call set). *SLC2A3* CNVs have been observed in many earlier studies, several of which have reported nominally significant associations with various clinical phenotypes; however, replication of these associations has been mixed^28^. In UK Biobank, *SLC2A3* deletions associated with delayed menarche (0.20 (0.03) years), increased height (0.25 (0.08) cm), and decreased basophil and lymphocyte counts, while duplications associated with reciprocal effects of roughly half the magnitude (Fig. 3b and Supplementary Table 9). No individuals carried zero *SLC2A3* copies (vs. 7.9 such individuals expected; P = 0.0009), consistent with previous literature suggesting that homozygous LoF mutations may be incompatible with life^28,29^ (Supplementary Fig. 4). These results support a dosage-sensitive role of GLUT3 in multiple organ systems.

Several other associations provided examples of loci at which SNP associations appeared to tag more-strongly-associated CNVs. Among the 37 associations for which a non-syndromic CNV attained the strongest association within 500kb, 21 involved loci at which a nearby SNP also reached significance. For six of those associations, the top SNP association became non-significant upon conditioning on the CNV. For example, a low-frequency (MAF = 2.2%) deletion upstream of *BMP5*, which encodes bone morphogenetic protein 5, associated strongly with increased bone mineral density (0.12 (0.01) s.d.; P = 9.2 × 10^−82^) and appeared to explain strong SNP associations nearby (P = 3.8 × 10^−51^, conditional P = 0.24; Fig. 3c and Supplementary Table 10), highlighting the importance of including structural variants in GWAS fine-mapping. *BMP5* SNP and indel PTVs associated with stronger effects on bone mineral density (0.48 (0.17) s.d.; P = 0.005), suggesting that the deletion might affect an upstream regulatory region for *BMP5*, and motivating further exploration of allelic series including CNVs and SNPs.

### Allelic series involving both regulatory and gene-altering CNVs

Several CNV-trait associations contributed to long allelic series involving both CNVs that appeared to modify regulatory elements as well as CNVs that directly affected genes, providing opportunities to explore the effects of such mutations relative to one another and to SNP and indel polymorphisms. At the *α*-globin locus, at which copy-number polymorphisms of *HBA2* and *HBA1* (both encoding *α*-globin) are known to cause thalassemias, an extended allelic series containing eight classes of CNVs enabled further insights into genetic control of alpha-globin expression (Fig. 4a, Supplementary Fig. 5, and Supplementary Table 11). *α*-globin and β-globin together compose hemoglobin, and both the production and balance of *α*- and *β*-globin are important for normal erythropoiesis (such that relatively too little *α*-globin can lead to *α*-thalassemia whereas *α*-globin duplication can increase the severity of *β*-thalassemia)^30,31^. In UK Biobank, ultra-rare deletions that spanned either the *α*-globin gene pair, the upstream *α*-globin locus control region (HS-40), or the entire *α*-globin locus all associated with strongly decreased (∼ 3 s.d.) mean corpuscular hemoglobin (MCH) and increased red blood cell (RBC) counts, consistent with such mutations causing *α*-thalassemia by inactivating the locus^30,32–35^. “Silent” deletions of only *HBA2* associated with a relatively milder 1.7 (0.2) s.d. decrease in MCH. Intriguingly, duplications of these genomic elements exhibited a further range of effects: while duplications that increased *α*-globin gene dosage by 1-2 copies appeared to have little or no impact on MCH, duplications of the entire *α*-globin locus appeared to have an effect similar to loss of one *α*-globin gene (1.9 (0.2) s.d. lower MCH). This allelic series suggests that increased and decreased *α*-globin expression result in similar hematological phenotypes (consistent with the importance of balance in *α*- and *β*-globin) and that enhancer function rather than *α*-globin gene dosage primarily limits increases in *α*-globin expression. These results illustrate the ability of biobank-scale CNV analyses to extend knowledge even at well-studied loci.

**Figure 4:**
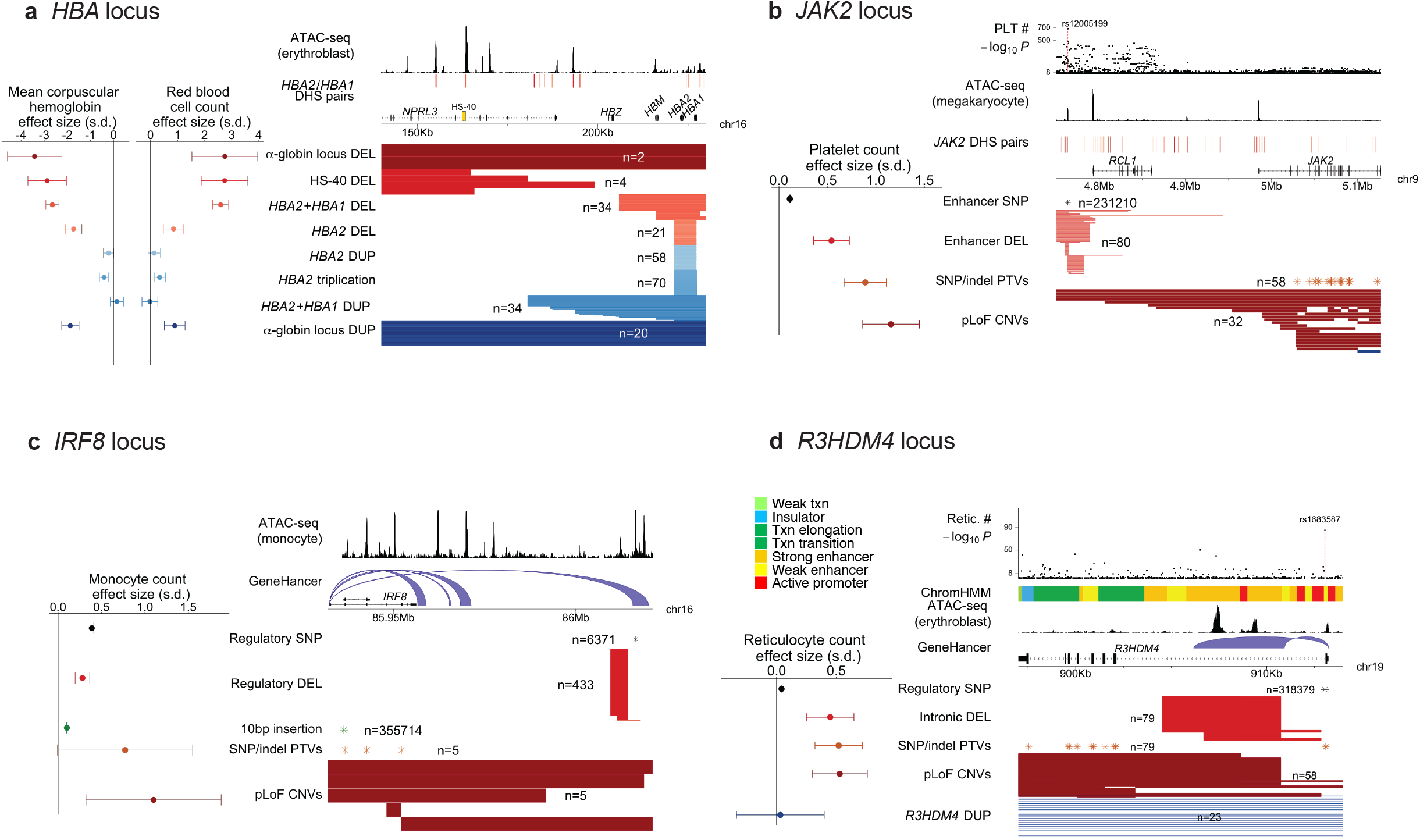
Allelic series involving both regulatory and gene-altering CNVs. **a. *HBA* locus**. Eight classes of CNVs at the *α*-globin locus and their effect sizes for mean corpuscular hemoglobin and red blood cell counts. Genomic annotations indicate accessible chromatin regions in erythroblasts^37^ and distal DNase I hypersensitive sites (DHS) for *HBA2/HBA1*^50^, highlighting the HS-40 super-enhancer. **b. *JAK2* locus**. Four classes of variants – *JAK2* pLoF CNVs, *JAK2* SNP and indel PTVs, a deletion of a distal enhancer, and the common SNP rs12005199 within the enhancer – and their effect sizes for platelet counts. Genomic annotations indicate accessible chromatin regions in megakaryocytes^37^ and *JAK2* distal DHS pairs^50^, which colocalize with common-SNP platelet count associations (top) at the enhancer region ∼220kb upstream of *JAK2*. **c. *IRF8* locus**. Fine-mapped common variants and rare pLoF variants at the *IRF8* locus – including a putatively regulatory distal deletion, *IRF8* pLoF CNVs, and *IRF8* SNP and indel PTVs – and their effect sizes for monocyte counts. Genomic annotations indicate accessible chromatin regions in monocytes^37^ and GeneHancer connections^42^ between downstream regulatory regions and *IRF8*. **d. *R3HDM4* locus**. Rare CNVs, SNP and indel PTVs, and a common intronic SNP at *R3HDM4* and their effect sizes for reticulocyte counts. Genomic annotations indicate ChromHMM^41^ annotations, accessible chromatin regions in erythroblasts^37^, and GeneHancer connections^42^, all indicating regulatory function in the first intron of *R3HDM4*. The lead-associated SNP rs1683587 (top) also lies within this intron, suggesting regulatory function. In **a** and **b**, DHS pairs are colored by their correlation value, from light red (correlation < 0.8) to dark red (correlation >0.95). Error bars on effect sizes, 95% CIs. Numerical results are available in Supplementary Tables 11-15.

Some allelic series involved known gene-trait relationships but appeared to reveal novel CNV effects with no SNP analogues. At *JAK2*, ultra-rare CNVs predicted to cause loss of JAK2 function associated with a 1.16 (0.15) s.d. increase in platelet counts (P = 9.9 × 10^−15^; Fig. 4b and Supplementary Table 12). This association, which replicated in an analysis of SNP and indel PTVs (*β* = 0.89 (0.11) s.d., P = 1.1 × 10^−15^; Fig. 4b), corroborated previous reports of an unexpected negative regulatory role for Jak2 in thrombopoiesis^36^. Interestingly, a distinct set of ultra-rare deletions centered ∼220kb upstream of *JAK2* associated with a 0.54 (0.09) s.d. increase in platelet counts (P = 9.5 × 10^−9^; Fig. 4b and Supplementary Table 12), roughly half the effect size of pLoF variants. The focal <4kb region shared by these deletions matched a strong megakaryocyte-specific accessible chromatin region previously implicated by common-SNP association and fine-mapping studies^37^ (Fig. 4b) that appeared likely to regulate *JAK2* (Supplementary Table 13). However, deletion of the entire enhancer element associated with a five-fold larger effect on platelet counts than the single-base pair modifications produced by SNPs within the enhancer (Fig. 4b and Supplementary Table 12), highlighting the ability of CNVs to enable further insights into complex trait genetics by altering the genome in ways that SNPs cannot.

Copy-number variants also contributed to an extended allelic series at *IRF8*, which encodes a transcription factor critical to monocyte differentiation^38^. Strong SNP associations with monocyte counts have previously been observed at the *IRF8* locus, led by a common noncoding 10bp insertion in *IRF8* with a mild effect size (0.102 (0.002) s.d.; P = 7.8 × 10^−587^; Fig. 4c and Supplementary Table 14). Multiple SNPs downstream of *IRF8* also associated independently with monocyte counts (consistent with the presence of multiple distal enhancers^39,40^), including a low-frequency SNP (rs11642657; MAF=0.8%) with a larger effect size (0.39 (0.01) s.d.; Fig. 4c and Supplementary Table 14). CNVs provided further insights into complex genetics at this locus: loss of one functional copy of *IRF8* (identified in 10 carriers of either pLoF CNVs or PTVs) appeared to produce a larger increase in monocyte count (0.94 (0.28) s.d.; P = 0.0009), while a downstream deletion near rs11642657 had a moderate effect size similar to this SNP (0.28 (0.04) s.d.; P = 4.7 × 10^−11^), suggesting the presence of an important regulatory region (Fig. 4c).

Some allelic series implicated new gene-trait associations. Ultra-rare deletions at *R3HDM4*, a gene with unknown function, associated with 0.54 (0.08) s.d. higher reticulocyte counts (P = 3.5 × 10^−11^; Fig. 4d and Supplementary Data 2). This association was corroborated by *R3HDM4* PTVs (*β* = 0.52 (0.10) s.d., P = 2.7 × 10^−7^), and a common intronic SNP also exhibited a mild-effect but strongly significant association with reticulocyte counts (*β* = 0.041 (0.002) s.d., P = 6.6 × 10^−86^; Fig. 4d and Supplementary Table 15). Interestingly, closer inspection of the deletions showed that they consisted of both exon-overlapping, pLoF deletions as well as intronic deletions falling fully within the first intron of *R3HDM4*, yet associating with a similar increase in reticulocyte counts (0.45 (0.10) s.d.; Fig. 4d). These results suggest a key regulatory role of the intronic region spanned by the deletions, which contains an accessible chromatin region (in erythroblasts) with predicted *R3HDM4* enhancer function^41,42^. Despite their associations with reticulocyte counts, neither type of deletion appeared to affect red blood cell counts (P = 0.17). These observations, which will require further understanding of R3HDM4 function to explain, again show the ability of regulatory CNVs to have significant phenotypic impacts, sometimes as strong as gene-dosage altering CNVs.

### Diverse potential functional impacts of CNVs

The remaining likely-causal CNVs involved in new gene-trait associations (Fig. 2e) appeared to alter gene dosage or function via a diversity of genomic modifications. Four rare deletions appeared to reduce or abolish gene function in a variety of ways. Two deletions associated with height: an inframe deletion of *DIS3L2* exon 9 previously reported to reduce ribonuclease activity and cause Perlman syndrome (an autosomal recessive disease characterized by congenital overgrowth)^43^ surprisingly appeared to *decrease* height by 0.44 (0.04) s.d. in heterozygous carriers (P = 3.9 × 10^−22^), and a whole-gene deletion of *SLC35E2B* associated with modestly decreased height and increased MCH (Supplementary Data 2). Two other deletions associated with ∼0.2-0.3 s.d. effects on platelet traits: an inframe deletion of *DOK3* exon 3 and a deletion of the final exon of *PARVB* (encoding 26 of 364 amino acids) (Supplementary Data 2).

Another novel gene-trait association involved ultra-rare (MAF=0.003%), large (>700 kb) duplications that appeared to target a single gene, *CXCR4*, and associated with a 0.99 (0.17) s.d. decrease in monocyte counts (P = 5.5 × 10^−9^, Supplementary Data 2). Gain-of-function mutations within *CXCR4* (chemokine receptor 4) cause autosomal dominant WHIM syndrome, an immunodeficiency disease^44^. Here, duplication of *CXCR4* appeared to produce relatively milder decreases in leukocyte counts (including 0.5 (0.2) s.d. reduced neutrophil and lymphocyte counts) with no apparent disease phenotypes.

A final association with platelet distribution width involved a low-frequency (MAF=0.7%) variant that initially appeared to be a duplication at *MTMR2* (Supplementary Data 2) but was surprisingly absent from CNV reference data sets^2,45^. Closer examination of sequencing reads from exome-sequenced carriers revealed that the structural variant actually constitutes a retroposition of the spliced *MTMR2* transcript into an intron of *LRCH1* (Supplementary Note). A common SNP haplotype in a different intron of *LRCH1* strongly and independently associated with increased platelet distribution width (P = 2.5 × 10^−172^), and both the SNP association and the insertion variant association (P = 3.5 × 10^−17^) appeared to be mediated by reduced *LRCH1* expression (based on analyses of GTEx data^46^; Supplementary Note), with the insertion exhibiting four-fold larger effects (Supplementary Fig. 6 and Supplementary Table 16). This unexpected finding from SNP-array analysis hints at further discoveries that will be enabled by sequencing technologies capable of comprehensively genotyping structural variants.

### Contrasting effects of deletions and duplications

Total genomic deletion burden and duplication burden have each been shown to associate with deleterious effects on several human traits^11,47,48^. We similarly observed negative associations of deletion and duplication burden with height and years of education (even after excluding syndromic CNVs), with deletions appearing to be roughly four-fold as deleterious as duplications (Fig. 5a,b and Supplementary Table 17). The consistent negative effect directions of deletion burden and duplication burden contrasted with the opposite effect directions that we observed at several loci involving focal reciprocal CNVs (Supplementary Data 2).

**Figure 5:**
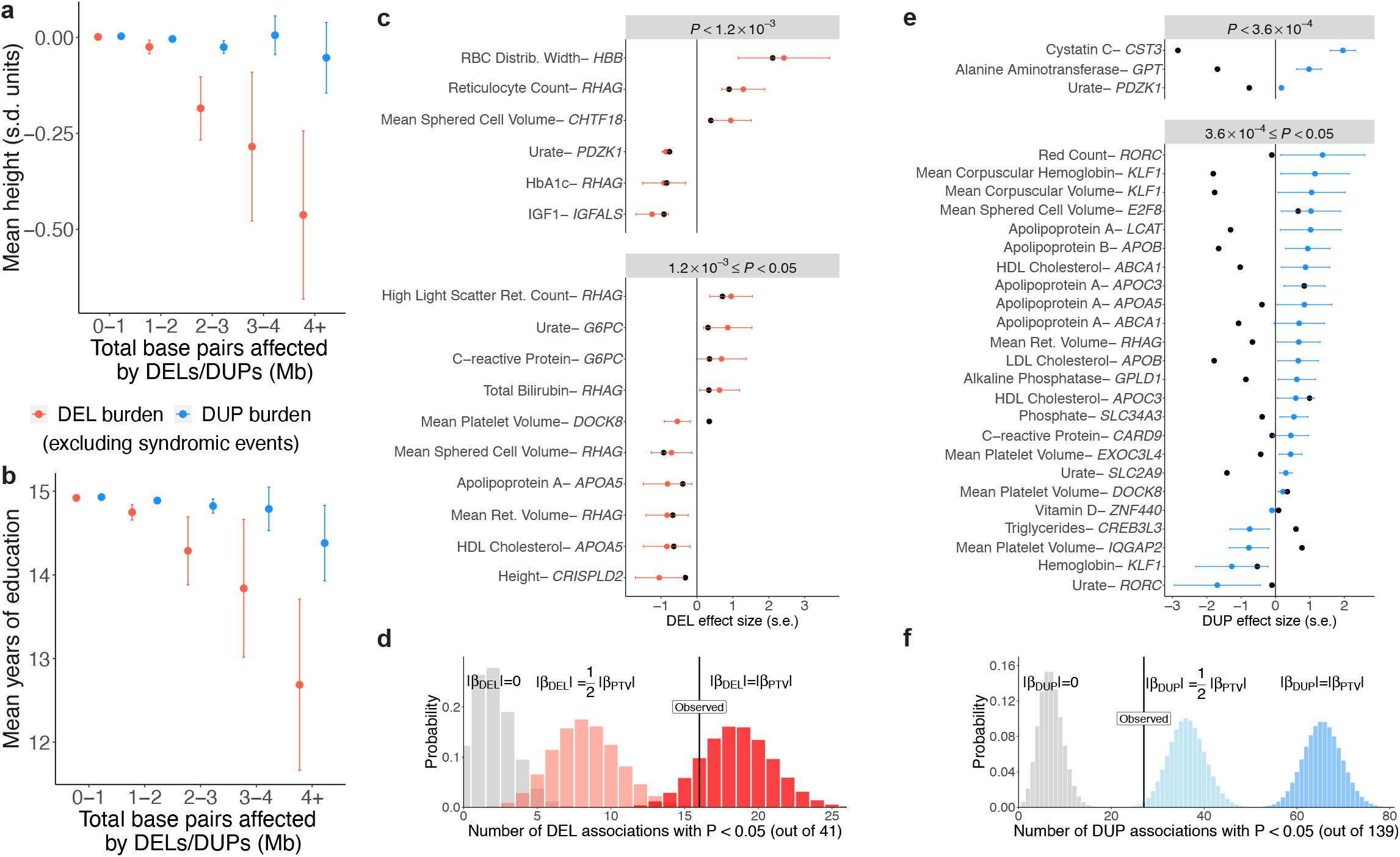
Contrasting phenotypic effects of deletions and duplications. **a**,**b**. Mean height (**a**) and years of education (**b**) as a function of total genomic length affected by deletions and duplications. Individuals carrying a known syndromic CNV were excluded from analysis. Numerical results are presented in Supplementary Table 17. **c**. Associations between whole-gene deletions and quantitative traits in targeted analyses of 41 gene-trait pairs for which we previously identified likely trait-altering PTVs^23^ and for which the HI-CNV call set contained at least two whole-gene deletions. Effect sizes and 95% confidence intervals are shown in red for 16 genes for which whole-gene deletions exhibited nominally significant associations (P < 0.05); effect sizes for SNP or indel PTVs^23^ are shown in black. **d**. Observing 16 nominally significant associations was consistent with whole-gene deletions having the same effects as PTVs. Probability distributions indicate numbers of significant associations in simulations in which whole-gene deletions have no effect (grey), half the effect magnitude as PTVs (light pink), or the same effect magnitude as PTVs (red). **e**,**f**. Analogous results for whole-gene duplications in targeted analyses of 139 gene-trait pairs, which produced 27 significant associations (P < 0.05), consistent with whole-gene duplications having less than half the effect magnitude of PTVs. (The aberrant effect directions of *DOCK8* deletions and duplications relative to the *DOCK8* PTV rs192864327 may be explained by this variant only causing loss of function in one of several transcripts.)

To more thoroughly explore the relative effects of focal deletions and duplications, we examined gene-trait pairs for which we had previously identified PTVs likely to alter quantitative traits^23^. For each gene, we compared the effects of likely-causal PTVs to those of whole-gene deletions and duplications (Supplementary Note). As expected, gene deletions acted similarly to PTVs, with 16 of 41 genes exhibiting nominally significant deletion associations (Fig. 5c), consistent with available power (Fig. 5d). In contrast, gene duplications tended to act in the opposite direction as PTVs and with smaller effect magnitudes: 27 of 139 genes exhibited nominally significant duplication associations (Fig. 5e), consistent with duplications tending to have less than half the effects of deletions (Fig. 5f, Supplementary Fig. 7, and Supplementary Table 18). These results suggest a contrast between CNV burden, which may be driven by large CNVs that disrupt many genes and tend to be deleterious regardless of deletion or duplication status, versus focal CNVs, which may tend to change the dosage of a specific key gene, resulting in reciprocal effects of deletions and duplications.

## Discussion

These results demonstrate the power of haplotype-informed structural variant analysis that leverages pervasive distant relatedness within large biobank cohorts to pool information about variants co-inherited by individuals who share extended SNP haplotypes. Applied to explore CNV-phenotype associations in UK Biobank, this approach uncovered many new ways in which genetic variation influences complex traits. At several loci, large-effect CNVs newly implicated putative target genes, and at several other loci, CNVs, together with nearby SNPs, created long allelic series illustrating the ability to CNVs to produce functional effects with no SNP analogues (e.g., gene copy-gain and regulatory element deletion or duplication).

Beyond the specific biological findings reported here, our study also provides a careful analytical approach for handling the statistical subtleties of performing association and fine-mapping analyses on difficult-to-call structural variants that can span large genomic regions. Additionally, the observation of several CNVs that represented lead associations at loci underscores the importance of considering structural variation even when performing statistical fine-mapping of SNP associations^15,49^.

These results also motivate further exploration of the far-larger set of CNVs that were not accessible to our analyses. While our approach enabled detection of 6-fold more CNVs than previous analyses of UK Biobank, and these CNVs appeared to account for roughly half of the rare LoFs estimated to arise from structural variation^2^, the CNVs we detected from SNP-array data still represent only a small fraction of the thousands of CNVs typically present in each human genome^2,3^. In particular, we were unable to ascertain CNVs smaller than the resolution of the SNP array (Supplementary Table 4), and we were also unable to genotype most common CNVs (MAF > 5%) due to inadequate SNP-array coverage and breakdown of modeling assumptions (Supplementary Table 5). These limitations could be overcome by extending the HI-CNV framework to whole-exome or whole-genome sequencing data, which is a promising direction for future research, especially at loci that are challenging to genotype. We anticipate that future studies using these and other approaches will provide further insights into the phenotypic consequences of copy-number variation.

## Acknowledgements

We thank R. Handsaker and P. Palamara for helpful discussions and S. Stankovic for providing PTV annotations on UK Biobank exome-sequencing variants. This research was conducted using the UK Biobank Resource under application number 40709. M.L.A.H. was supported by US NIH Fellowship F32 HL160061. M.A.S. was supported by the MIT John W. Jarve (1978) Seed Fund for Science Innovation and US NIH Fellowship F31 MH124393. A.R.B. was supported by US NIH grant T32 HG229516 and fellowship F31 HL154537. R.E.M. was supported by US NIH grant K25 HL150334. V.G.S. received support from NIH grants R01 DK103794 and R01 HL146500, as well as the New York Stem Cell Foundation. P.-R.L. was supported by US NIH grant DP2 ES030554, a Burroughs Wellcome Fund Career Award at the Scientific Interfaces, the Next Generation Fund at the Broad Institute of MIT and Harvard, and a Sloan Research Fellowship. The funders had no role in study design, data collection and analysis, decision to publish or preparation of the manuscript. Computational analyses were performed on the O2 High Performance Compute Cluster, supported by the Research Computing Group, at Harvard Medical School (http://rc.hms.harvard.edu).

## Methods

### UK Biobank genetic and phenotypic data

Genome-wide SNP-array data, including allelic dosages of pairs of alleles (labeled A and B) for 805,426 biallelic variants, was previously generated for 488,377 UK Biobank participants^18^. For CNV-calling, these allelic intensities are typically transformed to measures of total intensity (LRR) and relative intensity (B-allele frequency, BAF). We analyzed the LRR values provided by UK Biobank after first applying two de-noising steps: (i) GC-correction of total allelic intensities and (ii) principal component (PC)-correction of LRR^51^; and we directly computed relative genotyping intensities (Supplementary Note). We also analyzed whole genome sequencing (WGS) data available for 48 individuals (for validation analyses) and whole exome sequencing (WES) data available for 200,643 individuals^27^ (for follow-up analyses).

We restricted primary analyses to individuals of self-reported European ancestry included in the UK Biobank imputed dataset^18^, and we excluded individuals with trisomy 21, blood cancer, or those who had withdrawn at the time of our study (Supplementary Note), resulting in 454,759 participants with array data, 43 individuals with WGS data, and 186,105 individuals with WES data.

We analyzed 56 heritable quantitative traits measured on the majority of UK Biobank participants. These traits included anthropometric traits, blood pressure, measures of lung function, bone mineral density, blood cell indices, and serum biomarkers (Supplementary Data 1). Quality control and normalization of the quantitative traits was previously described^22,23^.

### Overview of HI-CNV method for haplotype-informed CNV detection

We reasoned that CNV detection sensitivity from SNP-array data available in UK Biobank could be considerably increased via two orthogonal strategies: (a) estimating SNP-specific priors for allele combinations corresponding to CNV states (to enable more accurate assessment of probabilistic information about copy-number variation provided by probe intensities); and (b) integrating probe intensity data across individuals likely to have co-inherited a large genomic tract. To estimate SNP-specific priors for allele combinations corresponding to CNV states, we (i) directly estimated SNP-specific genotype cluster priors at a subset of SNPs covered by large, easily-called CNVs; and then (ii) used these SNPs as a reference set from which SNP-specific priors for other SNPs could be predicted (based on which SNPs in the reference set exhibited most-similar probe intensity patterns). To incorporate probe intensity data across individuals likely to have co-inherited a large genomic tract, for each individual and genomic position on the SNP-array, we used a PBWT-based algorithm to find the 10 longest identical-by-descent (IBD) matches (per haplotype of the individual) spanning the position under consideration.

We used a hidden Markov model to call CNVs, integrating probabilistic information about copy-number state across an individual and their “haplotype neighbors” by weighting each neighbor’s information according to length of IBD-sharing. In more detail, at each SNP, for the individual and for each haplotype neighbor, we computed Bayes factors for deletion and duplication states based on genotyping intensities from the corresponding sample. We then created a weighted sum of log Bayes factors at each SNP, giving higher weights to haplotype neighbors with longer IBD. We ran this analysis using several different weighting schemes (trading off sensitivity to more recent vs. older mutations) and compiled calls made across these weighting schemes.

We filtered CNV calls to deletions larger than 75bp and duplications larger than 500bp and removed individuals with more than 100 CNV calls. Many of the samples with aberrantly many CNV calls appeared to share rare technical artifacts in LRR that had escaped denoising. We therefore computed the first 10 principal components of LRR in these aberrant individuals, ranked all individuals by the amount of LRR variance explained by these artifact PCs, and further removed individuals in the top 0.5%. Finally, for all downstream analyses, we removed calls on any chromosome in which we had previously detected a mosaic CNV^52^ as well as calls in regions with frequent somatic events. After these quality control filters, we had called CNVs in 452,500 UK Biobank participants (including 43 individuals with WGS data and 185,365 individuals with WES data). Further methodological details are available in the Supplementary Note.

### PennCNV call set in UK Biobank

We compared HI-CNV calls to previously-generated PennCNV^7^ calls made by analyzing Affymetrix CEL files (Return 1701)^10^. Following suggested quality control procedures^8^, we filtered individuals with more than 30 calls made, a genotype call rate less than 96%, or an absolute waviness factor greater than 0.3. To facilitate comparison to our HI-CNV call set, we then applied the same additional filtering of calls on chromosomes containing mosaic CNVs and in regions with frequent somatic events.

### Precision and recall of HI-CNV and PennCNV call sets

To benchmark performance of HI-CNV and PennCNV, we analyzed independent WGS data available for 43 individuals using CNVnator^53^ and DELLY^54^. To assess the precision, or validation rate, of array-based calls we computed the proportion of HI-CNV (respectively, PennCNV) calls that were either (1) replicated by CNVnator calls or (2) exhibited enrichment or depletion of read-depth (computed by CNVnator) consistent with the CNV call. To assess recall, or sensitivity, of HI-CNV and PennCNV, we analyzed calls from DELLY, which produced a merged call set across WGS samples that was helpful for computing recall of CNVs within allele frequency ranges. For each DELLY call, we annotated whether HI-CNV (respectively, PennCNV) called an overlapping event. Further details on computing precision and recall are provided in the Supplementary Note.

### Stratifying carrier counts of gene dosage-modifying CNVs by LOEUF score

For each protein-coding gene, we computed the number of UK Biobank participants of European ancestry carrying whole-gene deletions, whole-gene duplications, and CNVs predicted to cause loss of function (pLoF; Supplementary Note). We then annotated each gene with its LOEUF sextile bin (‘oe_lof_upper_bin_6’ from the pLoF Metrics by Gene TSV file downloaded from https://gnomad.broadinstitute.org/downloads), which estimates strength of selection against protein-truncating mutations^55^. We restricted to genes with a non-missing LOEUF sextile bin and genes with only one annotated canonical transcript. In Fig. 1g, we reversed the order of LOEUF sextile bins such that higher-numbered bins correspond to more-constrained genes.

### Association testing and statistical fine-mapping

We performed CNV-phenotype association analyses on three distinct classes of CNVs defined based on 1) SNP-array probe overlap, 2) gene overlap, and 3) specific CNVs. Analyses on the SNP probe level tested the hypothesis that a change in copy number (deletion or duplication, respectively) at the genomic location of the SNP alters the phenotype. Analyses on the gene level tested the hypothesis that a change in copy number affecting the gene in question (whole-gene deletion, whole-gene duplication, and pLoF, respectively) alters the phenotype. Analyses on the CNV level tested whether a specific CNV (allowing for slightly differing endpoints in calls from different samples) alters the phenotype. These tests comprised both burden-style analyses (the probe- and gene-level tests) and single-variant analyses (the CNV-level tests), for a total of ∼ 1.7 million tests. Given that these tests contained a high degree of redundancy (e.g., because probe-level tests at consecutive SNPs tended to be very strongly correlated), we used the standard genome-wide significance threshold (P < 5 × 10^−8^) to determine significant associations.

We conducted association tests using BOLT-LMM^21,22^ (--lmmForceNonInf) with assessment center, genotyping array, sex, age, age squared and 20 genetic principal components included as covariates. We fit the mixed model on directly genotyped autosomal variants with MAF > 10^−4^ and missingness < 0.1 and computed association test statistics for CNVs in the three categories defined above; a similar pipeline produced association test statistics for SNP and indel variants imputed by UK Biobank (the imp_v3 release) and variants we previously imputed from the first tranche of exome-sequencing of 49,960 participants^23^. We included all participants with non-missing phenotypes in the European-ancestry HI-CNV call set described above.

To filter significant associations to a set of likely-causal associations, we used a pipeline we previously developed^23^ to eliminate associations that could be explained by linkage disequilibrium (LD) with nearby variants (here, either SNP or indel variants from the UK Biobank imp_v3 release or variants we had imputed from WES^23^). This filter required CNVs to remain significant after conditioning on any other more strongly associated variant nearby. More explicitly, for every CNV *i* significantly associated with a given phenotype, we calculated its correlation *r*_*ij*_ with each more strongly associated variant *j* (including other CNVs and imputed SNPs and indels) within 3Mb using plink ‘--r’^56^. We then computed the approximate chi-square association statistic for CNV *i* conditioned on variant *j* as:

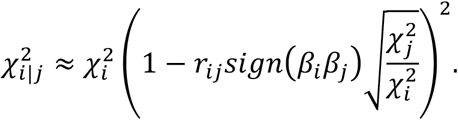

We defined likely-causal associations as those with the property that 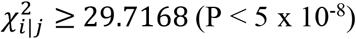 for all variants *j* more strongly associated with the trait than CNV *i*. We previously observed that this pairwise LD-based filter was effective for fine-mapping rare variant associations^23^.

### Defining and annotating CNV loci

To group phenotype-associated CNVs into genomic loci, we first identified a set of unique CNVs contributing to likely-causal associations (accounting for uncertainty in CNV breakpoints and for probe-level and gene-level tests aggregating signal across multiple CNVs; Supplementary Note). We then ordered this set of likely-causal CNVs from smallest to largest, and if a CNV fell within 100kb of a previous CNV, we considered it to be part of the same locus. We annotated a likely-causal CNV as syndromic if it overlapped a previously-curated pathogenic CNV^10^ by more than 50%. We identified putative target genes of non-syndromic, likely-causal CNVs either by observing that a focal CNV association only overlapped a single gene or by finding independent supporting evidence for a particular gene within or near the CNV region (specifically, a coding variant association or experimental literature). Further details on defining and annotating loci are provided in the Supplementary Note.

### Follow-up analyses at highlighted loci

At a subset of loci we investigated in greater detail (Fig. 3 and Fig. 4), we identified carriers of high-confidence loss-of-function SNP and indel variants (annotated using LOFTEE^55^) among the 185,365 individuals with whole-exome sequencing data^27^ in our analysis set. To increase power to assess phenotypic impacts of SNP and indel PTVs, we residualized phenotypes for polygenic predictions of the phenotype using array-typed SNPs (omitting those within 2Mb of the gene of interest) that we generated using BOLT-LMM ‘—predBetasFile’ in 10-fold cross-validation (emulating linear mixed model association analysis)^57^. Residualized phenotypes could then be modeled as a function of SNP and indel PTV carrier status, as well as carrier status for other CNVs or SNPs of interest. We performed these analyses after our initial association analyses, such that numbers of carriers of CNVs differ slightly between Supplementary Data 2 and the locus plots in Fig. 3 and Fig. 4 (generated using karyoploteR^58^) due to participant withdrawals.

## Data availability

Access to the UK Biobank Resource is available by application (http://www.ukbiobank.ac.uk/). Individual-level HI-CNV calls and summary association statistics for the 56 quantitative traits we analyzed will be returned to UK Biobank.

## Code availability

The following publicly available software packages were used to perform analyses: BOLT-LMM (v2.3.5), https://data.broadinstitute.org/alkesgroup/BOLT-LMM/; plink (v1.9), https://www.cog-genomics.org/plink/1.9/; CNVnator, https://github.com/abyzovlab/CNVnator; DELLY, https://github.com/dellytools/delly. Code and scripts used to perform CNV-calling and downstream analyses will be released prior to publication.

