## Supplementary Information for "Influences of rare copy number variation on human complex traits"

**Supplementary Figures 30**

**Supplementary Tables 43**

**References 63**

### Supplementary Notes

#### 1 Transforming and denoising SNP-array genotyping intensities

UK Biobank provided genotyping intensity data generated by Affymetrix in two formats:

1. `int` files containing intensity values for the A and B alleles of each genotyped variant
2. `baf` and `l2r` files containing B allele frequency (BAF) and $\log_2$ R ratio (LRR) transformed intensity values (measuring relative and total genotyping intensities across the two alleles) used by typical CNV-calling pipelines.

Affymetrix's genotype-calling algorithm modeled relative and total genotyping intensities by estimating bivariate normal distributions corresponding to "SNP clusters" for the three possible diploid (copy number 2; CN=2) genotypes (AA, AB, BB). We wished to extend this genotyping framework by additionally estimating bivariate normal SNP clusters for each possible genotype cluster corresponding to heterozygous CNVs, i.e., deletions (CN=1: A, B) and duplications (CN=3; AAA, AAB, ABB, BBB) (Supplementary Fig. 8).

To do so, we required relative and total genotyping intensity measurements that were reasonably well-modeled by normal distributions. For relative genotyping intensities, the BAF values provided by UK Biobank did not meet this criterion because they had been truncated to fall between 0 and 1 (such that many individuals with homozygous genotypes had BAF of either 0 or 1). We therefore computed relative genotyping intensities from the `int` data for the A and B alleles by applying a polar-like transformation [1]:

$$\theta = \frac{2}{\pi} \cdot \arctan\left(\frac{B}{A}\right) \tag{1}$$

For total genotyping intensities, we analyzed the LRR (`l2r`) values provided by UK Biobank after first applying two denoising steps described below.

##### 1.1 GC-correction of total allelic intensities (LRR)

We first corrected LRR values for "GC waves" [2] using a simplified version of a previously-described pipeline [3, 4]. Specifically, for each sample, we regressed LRR on proportions of GC and CpG content in 9 windows (spanning 50, 100, 500, 1k, 10k, 50k, 100k, 250k, and 1M bp) centered around each variant. We computed GC content using bedtools [5] on the human reference (hg19), and we computed CpG content using the EpiGRAPH CpG annotation [6].

##### 1.2 Principal component (PC)-correction of LRR

Even after GC-correction, top principal components of the LRR matrix explained large fractions of variance, indicating that the LRR data could be further-denoised by projecting out top PCs capturing unmodeled technical noise [7]. We took two precautions to guard against top PCs inadvertently capturing real signal from common CNVs:

1. We computed principal components on genome-wide LRR values for all autosomal variants at once (separately for each genotyping batch), reasoning that technical artifacts should behave similarly genome-wide (whereas inter-sample correlations in LRR driven by copy number variation would be locus-specific)—such that genome-wide PCs are more likely to pick up technical artifacts and less likely to "overfit" to local features.

2. We computed LRR PCs using only white British samples in order to reduce the potential for PCs to capture ancestry effects. (We then projected top PCs out of all samples in the genotyping batch).

We applied the above PC-correction procedure independently to each of the 106 genotyping batches, projecting the top 50 PCs out of LRR for each batch. We observed that these top 50 PCs explained an average of 58.5% of LRR variance. Additional PCs provided little marginal increase in variance explained (e.g., 100 PCs explained 61.6% of variance on average across batches; Supplementary Fig. 9 and Supplementary Table 19).

#### 2 Estimating genotype cluster parameters

SNP-array genotyping platforms use allele-specific oligonucleotide probes to quantify the abundance of each of two alleles (A and B) in a DNA sample. Genotyping of biallelic variants in regions of the genome that do not vary in copy number can then be performed by clustering measured probe intensities (across a batch of samples) into clusters corresponding to the three possible diploid genotypes (AA, AB, BB). Such clustering is usually performed using SNP-specific priors on the expected distribution of bivariate probe intensities assuming each possible genotype (AA, AB, BB), which for technical reasons can vary substantially among SNPs. Genotyping in this manner produced highly accurate genotype calls in UK Biobank ($\sim$99.9% accuracy with $<$1% missingness at most SNPs) [8].

SNP-array probe intensities are also informative of copy-number variants that overlap SNPs on an array, resulting in measured intensities that deviate from the clusters corresponding to the usual three diploid genotypes (AA, AB, BB) [9, 10]. Because these deviations are less dramatic than the differences in probe intensities that separate diploid genotypes, CNV-calling from the Affymetrix SNP-arrays used by UK Biobank (which produced relatively noisy probe intensity measurements) has tended to require combining signal across at least ∼10 SNPs, resulting in detection of only an average of ∼4–6 CNVs per sample [11, 12].

We reasoned that CNV detection sensitivity from SNP-array data available in UK Biobank could be considerably increased via two orthogonal strategies: (a) estimating SNP-specific priors for allele combinations corresponding to CNV states (Supplementary Fig. 8), thereby enabling more accurate assessment of probabilistic information about copy-number variation provided by probe intensities; and (b) incorporating probe intensity data from individuals likely to have co-inherited a large genomic tract. In this section we describe strategy (a), which was previously employed by the Birdsuite software [13]; here, we were able to leverage the very large size of the UK Biobank cohort to learn information about SNP-specific priors from the data, requiring less extrapolation. The basic idea of our approach was to (i) directly estimate SNP-specific genotype cluster priors at a subset of SNPs covered by large, easily-called CNVs; and then (ii) use these SNPs as a reference set from which SNP-specific priors for other SNPs could be predicted (based on which SNPs in the reference set exhibited most-similar probe intensity patterns).

#### 2.1 Partitioning samples into LRR-noise deciles

We first estimated a per-sample parameter reflecting overall amount of technical noise in probe intensities, which varied among samples. We computed this per-sample parameter as the RMSE (in standardized units) of LRR across autosomal variants on the SNP-array. That is, for each genotyped variant, we standardized LRR to have mean 0 and variance 1 across samples, and then for each sample, we computed the sample's "noise scale factor" as the root-mean-square of standardized LRR across all autosomal variants.

We used these estimated noise scale factors to partition samples into noise deciles for downstream modeling of probe intensities, reasoning that the shapes and positions of probe intensity distributions might change somewhat depending on the amount of technical noise present in a sample. We also further adjusted for within-decile variation in noise scale factors when estimating Bayes factors for copy-number states given observed probe intensities (both in our initial LRR-based model and our final HI-CNV model; see the descriptions of these computations below for details).

#### 2.2 Generating reference data via LRR-based calling of large CNVs

To obtain examples of probe intensities corresponding to copy-loss and copy-gain genotypes (loss = {A, B}; gain = {AAA, AAB, ABB, BBB}), we implemented a simple hidden Markov model (HMM) that called loss and gain events in each sample independently using only LRR values together with heterozygous SNP calls (used as evidence against deletions). This approach was designed to efficiently generate a high-confidence callset of large CNVs, providing data about probe intensity distributions for SNPs within these CNVs.

Specifically, the HMM contained three copy-number states (CN = 1, 2, 3), with transition and emission parameters defined as follows:

- Transition penalties of $10^{-3}$ were assessed for jumping between adjacent states and $10^{-6}$ for jumping between CN=1 and CN=3.
- Emission probabilities were computed assuming that LRR was generated from a Gaussian distribution with:
  - Mean equal to 0 for CN=2; mean equal to the empirical mean LRR in large deletions and duplications (estimated by iteratively running this HMM algorithm) for CN=1 and CN=3, respectively.
  - Standard deviation estimated per-SNP as the empirical standard deviation of LRR across samples in a noise decile, scaled by the relative noise scale factor (relative to the median-noise sample in the decile) for the sample being analyzed.
- To limit the influence of outliers, relative emission probabilities for CN=1 vs. CN=2 and CN=3 vs. CN=2 were cropped to the range $[10^{-4}, 10^{4}]$.
- An additional (multiplicative) emission penalty was assessed for the CN=1 state if a SNP had been called as heterozygous (since all SNPs within a deletion should be hemizygous). This penalty factor ranged from $5 \times 10^{-6}$ (for the highest-confidence SNP calls) to 1 (for zero-confidence calls), scaling logarithmically according to call confidence values provided by Affymetrix.

We used the Viterbi algorithm to identify putative CNVs (as segments of CN=1 and CN=3 states in the most likely path through the HMM). We then created a stringent set of (sample, SNP) pairs very likely to provide examples of probe intensity measurements arising from copy-loss or copy-gain states by restricting to:

- SNPs well within deletion calls spanning 15+ SNPs ($>$3 SNPs from either end).
- SNPs well within duplication calls spanning 50+ SNPs ($>$10 SNPs from either end).

We required deletions and duplications to be large both to minimize false positives in our reference data and to avoid ascertainment bias (which could occur for shorter CNVs if calls were only made in carriers for which LRR was especially large due to measurement noise). We also stringently trimmed the ends of CNV calls to avoid uncertainty in breakpoints (which was larger for duplications than for deletions), prioritizing data quality over quantity because reference data was relatively abundant in UK Biobank. We note that because we did not attempt to model CN=0 or

CN=4+ states, the reference data set we generated included a small fraction of homozygous CNVs; however, most of the large CNVs that we considered in this analysis were rare, and we ensured that our subsequent estimation of cluster priors was robust to this issue.

#### 2.3 Estimating parameters for clusters with available reference data

After identifying high-confidence within-CNV SNPs, we next needed to assign probe intensities from such SNPs (transformed to the ($\theta = \frac{2}{\pi} \arctan \frac{B}{A}$, LRR) scale) to genotype clusters for CN=1 (A, B) and CN=3 (AAA, AAB, ABB, BBB). We did so by dividing the ($\theta$, LRR) plane into zones designed to typically contain most data points from each possible cluster. We defined these zones in a SNP-specific, noise-decile-specific manner based on the locations and orientations of CN=2 clusters (i.e., Affymetrix genotype calls of AA, AB, BB):

- For CN=1, we split the plane left/right at the $\theta$ value of the CN=2 het (AB) cluster center.
- For CN=3, we additionally split the above half-planes at lines passing through the points 2/3 of the way from the CN=2 het (AB) cluster center to each CN=2 hom (AA, BB) cluster center. We drew these lines parallel to regression lines indicating the relationship between LRR (treated as the independent variable) and $\theta$ (treated as the dependent variable) among points in the respective CN=2 hom clusters: e.g., we approximately separated AAA and AAB clusters by drawing a line "parallel to the AA cluster" located 2/3 of the way from the AB cluster to the AA cluster. (If one of the CN=2 hom clusters was very rare ($n$<25), we did not perform the additional split, assuming that the corresponding AAA or BBB cluster would have negigibly low frequency.)

After provisionally assigning within-CNV probe intensity data points to clusters according to the above zones, we next removed outliers farther from the median (in either coordinate, $\theta$ or LRR) than twice the interquartile range.

The above partitioning and outlier removal strategy worked well for most clusters, but visual inspection of the data showed that a sizable minority of provisional clusters still contained data points that should have been assigned to other clusters. We therefore applied a few post-processing filters to flag questionable-quality clusters for exclusion from our reference set:

- Exclude all CN=1 minor-allele clusters for SNPs with MAF<0.05. Some of these provisional clusters contained a nontrivial fraction of data points that actually corresponded to CN=0, so we just excluded all such clusters (as we had no shortage of reference data from more-robust CN=1 clusters).
- Exclude any CN=3 cluster that overlaps with a neighboring CN=3 cluster with higher frequency (i.e., more data points). The rationale for this filter was that higher-frequency clus-

ters tend to be only mildly affected by mis-assigned points that actually belong in lower-frequency clusters, but not vice versa. We defined "overlap" as follows:

   - For the two CN=3 heterozygous clusters (AAB, ABB), we required the $\theta$-distance between the center of the cluster and the center of each of each neighboring CN=3 cluster to be at least the sum of the cluster width and the neighboring cluster width: $2 \cdot (\text{s.d.}(\theta)_{\text{cluster}} + \text{s.d.}(\theta)_{\text{neighbor}})$.
   - For the two CN=3 homozygous clusters (AAA, BBB) (which tended to be more affected by this problem), we required separation from the neighboring (het) CN=3 cluster center to be at least $2.5 \cdot (\text{s.d.}(\theta)_{\text{cluster}} + \text{s.d.}(\theta)_{\text{neighbor}})$.

- Exclude any cluster with aberrantly large variance in either coordinate ($\theta$ or LRR): i.e., variance greater than 1.5 times the sum of variance (of the same coordinate) in each of the three CN=2 clusters (AA, AB, BB). This filter tended to catch remaining CN=1 clusters containing CN=0 data points.

- Exclude all clusters from ultra-rare SNPs ($n$<25 het calls among samples in the noise decile).

For each noise decile, for each SNP, for each of the possible genotypes corresponding to CN=1 (A, B) and CN=3 (AAA, AAB, ABB, BBB), we considered the genotype cluster to be a suitable reference cluster if it contained at least 10 data points (after outlier removal) and had not been excluded by any of the above filters. Approximately 1% of all clusters satisfied this criterion. For each such cluster, we then estimated its five bivariate normal parameters—mean($\theta$), mean(LRR), var($\theta$), var(LRR), and cov($\theta$, LRR)—from its data points.

Finally, we also estimated bivariate normal cluster parameters for CN=2 genotype clusters simply by assigning all samples with non-missing genotype calls from Affymetrix to the corresponding cluster (and then removing outliers with either coordinate ($\theta$ or LRR) farther from the median than three times the interquartile range). As above, we required at least 10 data points to be assigned to a cluster in order to proceed with estimation of bivariate normal parameters; otherwise we set the cluster to missing. We did not attempt to identify and exclude data points corresponding to CNVs from this analysis given that (i) the vast majority of variants included on the UK Biobank SNP array were (by design) not in regions of the genome that harbor common copy-number variation; and (ii) our focus was on identifying rare, potentially-deleterious CNVs.

#### 2.4 Predicting cluster parameters for all genotyped variants

Having determined approximate location and shape parameters for ~1% of all clusters using the above procedure, we then sought to use this information to predict bivariate normal parameters for the remaining ~99% of clusters (for which we had insufficient or questionable-quality data from overlapping large CNVs). For each such "missing" cluster, the basic idea of our approach was to find the 20 reference SNPs with CN=2 clusters most similar to CN=2 clusters of the SNP in question, and then predict the "missing" cluster based on the location and shape of the corresponding (non-missing) cluster in the 20 reference SNPs.

Explicitly, for each noise decile, for each SNP, for each side (left/right) of the cluster plot, for each CN=1 and CN=3 cluster on the side under consideration (i.e., A, AAA, AAB on the left side; B, ABB, BBB on the right side), we matched the SNP's CN=2 clusters on the side under consideration (i.e., AA, AB on the left side; AB, BB on the right side) to the corresponding CN=2 clusters of reference SNPs at which the cluster had been estimated (typically $\sim$10,000 SNPs). We used squared Hellinger distance as a metric for assessing agreement between corresponding CN=2 clusters, summing across the two CN=2 clusters on the side under consideration. For example, for the left side:

$$d(\mathrm{pred}, \mathrm{ref}) = H^2(\mathrm{AA}_{\mathrm{pred}}, \mathrm{AA}_{\mathrm{ref}}) + H^2(\mathrm{AB}_{\mathrm{pred}}, \mathrm{AB}_{\mathrm{ref}}), \quad (2)$$

where "pred" denotes the SNP with clusters being predicted, "ref" denotes a reference SNP, and Hellinger distances are computed between the bivariate normal distributions at the "pred" and "ref" SNPs for each of the two left-side CN=2 clusters (AA and AB). We computed Hellinger distances on bivariate normal distributions for CN=2 clusters that we estimated cohort-wide (instead of within noise deciles) to allow more-robust cluster-matching at rare SNPs, which had few data points in the het and hom-minor CN=2 clusters.

After ranking reference SNPs in this manner, we selected the top 20 reference SNPs that genotyped most similarly to the SNP whose cluster was being predicted. By design, such "ref" SNPs had CN=2 clusters that closely matched those of the "pred" SNP; however, this alignment was not perfect. To adjust for small offsets between "pred" SNP vs. "ref" SNP CN=2 cluster centers, we shifted each "ref" SNP's CN=1/CN=3 clusters by the estimated offset (measuring the offset at the CN=2 cluster closest to the cluster being predicted, in the noise decile under consideration). Finally, we predicted bivariate normal parameters for the missing cluster of the "pred" SNP by computing its mean and covariance assuming that it was an equal mixture of the 20 reference clusters.

This approach also allowed us to predict clusters for rare SNPs at which the hom-minor CN=2 cluster was missing (due to insufficient data points). For such SNPs, we predicted all clusters (CN = 1, 2, 3) on the missing (minor-allele) side using the same approach as above, but defining most-similar reference SNPs based on the opposite-side CN=2 clusters (hom-major and het).

We developed the above approach using cross-validation analyses (in which we attempted to predict held-out reference clusters using other reference SNPs), and visual inspection of predicted clusters corroborated good cross-validation performance as well as good containment of CNV data points in clusters for which coverage by large CNVs had been too low to estimate reference clusters (Supplementary Fig. 8).

### 3 Finding longest identical-by-descent (IBD) matches per haplotype

Beyond optimizing modeling of genotyping probe intensities, the main source of HI-CNV's improved detection sensitivity is its use of IBD-sharing across distantly related individuals to amplify weak signals of CNV presence. This approach is inspired by the idea of validating variant calls in related samples (e.g., trios) by checking for Mendelian inheritance, a paradigm that is frequently used to benchmark variant callers or increase confidence in difficult-to-call variants. HI-CNV leverages the fact that population-scale cohorts such as UK Biobank contain extensive distant relatedness (Supplementary Fig. 10 and Supplementary Table 20), such that any polymorphic variant present in at least a few individuals is likely to have been co-inherited on a long, readily-identifiable shared haplotype. In such scenarios, combining probabilistic information about CNV presence across individuals who share long IBD can dependably aid detection. This idea builds upon previous approaches that modeled linkage disequilibrium between CNVs and common SNPs by considering short ancestral haplotypes [14] and that performed SNP-haplotype-based refinement of CNV likelihoods [15, 16].

In this section, we describe the algorithm we implemented to efficiently identify top IBD matches within UK Biobank: specifically, for each haplotype of each individual, and for each genomic position on the SNP-array, we wished to find the longest 10 IBD matches spanning the position under consideration. While several methods based on the positional Burrows-Wheeler transform (PBWT) [17] have recently been developed for rapid IBD detection in large cohorts [18–20], these methods aim to find all IBD segments above a fixed length (e.g., 2 or 3 cM) shared by pairs of haplotypes in a cohort—which could either result in too much output for our purposes (at loci containing very many IBD matches) or too little output (for haplotypes with only smaller lengths of IBD-sharing). We therefore implemented a simple PBWT-based algorithm (using a seed-and-extend approach similar to hap-IBD [19]) tailored to the specific task of finding longest-IBD matches. (Note that here we will be loose about the definition of "IBD"; a short, $\sim$1-cM match might not arise from a recent-enough common ancestor to typically be considered "IBD" but might still be helpful for calling common CNVs contained within it.)

#### 3.1 Identifying seed matches using the positional Burrows-Wheeler transform (PBWT)

The first step of our approach was to identify a set of long identical-by-state (IBS) segments among pairs of phased SNP-haplotypes, serving as seeds for extension into (potentially longer) IBD segments. We performed this search using the PBWT, which produces, at each genotyped SNP, a lexicographic sort of haplotype suffixes (when operating right-to-left) from which longest-IBS matches starting at each position can readily be obtained as bands of consecutive sorted haplotype suffixes [17]. Explicitly, every 32 SNPs processed by the PBWT, we augmented our set of IBS seed segments by selecting, for each haplotype, a band of adjacent haplotypes corresponding to $K = 5$ (first algorithmic iteration; see below) or $K = 10$ (second algorithmic iteration) longest IBS-suffix matches spanning at least 128 SNPs. For any IBS-suffix match that extended a sub-IBS-suffix previously selected, we eliminated the redundant, previously-selected sub-IBS-suffix from the seed set.

#### 3.2 Extending IBD seeds

Most IBD segments do not consist of a single segment of perfect IBS (i.e., exact matching of a contiguous sequence of alleles along a pair of SNP-haplotypes); instead, IBD segments usually contain a sequence of IBS segments punctuated by mismatches (typically arising from genotyping errors or gene conversions). For each IBS seed identified by the PBWT-based algorithm above, we therefore attempted to extend the IBS segment into a longer IBD segment using an approach similar to hap-IBD [19]. (We did not attempt to model phase switch errors given that our phased haplotypes for UK Biobank had chromosome-scale accuracy [21].)

Explicitly, we attempted to extend each IBS seed to the left and right in an error-tolerant manner based on matching scores that we computed on blocks of 64 SNPs (using fast parallelization of bitwise operations):

$$\text{64-SNP match score} = 1 - 2 \times (\text{\# ``soft'' errors}) - 4 \times (\text{\# ``hard errors}), \tag{3}$$

where “soft” and “hard” errors were defined based on genotype call confidences (on a 0–1 scale) provided by Affymetrix and UK Biobank. Specifically:

- We ignored errors at SNPs for which either sample in the pair had a genotype confidence $< 0.5$.
- Otherwise, we considered a “soft” error to be a mismatch at a SNP for which at least one sample had a genotype confidence in the range 0.5–0.75.
- The remaining errors (involving SNPs confidently genotyped in both samples) were considered “hard” errors.

Under this scoring scheme, perfect matches of 64-SNP blocks incremented the score of a segment being extended by 1, while matches with non-ignored errors reduced the score by 1 or more (depending on the number and type of errors). Upon encountering a negatively-scored block, we required the total score to break even within the next 12 blocks; otherwise, we ended IBS seed extension at the first error encountered within the block. This approach effectively required that each "soft" error be counterbalanced by 127 matched SNPs and each "hard" error be counterbalanced by 255 matches.

#### 3.3 Filtering to longest IBD matches per position per haplotype

From the list of IBD segments identified above, we wished to efficiently identify, for each haplotype and at each SNP-array position, a list of the top-$K$ longest IBD segments spanning this position. To do so, we first post-processed the set of IBD segments by merging any duplicated or overlapping segments (involving the same pair of haplotypes). Then, for each haplotype, we identified top-$K$ longest IBD matches at each SNP-array position using the following algorithm:

- Sort all IBD matches involving the haplotype by start coordinate.
- Walk left to right across the chromosome, maintaining an "active set" of IBD matches spanning the current position, sorted in two ways: (i) by IBD length (longest to shortest); and (ii) by end coordinate (left to right). At each position:
  - Update the active set if:
    - Current position starts a new IBD match: add new match to active set.
    - Current position ends an IBD match in the active set: delete ended match.
  - Read off the top-$K$ longest matches spanning the current position from the active set.

#### 3.4 Correcting potential genotype errors

Our identification of top IBD matches for each haplotype at each genomic position provided an opportunity to correct some of the occasional SNP-allele mismatches that interrupted IBS within IBD tracts. Doing so could potentially improve the quality of IBS seeds identified by the PBWT, which is not robust to mismatches. We therefore implemented an "error-correction" strategy in which we used IBD information to identify haploid SNP-alleles that were inconsistent with haplotypes sharing longest IBD, and we subsequently ran a second iteration of the entire IBD-finding algorithm after modifying these SNP-alleles. We limited error-correction to SNP-alleles for which genotyping call confidence had been reported to be low ($< 0.5$) by Affymetrix.

In more detail, for each haplotype, for each SNP-allele corresponding to a low-confidence genotype call, we identified the longest five IBD matches spanning the SNP-allele, as described above. We then examined the corresponding SNP-allele in each of the 5 IBD-neighbor haplotypes for which the SNP in question was located $> 0.5$ cM from the edge of the IBD segment. If at least four IBD-neighbors satisfied this requirement and only at most one of them agreed with the SNP-allele in the original haplotype, we recorded a likely error.

After analyzing all haplotypes in the above manner, we flipped the (haploid) SNP-allele genotypes at all recorded likely errors. We also used the information about potential errors to perform quality control on SNPs: for any SNP with likely errors in 0.25% or more haplotypes, we ignored this SNP in the next iteration of IBD-finding.

### 4 Calling CNVs using intensity data across haplotype neighbors

The methods described in the previous sections provided the two key ingredients of the HI-CNV algorithm: (i) detailed, SNP-specific (and sample noise decile-specific) priors on probe intensities produced by different genotypes; and (ii) information about longest IBD matches for each haplotype at each genomic location. Here we describe the algorithm that we used to convert probe intensity data into probabilistic information about copy-number state and robustly integrate such information from individuals and their "haplotype neighbors" to call CNVs.

#### 4.1 Estimating per-SNP Bayes factors for copy-number states

Our first task was to quantify the extent to which a given SNP-array measurement—i.e., observed relative intensity ($\theta$) and total intensity (denoised LRR) for a given sample—supported the presence of a copy-gain, copy-loss, or no CNV spanning the SNP. We performed this quantification by estimating approximate Bayes factors for copy-gain vs. no CNV and for copy-loss vs. no CNV. To do so, we computed the probability density at the observed intensity data point ($\theta$, LRR) for each of the bivariate normal genotype clusters we estimated above: two probability density values for the CN=1 clusters (A, B), three for the CN=2 clusters (AA, AB, BB), and four for the CN=3 clusters (AAA, AAB, ABB, BBB). We also included a cluster that accounted for occasional CN=0 data points; we situated this cluster at a constant offset below the CN=2 het (AB) cluster, with twice its variance parameters. We then computed maximum probability densities among the values obtained from copy-gain clusters (AAA, AAB, ABB, BBB), copy-loss clusters (A, B, and CN=0), and no-CNV clusters (AA, AB, BB) and set the approximate Bayes factors for copy-gain vs. no CNV and copy-loss vs. no CNV to equal the ratios of the relevant maxima. Finally, we cropped these ratios to the range $[10^{-4}, 10^{4}]$ to limit the influence of potential outlier values.

We note that our use of maximum probability density values across genotype clusters within a copy-number state (e.g., AAA, AAB, ABB, BBB for CN=3) does not result in true Bayes factors: a formal Bayesian analysis would require a generative model that, for a given CN state, first defines a probability distribution over the genotype clusters corresponding to the CN state. We did not attempt to model the relative frequencies of genotype clusters because in practice, such modeling only becomes relevant for rare SNPs (with highly unbalanced cluster probabilities); but for such SNPs, almost all observations come from major-allele clusters, such that detailed modeling of cluster frequencies is rarely relevant. We found that in practice, CNV detection using the approximate Bayes factors we computed already increased detection sensitivity relative to previous PennCNV analyses of UK Biobank (Supplementary Tables 1-4; HI-$\text{CNV}_0$ denotes analysis using our approximate Bayes factors without incorporating information from haplotype neighbors).

An additional detail regarding our computation of bivariate normal probability density values is that we applied individual-specific scale factors to the per-noise-decile bivariate normal clusters we had estimated. We did so because even though we estimated cluster parameters separately for each LRR-noise decile of samples, the samples within a decile still exhibited varying levels of noise. To account for this remaining variation in noise, we scaled all genotype cluster standard deviation parameters for a given sample by the ratio of s.d.(LRR) in the sample to the median s.d.(LRR) in the sample's decile.

#### 4.2 Masking genotyping intensities potentially influenced by nearby SNPs

We found that for some variants on the UK Biobank SNP-array, the presence of nearby SNPs (within $\pm 30$ bp) resulted in genotyping intensities similar to deletions, presumably because the additional nearby variant caused the local sequence no longer to hybridize to either of the oligonucleotide probes for the A or B allele of the variant being genotyped. To prevent such scenarios from potentially producing spurious deletion calls, we attempted to mask all genotyping intensity measurements that might be influenced by nearby SNPs. We did so by masking, in each individual, intensity data from all variants for which a nearby SNP (within $\pm 30$ bp) had been imputed (in the UKB imp_v3 release) with an imputed dosage $>0.1$ for the minor allele. This filter removed only a small fraction of the available data: at a typical heterozygosity rate of $\sim 1$ heterozygote per 1,000 basepairs, filtering when observing a SNP in the 60 bases within $\pm 30$ bp of a genotyped variant results in a variant being filtered $\sim 6\%$ of the time.

We also applied a similar mask to multi-allelic SNPs. Intuitively, a probe designed to look for the two most common alleles at a site may make carriers of a third allele look like carriers of deletions (no signal for either of the two common alleles). As such, we masked genotype intensities for imputed carriers of a third allele at a given SNP.

#### 4.3 Hidden Markov model (HMM) using IBD-based weights

As in previous CNV-calling methods such as PennCNV [9], we used a hidden Markov model to identify sequences of consecutive SNPs at which genotyping intensity measurements consistently indicated the presence of a CNV (based on the Viterbi path passing through copy-gain or copy-loss states). Here, we needed to adapt this approach to incorporate probabilistic information not only from an individual but also from haplotype neighbors sharing IBD tracts. This task was nontrivial because fully modeling genotyping intensity data from all of these samples would require considering a combinatorial state space including the copy-number states of all haplotype neighbors (which might or might not match that of the individual in question, depending on recentness of IBD-sharing).

To retain computational tractability, we therefore incorporated information from haplotype neighbors using a simple heuristic approach somewhat analogous to a variational approximation. Specifically, at each genotyped SNP, we simply augmented the approximate Bayes factors for the individual (for copy-gain vs. no CNV and copy-loss vs. no CNV) with the corresponding Bayes factors from each haplotype neighbor, downweighted in such a way as to reflect the possibility that haplotype neighbors with shorter IBD-sharing might be too distantly related to the individual to have co-inherited a CNV. We ran this analysis using several different weighting schemes (trading off sensitivity to more recent vs. older CNV mutations, as described below) and compiled calls made across these weighting schemes (as described in the next section).

**HMM states.** We used a three-state HMM with copy-gain, copy-loss and no-CNV states. We did not attempt to have the HMM distinguish between CN=1 and CN=0 or between CN=3 and higher copy numbers given that our focus was on detecting rare biallelic CNVs.

**Emission probabilities.** Given that we ultimately wanted to perform inference based on the Viterbi path through the HMM, we could perform all computations in log space and work only with relative emission probabilities (i.e., log Bayes factors). As described above, at each SNP, our genotype cluster models allowed us to compute approximate log Bayes factors for copy-gain vs. no CNV and for copy-loss vs. no CNV from the genotyping intensities of the individual and likewise for each of the individual's haplotype neighbors. To aggregate this information into a single log Bayes factor for copy-gain (respectively, copy-loss) vs. no CNV, we computed a weighted sum in which the individual's log Bayes factor received a weight of 1 (corresponding to fully utilizing probabilistic information about CNV status from the individual's genotyping intensities) and the haplotype neighbors' log Bayes factors received weights between 0 and 1 depending on their lengths of IBD-sharing (so as to downweight information from individuals with shorter, less-confident IBD with the individual).

Explicitly, we considered a 1-parameter family of weighting functions that map a given IBD length to the probability that the time to the most recent common ancestor (TMCRA) is within $T$ generations. Intuitively, this weighting scheme optimizes for detecting CNVs that arose roughly $T$ generations ago (by incorporating information from haplotype neighbors who share more recent IBD—and thus have genotyping intensities informative of the co-inherited CNV—while discarding information from haplotype neighbors with TMRCA predating the CNV mutation). To power detection of CNVs of different ages, we ran HMM inference using six different values of

$T \in \{0, 5, 10, 25, 50, 100\}$ generations, where $T = 0$ corresponds to ignoring haplotype neighbors entirely (i.e., performing single-sample analysis). For each $T > 0$, we performed two HMM runs, incorporating information from neighbors of each of the individual's two haplotypes in turn.

To calculate the approximate probability that an IBD segment of length $l$ Morgans has TMRCA (denoted $t$) less than $T$ generations, we used the following derivation. For a population of constant size $N$, we have (from page 117 of ref. [22]):

$$P(t|l, N) = t(N^{-1} + 2l)^2 e^{-t(N^{-1}+2l)} \times (N^{-1} + 2l)\frac{t}{2}.$$

Letting $N \to \infty$, we obtain:

$$P(t|l) = t(2l)^2 e^{-t(2l)} \times (2l)\frac{t}{2}.$$

Integrating from $T$ to infinity,

$$P(t \geq T|l) = \int_T^\infty t(2l)^2 e^{-t(2l)} \times (2l)\frac{t}{2}dt = e^{-2lT}(1 + 2lT + \frac{1}{2}(2l)^2T^2).$$

Thus, the probability that an IBD segment of length $l$ Morgans has TMRCA within $T$ generations is approximately given by:

$$P(t < T|l) = 1 - P(t \geq T|l) = 1 - e^{-2lT}(1 + 2lT + \frac{1}{2}(2l)^2T^2).$$

**Transition probabilities.** We specified a transition probability matrix similar to PennCNV [9] in which the probabilities of changes in copy-number state between two consecutive probes depended on the distance between the probes (corresponding to the idea that copy-number state changes between nearby probes should be less likely than between distant probes; Supplementary Fig. 11). Explicitly, we used the transition matrix:

| | | To | | |
|---|---|---|---|---|
| | | CN=1 | CN=2 | CN=3 |
| From | CN=1 | $e^{-d_i/D_{del}}$ | $(1-10^{-4})(1-e^{-d_i/D_{del}})$ | $10^{-4}(1-e^{-d_i/D_{del}})$ |
| | CN=2 | $p_{21} = min\begin{cases}1-e^{-d_i/D}\\ \overline{\# \ del}/\# \ probes\end{cases}$ | $1 - p_{21} - p_{23}$ | $p_{23} = min\begin{cases}1-e^{-d_i/D}\\ \overline{\# \ dup}/\# \ probes\end{cases}$ |
| | CN=3 | $10^{-4}(1-e^{-d_i/D_{dup}})$ | $(1-10^{-4})(1-e^{-d_i/D_{dup}})$ | $e^{-d_i/D_{dup}}$ |

where $d_i$ is the distance between probes, $\overline{\# \ del}$ and $\overline{\# \ dup}$ are the average number of deletions and duplications called using SNP-array data (set to 15 and 5, respectively), $D_{del}, D_{dup}$ are the average lengths of deletions and duplications (both set to 100kb), $D$ is the genome size divided by the number of copy number variants (set to $3 \times 10^9/20 = 150$ Mb) and finally $\# \ probes$ is the number of SNPs on the array (set to 784,256 autosomal variants for UK Biobank).

### 5 Filtering, merging, and genotyping CNVs

In the previous sections, we described how we set up HMMs to call CNVs using information from haplotype neighbors. We incorporated such information via a set of IBD length-based weighting schemes (parameterized by a TMRCA parameter $T \in \{0, 5, 10, 25, 50, 100\}$ generations). Here we describe how we post-processed CNV calls from these HMMs to obtain a high-confidence set of CNVs (that merged calls across different values of $T$) and how we subsequently genotyped CNVs across samples.

#### 5.1 Filtering and post-processing CNV calls from each HMM run

For each individual, for each run of the HMM (parameterized by $T$ and by the haplotype of the individual used to identify neighbors), we extracted potential deletions (respectively, duplications) as consecutive sequences of copy-loss (respectively, copy-gain) states in the Viterbi path through the HMM. For each such sequence of states, we computed the $\log_{10}$ Bayes factor (BF) supporting the putative CNV event (as the sum of $\log_{10}$ BFs across the sequence of SNPs within the segment). We then applied an initial set of filters to these potential CNV segments: we required putative deletions to span at least 50 bp, and we required duplications to span at least 500 bp and have $\log_{10}\text{BF} > 9$ support.

We further post-processed the segments that survived filtering by bridging short gaps between consecutive segments of the same copy-number state (because the Viterbi path through long CNVs was sometimes interrupted by short sequences of no-CNV states). Specifically, we bridged gaps between nearby CNV segments if either (i) they included $\leq$4 probes and spanned $<$20 kb; or (ii) they spanned $\leq$20% of the combined length after bridging.

#### 5.2 Merging CNV calls across HMM runs

To synthesize post-processed CNV calls across HMM runs from different values of the TMRCA parameter $T$ (which had differing sensitivity to CNVs of different mutational ages and also exhibited stochastic variation in endpoints), we next performed a deduplication step to identify a nonredundant set of CNVs discovered in each individual. We performed this deduplication procedure on the aggregate set of CNV calls made across values of $T$ and across which of the individual's haplotypes had been used to identify neighbors. (Homozygous CNVs present on both haplotypes were collapsed into a single call during this step but handled later in a separate genotyping step described below.)

Specifically, we considered two CNV calls of the same type (DUP or DEL) to be duplicates if their endpoints matched within 3 SNP-array probes (i.e., $\Delta start \leq 3$ and $\Delta end \leq 3$). For each such duplicate pair, we retained the call with higher $\log_{10}$BF. We refer to the set of CNV calls remaining after this procedure as the "deduped" callset.

Because the deduped callset could still contain overlapping CNV calls (that were unwieldy for some downstream analyses), we also created a "unioned" callset in which we merged overlapping CNV calls of the same type (DUP or DEL). Lastly, we applied a final set of length filters on the CNV calls, requiring deletions to be >75 bp and duplications to be >500 bp (based on empirical validation analyses).

#### 5.3 Creating CNV genotypes for association tests

We used the deduped and unioned callsets described above to create genotypes for single-variant and burden-style association tests on various classes of CNVs (grouping CNVs at the probe, gene, or CNV level). In more detail:

- Probe-level tests: For each probe on the SNP-array, we used the unioned CNV callset to determine which individuals had a deletion or duplication spanning the given probe. This procedure created two 0/1 genotypes (for DEL and DUP) at each probe. (We did not distinguish homozygous from heterozygous genotypes for these tests.)

- Gene-level tests: Similarly, for all protein-coding genes, we used the unioned CNV callset to construct three gene-level 0/1 genotypes (for DEL, DUP, and pLoF):

  - Deletion (DEL): 1 if a deletion spans the entire gene (CNV boundaries $\geq$1 probe beyond first and last probe within coding sequence of gene);
  - Duplication (DUP): 1 if a duplication spans the entire gene (CNV boundaries $\geq$1 probe beyond first and last probe within coding sequence of gene);
  - Predicted loss of function (pLoF): 1 if a deletion spans any part of the coding sequence or a duplication is contained within coding sequence (i.e., CNV starting probe is at or after the first probe in coding sequence and last probe is at or before the last probe in coding sequence).

  We created these genotypes using canonical transcripts for 20,091 genes (downloaded from `https://github.com/im3sanger/dndscv/blob/master/data/refcds_hg19.rda`).

- CNV-level tests: For each CNV in the deduped callset with $\geq$5 carriers within the entire UK Biobank cohort, we constructed four versions of 0/1/2-genotypes for the CNV (parameterized by $\delta = \{0, 1, 2, 3\}$), reflecting four levels of tolerance to noise in breakpoints of CNV calls. Specifically, for a given CNV to be genotyped and a given value of $\delta$, we considered an individual to be a carrier if any run of the HMM (using any value of the TMCRA parameter $T$ and using either of the individual's two haplotypes to identify neighbors) had produced a CNV call with breakpoints that matched to within $\delta$ probes (i.e., $\Delta start \leq \delta$ and $\Delta end \leq \delta$). We considered an individual to be homozygous for a CNV if for some $T > 0$, **both** HMM runs (using neighbors from the individual's haplotype 1 and haplotype 2, respectively) had produced an approximately-matching CNV call with strong support from haplotype neighbors (i.e., neighbor-only $\log_{10}$BF $> 6$).

### 6 Quality control filtering

To obtain a robust CNV callset, we performed several stages of filtering at the sample-level, chromosome-level, and CNV-level.

#### 6.1 Individuals with trisomy 21 or blood cancer

To identify individuals with trisomy 21 we computed each individual's mean denoised LRR across probes on chromosome 21. We identified 15 individuals (in the full UK Biobank cohort) with outlier values of chromosome 21 mean LRR consistent with potential trisomy 21 and removed these individuals from analysis.

To filter individuals whose DNA samples might be affected by blood cancers or premalignant conditions, we removed all individuals who self-reported any blood cancer at assessment or had a recorded date of first occurrence of blood cancer $<$5 years after assessment.

#### 6.2 Technical artifacts producing aberrantly many CNV calls

We found that a small subset of samples with very low lymphocyte counts and red blood cell counts had aberrantly many duplication calls, apparently due to a technical artifact in LRR that had escaped denoising. We therefore filtered all samples with $>$100 CNV calls. Additionally, to identify individuals potentially affected more subtly by this type of artifact, we computed the first 10 principal components of LRR in these aberrant individuals, ranked all individuals by the amount of LRR variance explained by these artifact PCs, and removed individuals in the top 0.5% (corresponding to $>$1.1% of LRR variance explained by the 10 PCs).

#### 6.3 Chromosomes with mosaic chromosomal alterations

For individuals with a mosaic chromosomal alteration call with cell fraction greater than 20% [21], we set all probe-level, gene-level, and CNV-level genotypes on the affected chromosome(s) to missing.

#### 6.4 Somatic CNVs

We filtered calls intersecting the following regions (in hg19) frequently affected by somatic CNVs:

- Immunoglobulin genes (*IGK*: chromosome 2; 89000000 - 90274235, *IGH*: chromosome 14; 106032614 - 107288051, *IGL*: chromosome 22; 22380474 - 23265085)
- T cell receptor genes (*TRG*: chromosome 7; 38279625 - 38407656; *TRB*: chromosome 7; 141998851 - 142510972; *TRA*: chromosome 14; 22090057 - 23021075; *TRD*: chromosome 14; 22891537 - 22935569)
- *DLEU1* / *DLEU2* locus (chromosome 13; 50556688 - 51297372)

### 7 Summary measures of HI-CNV callset

#### 7.1 Validation rate of HI-CNV and PennCNV callsets

To assess the precision (i.e., validation rate) of CNVs called by HI-CNV (or PennCNV), we computed the proportion of HI-CNV (respectively, PennCNV) calls that were either (i) replicated by WGS-based CNV calls or (ii) exhibited enrichment or depletion of WGS read-depth consistent with the CNV call. We performed these analyses using whole-genome sequencing pilot data available for 43 individuals in our primary analysis set. For both the HI-CNV and PennCNV callsets, we removed calls that intersected regions that commonly contain somatic CNVs as well as all calls on chromosomes containing high-cell-fraction mosaic chromosomal alterations (see above). We note that the WGS data was aligned to hg38, whereas the SNP-array data analyzed by HI-CNV and PennCNV used hg19, so we lifted over the start and end of each HI-CNV and PennCNV call to hg38 and removed events which had an unmapped start or end.

We used CNVnator [23] to call CNVs following a standard pipeline (`https://github.com/abyzovlab/CNVnator`), using the `-unique` flag when extracting read mapping data from bam files and a binsize of 100 bp for computing WGS read-depth. We restricted to calls with a q0 (fraction of reads mapped with q0 quality) $\leq$ 0.5 (non-redundant) and a q0 not equal to -1 (couldn't be calculated). We then used the python module `pytools.io` to extract CNVnator read depth data from the root file.

For all CNVs called by HI-CNV (or PennCNV), we annotated whether CNVnator called an overlapping CNV containing at least 50% of the probes in the SNP-array-based call. We also computed mean normalized read depth across the 100 bp windows spanning the CNV being validated (normalized by read depth across entire chromosome). We then compared this mean normalized read depth to the distribution of mean normalized read depth across the same CNV region among individuals with no CNV call in the region. We used the mean and standard deviation from this background distribution to compute a $z$-score and determine if there was a significant excess or depletion of read depth (P $<$ 0.05).

The above computations allowed us to classify CNV calls into five categories containing CNVs (1) replicated by CNVnator, (2) supported by significant read-depth signal in the correct direction (e.g., depletion of read-depth for a deletion), (3) supported by non-significant read-depth signal in the correct direction, (4) with non-significant read-depth signal in the incorrect direction and (5) with significant read-depth signal in the incorrect direction. We estimated validation rate as the sum of the proportions of CNVs replicated by CNVnator, supported by significant read-depth signal in the correct direction and the excess supported by non-significant read-depth signal in the correct direction: (1) + (2) + (3) – (4).

#### 7.2 Recall of HI-CNV and PennCNV callsets

We assessed the recall, defined as the proportion of CNVs called by WGS-based analysis for which overlapping calls were made by HI-CNV (or PennCNV). We removed WGS-based calls that intersected regions that could correspond to somatic events as well as all calls on chromosomes containing high-cell-fraction mosaic chromosomal alterations. We restricted analyses to rare or low-frequency CNVs, i.e., those with AC $\leq$ 5 in the full set of 48 UK Biobank participants with available pilot WGS data. As above, we lifted HI-CNV and PennCNV calls from hg19 to hg38 and removed events which had an unmapped start or end.

We used Delly [24] to call CNVs following a standard pipeline for germline SV calling (`https://github.com/dellytools/delly`). We considered a Delly call to have been re-identified if a CNV of the same type (DEL/DUP) was called overlapping the CNV called by Delly. We assessed the recall of a variety of different subsets of CNVs (Supplementary Fig. 1 and Supplementary Table 4).

#### 7.3 Unique CNVs

For some downstream analyses, we wished to analyze the set of unique CNVs identified by HI-CNV. This task was nontrivial because a CNV mutation co-inherited by multiple individuals could be called with slightly different breakpoints in different carriers. Consequently, the set of unique CNV calls—i.e., unique pairs of (start, end) breakpoints for DELs and for DUPs—overcounted the actual number of unique mutational events identified.

To obtain a more accurate set of unique CNVs, we performed analyses to assess which CNV calls with similar breakpoints were likely to represent the same underlying CNV. Specifically, we started with CNV-level genotypes we had created for each unique CNV call (using the $\delta = 2$ version of genotyping, in which individuals with CNV call endpoints matching within $\delta = 2$

probes were considered to be carriers) and then pruned this set of CNV genotype vectors to an approximately independent subset.

Explicitly, given the complete set of $\delta = 2$ CNV-level genotypes, we computed allele frequencies and pairwise $D'$ in unrelated self-reported Europeans using PLINK [25] and clumped CNVs (with frequency $\geq 5 \times 10^{-6}$, corresponding to $\geq$5 carriers per CNV) with $D' \geq 0.5$, retaining higher-frequency CNVs. We then removed all remaining CNVs that overlapped somatic event loci (see Section 6.4).

Finally, we refined the boundaries of the remaining, independent, $\delta = 2$ CNV-level genotypes (because the breakpoints of the CNV calls used to "seed" these CNV-level genotypes could be off by 1–2 probes). To perform this refinement, for each remaining $\delta = 2$ CNV genotype vector, we identified the most common (start, end) breakpoint pair among the CNV calls that contributed to this CNV-level genotype, and we took this most-common breakpoint pair to be our best guess of the breakpoints of the underlying CNV.

### 8 Association testing and statistical fine-mapping

We ran BOLT-LMM [26, 27] to compute association statistics between CNV genotypes—at the probe, gene, and CNV level (see Section 5.3)—and 58 quantitative traits (Supplementary Data 1). We then used a pairwise linkage disequilibrium (LD)-based filter (that we previously developed for identifying likely-causal rare variant associations [28]) to remove CNV associations that could be explained by LD with a more strongly associated variant—either another CNV or an imputed SNP or indel [8, 28]—within 3 Mb of the start of the CNV.

#### 8.1 Filtering and annotating fine-mapped associations

We annotated all CNV-phenotype associations that passed our LD-based fine-mapping filter (involving either probe-, gene-, or CNV-level tests) with the following information (Supplementary Data 2):

- Trait associated with the CNV (`trait`)
- Lead CNV (i.e., the tested CNV genotype vector with highest $\chi^2$ association statistic) and tied CNVs (all tested CNVs that had identical $\chi^2$ value as lead CNV; `leadCNV,tiedCNVs`)
- Number of carriers and allele frequency among self-reported European UK Biobank participants (`nCarriers,A1FREQ`)
- Genomic location: the chromosome, the median start and end location among CNV calls considered in the test, size (in kb) using the median start and end location; the median loca-

tion of the probe before and after the CNV calls (`Chr, medianStart, medianEnd, size_kb, median_loc_beforeStart, median_loc_afterEnd`)

- Genic context: all genes intersecting the interval between the median start and end (either considering full genes or only exons; `genes_exon_or_intron,gene_exons`), genes that intersect an expanded interval $\pm$ 100 kb (`genes_100kb`).

- Effect size and association strength: beta (effect size), standard error, $\chi^2$ and $P$ from BOLT-LMM (`BETA,SE,CHISQ_BOLT_LMM,P_BOLT_LMM`)

- Nearby SNP associations: most associated SNP (imputed in the UK Biobank imp_v3 release or from WES [28]) within 1Mb. For each such SNP, we annotated the ID of the SNP, the genomic location, the effect size, the P-value, and the $\chi^2$ statistic from BOLT-LMM (`mostAssocimpv3_SNP_1Mb, mostAssocimpv3_SNP_1Mb_BP, mostAssocimpv3_SNP_1Mb_beta,mostAssocimpv3_SNP_1Mb_P, mostAssocimpv3_SNP_1Mb_CHISQ, mostAssocWES_SNP_1Mb, mostAssocWES_SNP_1Mb_BP, mostAssocWES_SNP_1Mb_beta, mostAssocWES_SNP_1Mb_P, mostAssocWES_SNP_1Mb_CHISQ`)

We filtered all associations that involved CNVs that overlapped regions prone to somatic CNVs (see Section 6.4). We also filtered associations in the MHC region that had escaped our pairwise LD-based fine-mapping filter due to subtle differences in the genetic principal components we used as covariates in these analyses vs. the PCs that we had previously used as covariates when computing association test statistics for SNPs and indels [28]. We verified (using linear regression analyses) that the difference in PCs only affected a small number of associations in the MHC region, at which long-range LD influenced one set of PCs more than the other.

#### 8.2 CNVs contributing to likely-causal phenotype associations

Most of the CNV-phenotype associations that passed our fine-mapping filters (and were thus deemed likely-causal) involved burden-style tests: probe-level tests that considered all DELs or DUPs spanning a genomic position, and gene-level tests that considered all CNVs with a particular effect on a gene. CNV-level tests could also potentially include multiple distinct CNVs with slightly different breakpoints. We therefore undertook further analyses to roughly identify which unique CNVs underlay each association.

For each trait, we identified all $\delta = 2$ CNV-level genotype vectors associated at nominal significance ($P < 0.05$). We then subsetted to genotype vectors that appeared to contribute to the association of interest, based on satisfying three additional criteria: (1) $D' \geq 0.75$ with the CNV genotype of interest (be it a probe, gene, or CNV level test), (2) MAF $< 2\times$ MAF of the CNV genotype of interest, and (3) length > $\frac{1}{2}$ median size of the CNV genotype of interest. Finally, among the remaining $\delta = 2$ CNV-level genotypes, we pruned to an independent set following the same approach we used to identify unique CNVs (see Section 7.3).

The above procedure produced a satisfactory set of unique CNVs underlying most phenotype associations, but for a few associated CNV genotypes that were very rare or combined deletions and duplications (specifically, pLoF gene-level tests), no $\delta = 2$ CNV-level genotype was both in high $D'$ with the CNV genotype of interest and nominally associated with the trait. In these instances, we did not attempt to further identify specific unique CNVs contributing to the association.

##### 8.3 Defining CNV loci

The above approach identified a set of CNVs likely to contribute to causal phenotype associations. To group these CNVs into loci, we sorted the CNVs by increasing size. For each chromosome, we denoted the smallest CNV on the chromosome as belonging to "locus1" and then iterated through other CNVs on the chromosome in order of size. For each CNV in turn, if it overlapped or fell within $\pm 100$ kb of one or more previously-defined loci, we annotated it as belonging to those loci, and otherwise we considered it create a new locus.

##### 8.4 Syndromic and non-syndromic loci, CNVs, and associations

We annotated a likely-causal CNV as syndromic if it overlapped a previously-curated pathogenic CNV (from the set of 92 pathogenic CNVs curated by [29]) by more than 50%. We annotated a locus as syndromic if any CNV assigned to only that locus was annotated as syndromic. To annotate a CNV-phenotype association as being syndromic or non-syndromic, we examined all likely-causal CNVs that belonged to a single locus and contributed to the association and annotated the association as syndromic if at least one such CNV was syndromic.

#### 9 Follow-up analyses at loci of interest

Here we provide details of additional analyses we performed at loci of interest, including refined analyses of specific phenotypes, corroborating analyses of SNP and indel PTVs, and further characterization of specific CNV events.

##### 9.1 Extreme blood phenotypes

The CNV-phenotype association tests we ran using BOLT-LMM analyzed blood cell traits that we had previously normalized using an approach that included removal of outliers, defined as deviating from the median by >7x the interquartile range (IQR). However, we subsequently found that certain CNVs had large enough effect sizes that a substantial fraction of carriers had been removed as outliers. As such, when further investigating loci related to blood traits, we renormalized blood phenotypes without outlier removal (using covariate adjustment and inverse normal transforms as previously described) [28].

#### 9.2 Residualization of phenotypes to emulate mixed model analysis

In follow-up analyses (e.g., of SNP and indel PTVs genotyped in a subset of individuals, or for categories of CNVs we did not initially genotype), we performed linear regression on phenotypes that we residualized for polygenic predictions using array-typed SNPs (omitting those within 2Mb of the gene of interest) that we generated using BOLT-LMM (`--predBetasFile`) in 10-fold cross-validation (to emulate the power of linear mixed model association analysis) [30]. We normalized residualized phenotypes to have a mean of zero across all non-removed individuals with non-missing phenotype.

#### 9.3 PTVs in UK Biobank exome sequencing data

We identified carriers of high-confidence loss-of-function SNP and indel variants (on canonical transcripts annotated using LOFTEE [31]) from the 185,365 UK Biobank participants in our analysis set with whole-exome sequencing data available [32]. However, for *R3HDM4* we analyzed carriers of high-confidence loss-of-function SNP and indels in any transcript as there were no high-confidence loss-of-function SNP and indels on the canonical transcript.

#### 9.4 $\alpha$-globin locus

Exons of *HBA2* are located at 16:222911-223006; 16:223123-223328; and 16:223470-223599 whereas for *HBA1* they are at 16:226715-226810, 16:226927-227132, and 16:227281-227410 (hg19 coordinates). The UK Biobank SNP-array contained 3 probes within either *HBA2* or *HBA1*, with genomic coordinates listed as 227306, 227333, and 227365 (all within the last exon of *HBA1*, and all with extremely rare minor alleles). Due to the sequence similarity of *HBA2* and *HBA1* these 3 probes effectively measured copy number of both *HBA2* and *HBA1*. The probe before these probes was at 221057 and the one after them was at 228306. HS-40 is located 40 kb upstream of the zeta-globin gene, around 162686.

Given the above information, we categorized CNV calls at the $\alpha$-globin locus as follows:

- Alpha-globin locus DEL: a deletion call with a start $\leq$ 140000 and end $\geq$ 230000.
- HS-40 DEL: a deletion call with a start $\leq$ 162240 and end $\geq$ 162240 and $<$ 226715.

- *HBA2+HBA1* DEL: a deletion call with a start of 205897 and end of 231021, or a deletion with a start of 216041 and end of 228306 or 231021.

- *HBA2* DEL: a deletion call with a start of 221057 and end of 227306 (indicating an -$\alpha^{4.2}$ deletion; Supplementary Fig. 12).

- *HBA2* DUP: a duplication call with a start of 221057 and end of 227306 or 227333 (indicating an $\alpha\alpha\alpha^{anti\ 4.2}$ duplication; Supplementary Fig. 12).

- *HBA2* triplication: a duplication call with a start of 221057 and end of 227365 (suggesting an $\alpha\alpha\alpha\alpha^{anti\ 4.2}$ triplication; Supplementary Fig. 12). Whole-exome sequencing read-depth for carriers of such events confirmed triplication of *HBA2* (Supplementary Fig. 13).

- *HBA2+HBA1* DUP: a duplication call with a start $\geq$ 176743 and $\leq$ 221057 and end $\geq$ 230000.

- Alpha-globin locus DUP: a duplication call with a start $\leq$ 140000 and end $\geq$ 230000.

#### 9.5 Retroposition of spliced *MTMR2* transcript into an intron of *LRCH1*

Our callset included a duplication call in *MTMR2* on chromosome 11 with length ~10-20kb that was called in 2,522 UK Biobank participants (MAF=0.003). This variant associated with an increase in platelet distribution width of +0.12 (0.02) s.d. ($P = 1.7 \times 10^{-10}$) and passed our LD-based fine-mapping filter, with no nearby SNP on chromosome 11 reaching genome-wide significance. Surprisingly, this event was not called in gnomAD-SV [33] or the 1000 Genomes 30x SV callset [34], prompting further investigation.

Examination of sequencing reads from exome-sequenced carriers showed that the event was actually a retroposed pseudogene insertion of the *MTMR2* processed transcript into an intron of *LRCH1* on chromosome 13. We observed increases in read coverage only in exons of *MTMR2* and split reads corresponding to splice junctions (usually seen in RNA-seq data rather than DNA sequencing). Split reads that partially aligned to the 5' UTR of *MTMR2* and partially aligned to chromosome 13 showed that that the *MTMR2* transcript had been inserted into an intron of *LRCH1*.

Closer examination of UK Biobank SNP-array probes at *MTMR2* contributing to the initial signal showed that an "indel" probe (Affx-52351109) actually directly genotyped the retroposed pseudogene insertion. Carriers of the duplication calls exhibited increased LRR at seven probes (not usually enough to sensitively call a duplication event—suggesting that MAF=0.003 was an underestimate, representing calls in only a subset of carriers). Six of the probes with increased LRR fell within coding exons or UTRs, as expected; the remaining probe (Affx-52351109, intended to genotype an indel 11:95595151:TTTA>T) fell just within intron 7-8 of *MTMR2*, 2bp beyond the end of exon 7. Inspection of the *MTMR2* transcript showed that the minus-strand sequence ending in this "indel" actually corresponds to the splice junction created by joining exon 7 to exon 8. Further analysis of LRR at the seven probes confirmed that Affx-52351109 directly genotyped the retroposed insertion (identifying an expanded set of carriers; MAF=0.007). The 3bp indel that the probe was designed to genotype does not actually exist according to gnomAD [31].

Analyses of population allele frequencies and linkage disequilibrium of the *MTMR2* retroposed insertion showed that the variant sits on a European haplotype (MAF=0.7%) containing rs145057384, a good tag SNP ($R$=0.87, MAF=1%). Allele frequencies in UK Biobank (based on Affx-52351109) were 0.69% in Europeans and 0.02-0.05% in non-Europeans (SAS, AFR, EAS). The insertion was also called in the 1000 Genomes 30x SV callset [34], which contains a 2,529 bp insertion consisting of most of the processed transcript of *MTMR2* (excluding some 3' UTR sequence typically present in transcripts according to GTEx v8 data [35]), plus a poly-A tail, followed by another 15bp; the 1000 Genomes data set contained 11 carriers among N=3,202 individuals (10 EUR + 1 Colombian).

Our next question was whether the retroposed insertion affected platelet traits by disrupting *LRCH1* in some way. *LRCH1* LoFs were too rare to evaluate the effect of LoF on platelet distribution width (PDW), so we focused on investigating potential effects of *LRCH1* variants on gene expression or splicing.

*LRCH1* is broadly expressed in many tissues, and 11 carriers of the insertion in GTEx v8 [35] appeared to have reduced *LRCH1* expression. Carriers were identified based on chimeric sequence that we detected in 11 of 13 carriers of the tag SNP rs145057384. RNA-seq data was available for 0-8 carriers per GTEx tissue. Among the 25 tissues with RNA-seq data available for 4+ carriers (providing reasonable power), 22 of 25 tissues exhibited negative mean normalized expression of *LRCH1* in carriers ($P = 1.6 \times 10^{-4}$; two-sided sign test). We were unable to determine a mechanism by which this $\sim$2.5kb insertion might reduce expression: the inserted *MTMR2* processed transcript does not appear to be transcribed (based on no evidence of expression of the truncated 3' UTR), consistent with it lacking a promoter, and the insertion does not appear to affect splicing.

A common-SNP association with PDW (in a different intron of *LRCH1*) also appeared to be mediated by *LRCH1* expression (Supplementary Fig. 6). Interestingly, the association of the retroposed pseudogene insertion with PDW—which our GTEx analyses suggested was likewise *LRCH1* expression-mediated—exhibited $\sim$4-fold larger effect sizes on *LRCH1* expression and PDW than the common SNPs (Supplementary Table 16).

### 10 Contrasting effect sizes of deletions and duplications

#### 10.1 Selection of gene-trait pairs with likely-causal rare coding variants

To explore the relative effects of focal deletions and duplications, we examined 199 gene-trait pairs for which we had previously identified PTVs likely to alter quantitative traits (Supplementary Table 3 of [28]). For each gene on this list, we then compared the effects of likely-causal PTVs to those of whole-gene deletions and duplications.

At the level of individual loci, gene deletions acted similarly to PTVs; of the 41 genes for which there were at least 2 carriers of gene-deletions, 16 deletions were nominally significant for the given trait and 6 were Bonferroni significant (Fig. 5). At the level of individual loci, gene duplications tended to act in the opposite direction as PTVs and with a smaller magnitude of effect; of the 139 genes for which there were at least 2 carriers of gene-duplications, 27 duplications were nominally significant for the given trait and 3 were Bonferroni significant (Fig. 5).

#### 10.2 Comparison of deletion and duplication effect sizes: power analysis

Consistent with the idea that duplications tend to have a weaker effect, there were far more examples of gene duplications than gene deletions with at least 2 carriers (139 vs. 41; Fig. 5). We next wished to quantify the difference in effect sizes. For each of the 199 gene-trait pairs we could assess whether at least two individuals in UK Biobank carried a gene deletion or duplication, and for these events compare the effect sizes of likely-causal PTVs to the gene deletions and duplications.

More concretely, for a given trait $t$, and gene $g$, we are given:

- $\hat{\beta}_{CNV,g-t}, se(\hat{\beta}_{CNV,g-t})$ for $CNV = \{DEL, DUP\}$
- Number of carriers of $CNV = \{DEL, DUP\}$ ($\geq 2$)
- Sample size (N)
- Increase in effective sample size from using BOLT-LMM (equivalently, residualizing on genome-wide SNPs reduces $\sigma_{trait}$ to $< 1$); in BOLT-LMM output files the line "Absolute prediction MSE, fold-best" contains an estimate of BOLT-LMM's $\sigma^2_{trait}$ (after conditioning on genome-wide SNPs); denoted $boltlmm_{boost}$
- Given multiple PTVs indexed by $i$, we compute the inverse variance weighted mean effect:

$$\hat{\beta}_{PTV,g-t} = \frac{1}{\sum_i 1/se(\hat{\beta}_{PTV_i,g-t})^2} \sum_i \hat{\beta}_{PTV_i,g-t}/se(\hat{\beta}_{PTV_i,g-t})^2;$$

$$se(\hat{\beta}_{PTV,g-t}) = \sqrt{\frac{1}{\sum_i 1/se(\hat{\beta}_{PTV_i,g-t})^2}}.$$

For each trait-gene pair, we can compute the power ($power_{g-t,CNV}$) for two sample (different sizes) t-test of means assuming the effect size $d = |\mu_{CNV} - \mu_{nonCNV}|/\sigma_{trait} = |f \cdot \hat{\beta}_{PTV,g-t}|/\sqrt{boltlmm_{boost}}$ (with $f = \{0, 0.5, 1\}$), significance level 0.05, and the number of carriers and non-carriers for a given $CNV = \{DEL, DUP\}$.

For a given trait-gene pair, assuming independence across gene-trait pairs, we can consider the random indicator variable of whether a significant effect was seen for the $CNV = \{DEL, DUP\}$; $\mathbb{1}(p_{\beta_{g-t,CNV}} < 0.05) \sim Ber(power_{g-t,CNV})$. Across all trait-gene pairs we can then consider the observed number of significant $CNV$ effects:

$$\Gamma_{CNV} = \sum_{g,t} \mathbb{1}(p_{\beta_{g-t,CNV}} < 0.05) \sim \text{Poisson binomial}.$$

We can then compare the expected number of significant $CNV$ effects for $f = \{0, 0.5, 1\}$ to the number of observed significant $CNV$ effects. We note that this approach ignores the sign of effect size (e.g., whether duplications have opposite vs. same effect directions as PTVs). Results were consistent with deletions having similar effect sizes as PTVs; assuming deletions had the same effect size as PTVs resulted in 18.5 expected nominally associated associations whereas assuming half the magnitude of PTVs resulted in 8.3 expected associations (Fig. 5). Similar power analysis results for gene duplications show results are consistent with duplications having the opposite direction, and a smaller magnitude compared to the PTV-effect (Fig. 5).

An extension of this approach is to search across the space $0 \leq f \leq 1$, and for each value compute the expected value of number of significant associations and find the value for which $f$ results in the observed number of significant associations (Supplementary Fig. 7 and Supplementary Table 18).

As a sensitivity analysis we further performed likelihood-based analyses. We can compute the likelihood of observing $c \cdot \hat{\beta}_{PTV,g-t}$ assuming it came from $\sim N(\hat{\beta}_{CNV,g-t}, se(\hat{\beta}_{CNV,g-t}))$; assuming independence across gene-trait pairs, we can then compute the maximum likelihood estimate for $c$. We note that this approach incorporates the sign of effect size; however, one can also ignore the sign and quantify the absolute effect (agnostic to effect direction) by computing the likelihood of observing $c \cdot |\hat{\beta}_{PTV,g-t}|$ assuming it came from $\sim N(|\hat{\beta}_{CNV,g-t}|, se(\hat{\beta}_{CNV,g-t}))$. We note that this approach ignores the standard error of $\hat{\beta}_{PTV,g-t}$; however, these PTVs come from a published set of significant ($P < 5 \times 10^{-8}$) variants and therefore the standard error can be considered to be much smaller than that of $\hat{\beta}_{CNV,g-t}$. Results can be found in Supplementary Fig. 7 and Supplementary Table 18.

**Supplementary Figure 1. Detection sensitivity (recall) of SNP-array based methods on a benchmark set of CNVs called from WGS data.** We assessed recall of low-frequency (MAF$\leq$5%) CNVs called by Delly in 43 individuals with whole-genome-sequence data available. Sensitivity increased with CNV size and probe overlap (left-to-right columns) and for gene-overlapping CNVs (bottom vs. top row). HI-CNV$_0$ denotes analysis without incorporating information from haplotype neighbors (but still using our SNP-specific probabilistic models of genotyping intensities). Numerical results are available in Supplementary Table 4.

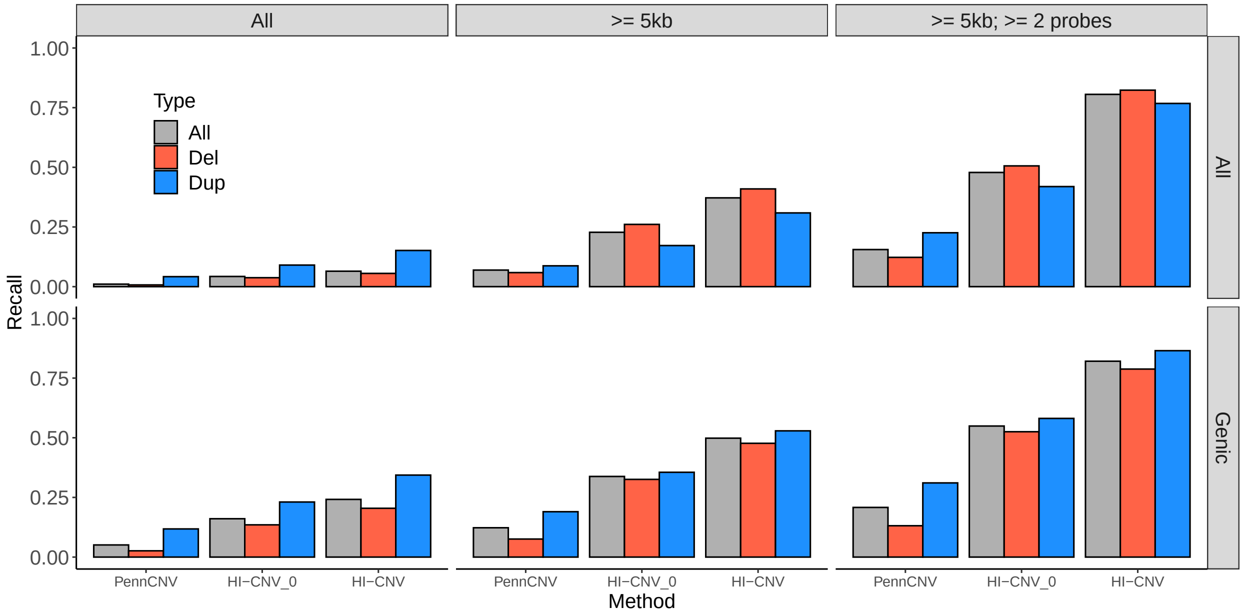

**Supplementary Figure 2. Minor allele frequency (MAF) distribution of unique CNVs detected by HI-CNV.** Red dashed line indicates MAF=0.05. The callset is depleted for common CNVs, probably due to a combination of SNP-array probe placement (avoiding common CNV regions) and less-accurate genotype cluster priors for SNPs within common copy-number polymorphisms. Numerical quantiles are shown in Supplementary Table 5.

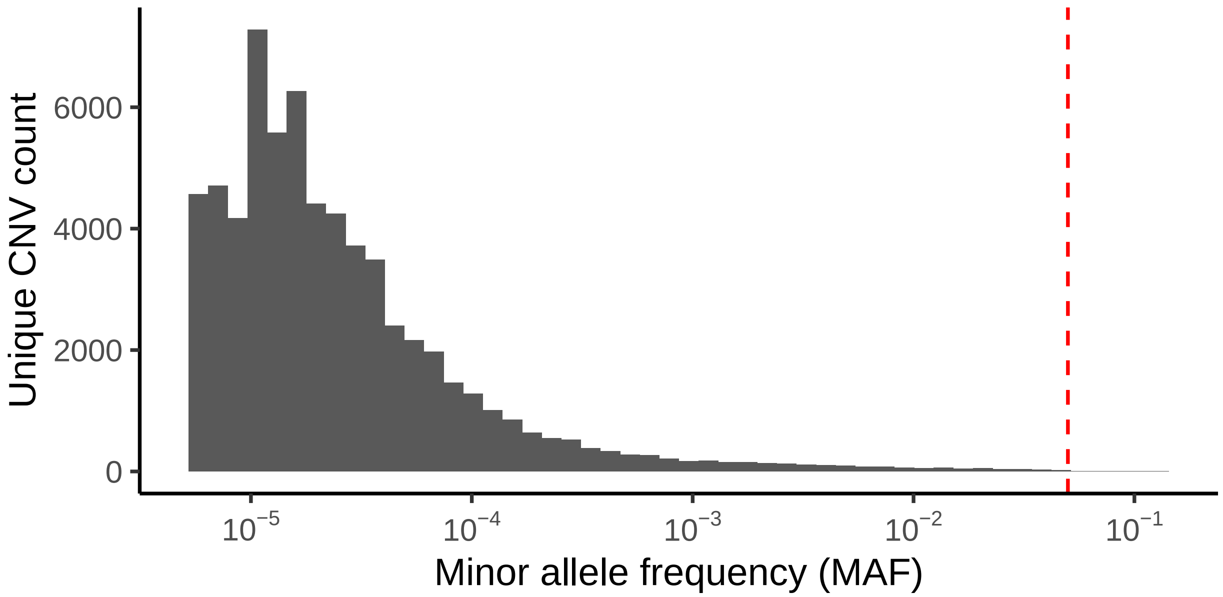

**Supplementary Figure 3. Sensitivity of PennCNV to detect CNVs contributing to likely-causal phenotype associations.** For each of the 269 likely-causal CNV-phenotype associations found via analysis of our HI-CNV callset, the number of carriers found using PennCNV versus HI-CNV is plotted. The four panels stratify the data by CNV size (median among the CNV calls that were considered in each association test). The dashed line shows the $y$=$x$ line.

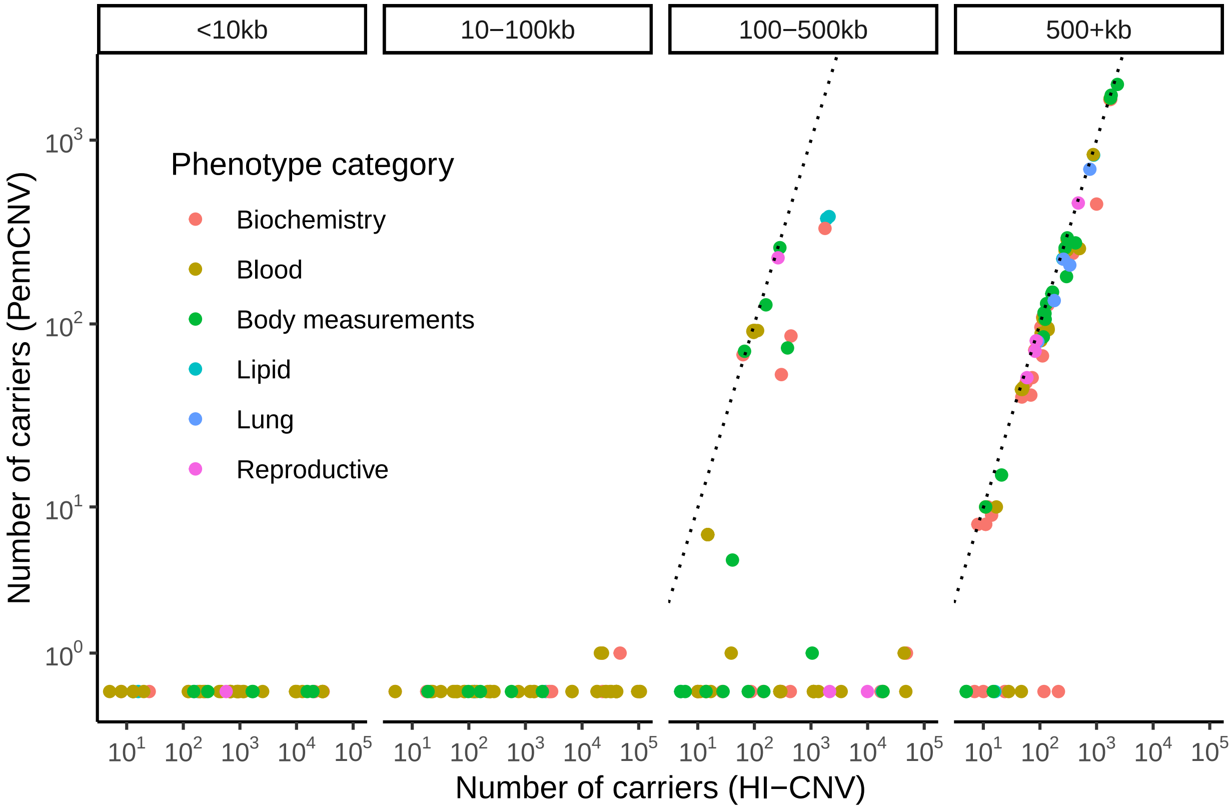

**Supplementary Figure 4. LRR signal for individuals with *SLC2A3* deletion calls shows no evidence of homozygous deletions.** Distributions of mean LRR (denoised and rescaled) across 60 probes spanned by reciprocal *SLC2A3* DEL/DUP CNVs are shown for carriers of *SLC2A3* deletion calls (3,768 individuals), *SLC2A3* duplication calls (16,821 individuals), and 1,000 randomly selected controls. The mean LRR histogram for duplications has a heavy right tail suggesting the presence of individuals with CN=4, whereas the mean LRR histogram for deletions shows no evidence of homozygous deletions (binomial $P$=0.0009 assuming Hardy-Weinberg equilibrium).

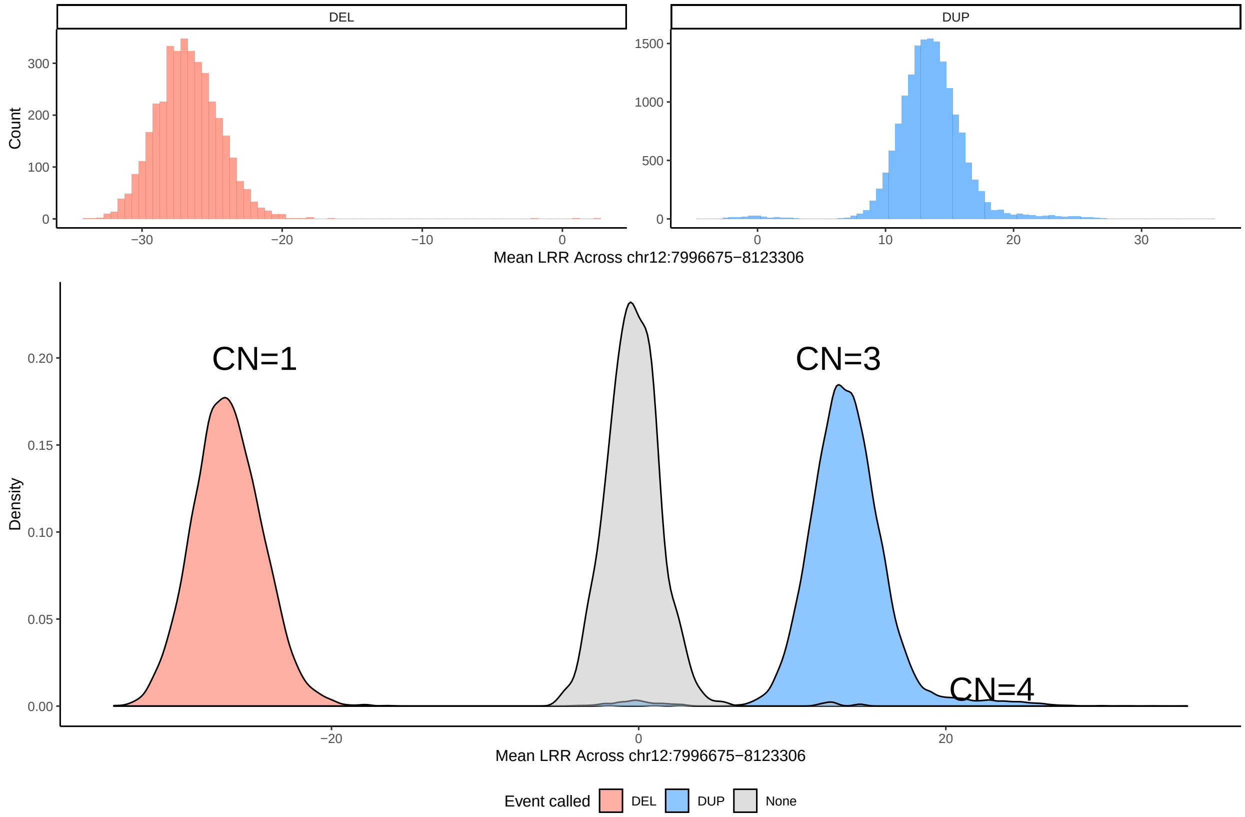

**Supplementary Figure 5. CNVs at the $\alpha$-globin locus and their effects on blood traits.** For carriers of each category of $\alpha$-globin CNVs shown in Fig. 4a, we plot the average mean corpuscular hemoglobin (MCH; both raw and units of standard deviation; s.d.), red blood cell count (RBC; both raw and units of s.d.), mean corpuscular volume (MCV; both raw and units of s.d.), and the Mentzer index (MCV/RBC). Vertical lines denote population averages.

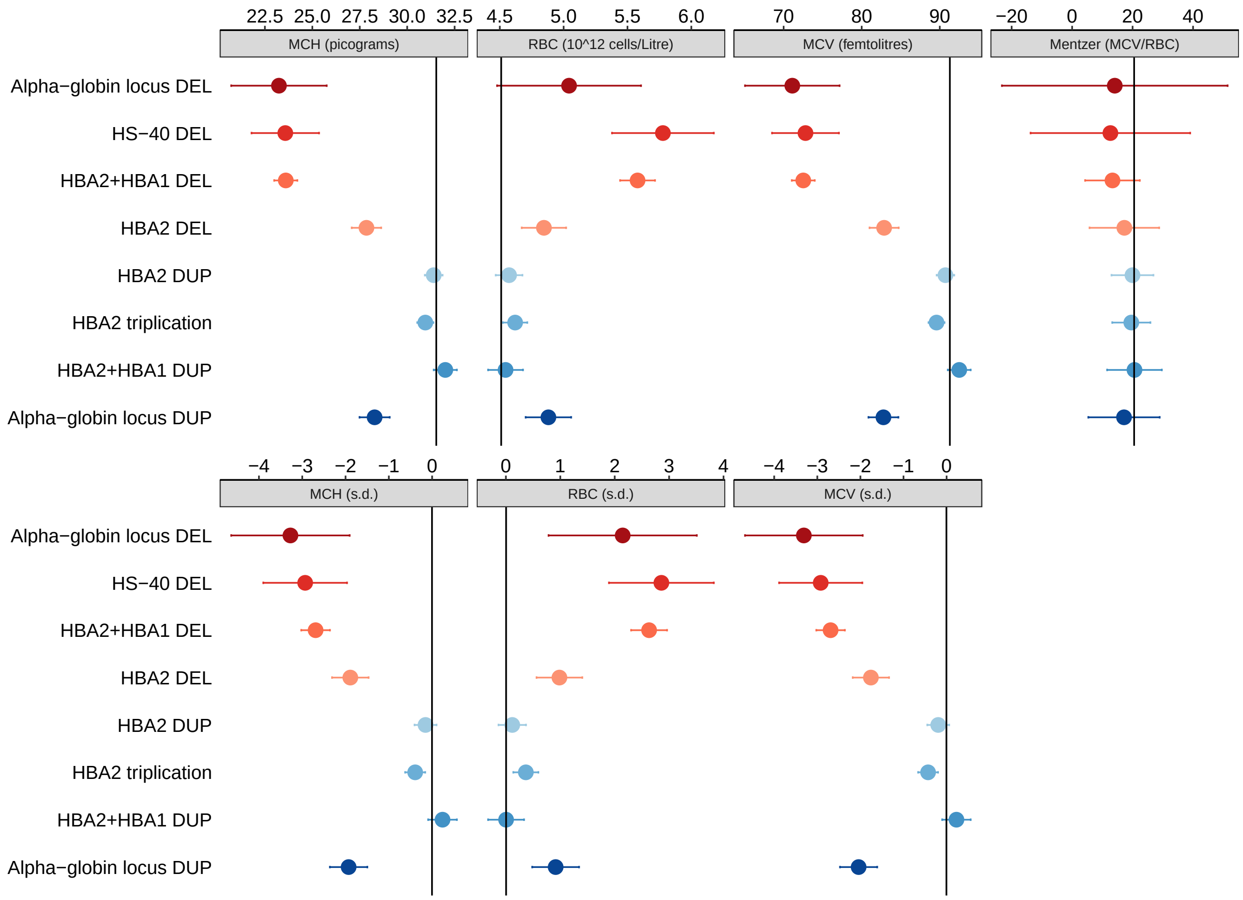

**Supplementary Figure 6. A common-SNP association with platelet distribution width appears to be mediated by *LRCH1* expression.** Top: Manhattan plot of associations between SNPs and platelet distribution width (linear regression $P$-values computed using UK Biobank participants of self-reported European ancestry). Bottom: *LRCH1* eQTL $P$-values from GTEx for the tissues with the top three signals. The red points toward the right of each plot correspond to the retroposed pseudogene insertion (directly genotyped by Affx-52351109 in the top plot) and the tag SNP rs145057384 in the bottom plots; the insertion has ~4-fold larger effect sizes than the top SNPs but has weaker associations because it is much rarer.

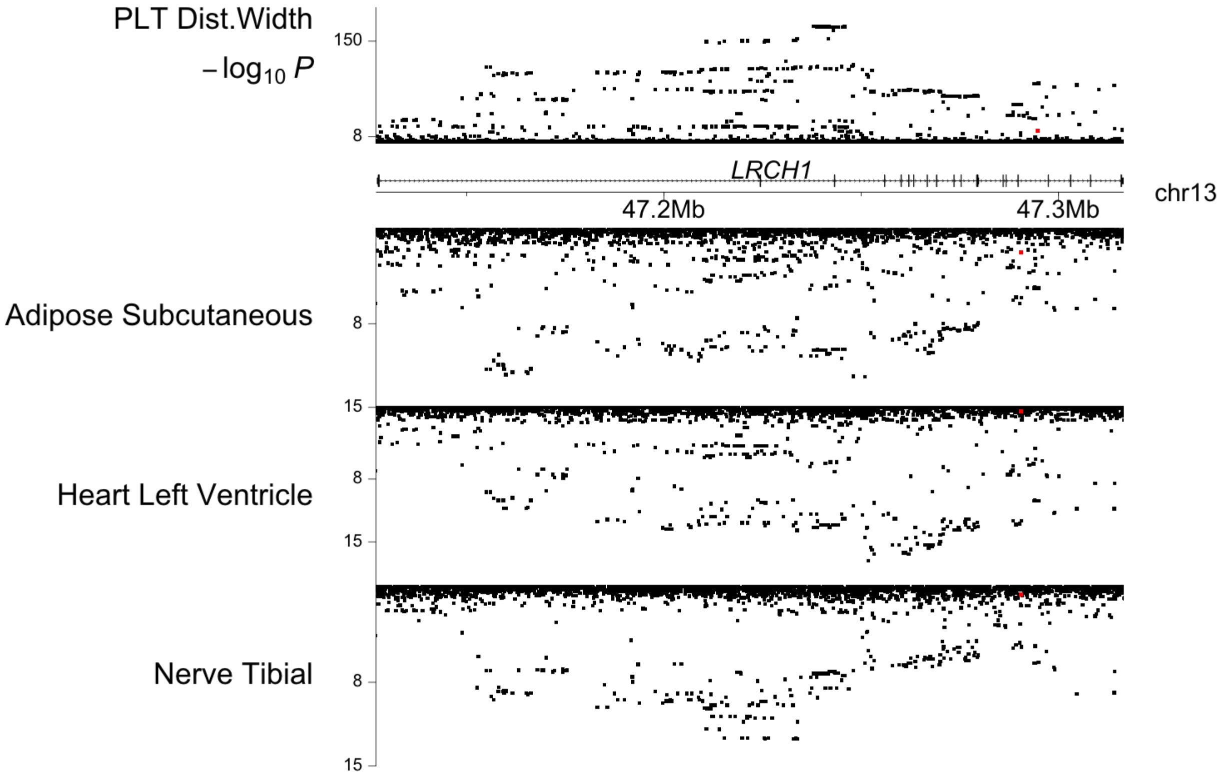

**Supplementary Figure 7. Estimation of effect sizes of duplications and deletions relative to PTVs.** Numerical estimates and further information about analyses are provided in Supplementary Table 18.

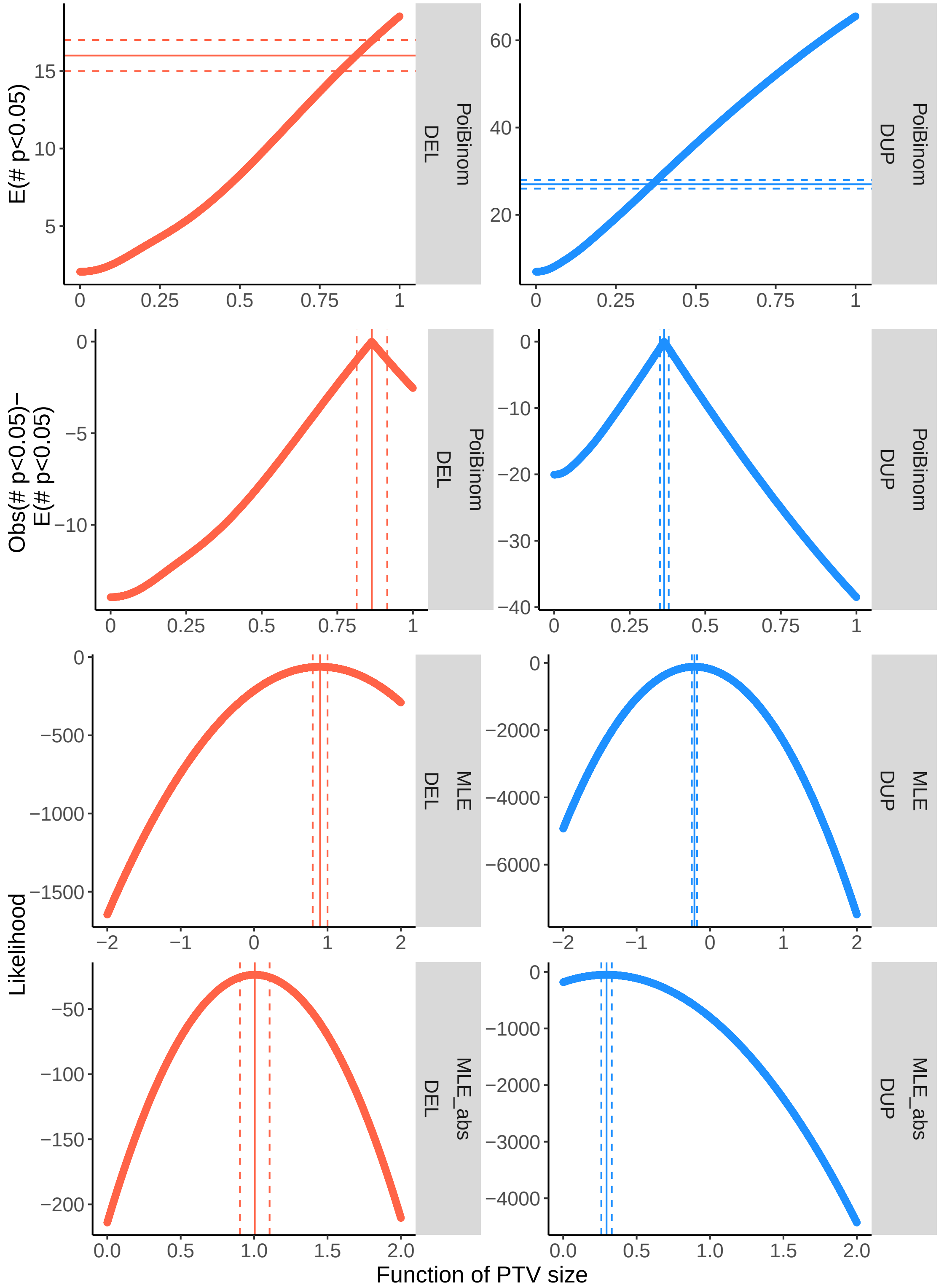

**Supplementary Figure 8. Examples of SNP-specific genotype cluster predictions for copy numbers 1, 2, 3.** The axes of each plot are $\theta = \frac{2}{\pi} \arctan \frac{B}{A}$ ($x$-axis, analogous to BAF), and LRR ($y$-axis), scaled to $[-127, 127]$. Darker, bold 95% confidence ellipses indicate bivariate normal clusters estimated directly from assigned data points (based on Affymetrix genotype calls, for CN=2 clusters (green); or based on large CNV calls, for CN=1 (red) and CN=3 (blue) clusters). Lighter, thin 95% confidence ellipses indicate predicted bivariate normal clusters. Plotted data points correspond to ($\theta$, LRR) pairs from SNPs within large CNV calls. Each subplot shows data from the 5th noise decile for a different example SNP on chromosome 7 (with $m$ indexing into the plink bim file for chr7); "Ndel" and "Ndup" indicate the numbers of large CNV calls providing data for the two CN=1 and four CN=3 clusters, respectively.

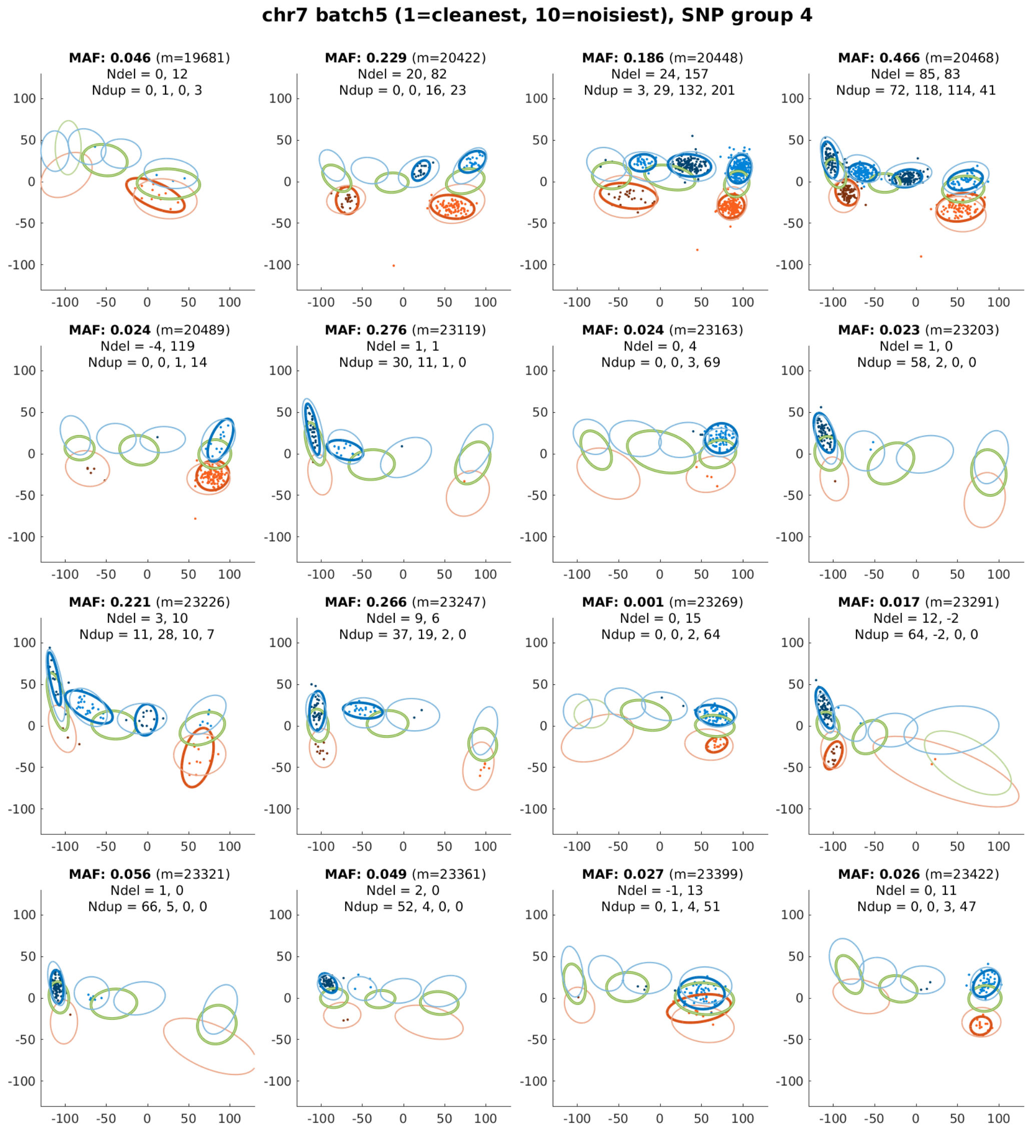

**Supplementary Figure 9. Fractions of LRR variance explained by top 10, 20, . . ., 100 principal components.** For each of the 106 genotyping batches in turn, principal components were computed on LRR values for all autosomal variants, restricting to white British samples. Fractions of LRR variance explained by the top 10, 20, . . ., 100 PCs are plotted per batch (grey dots) and averaged across all batches (black dots). Numerical results are available in Supplementary Table 19.

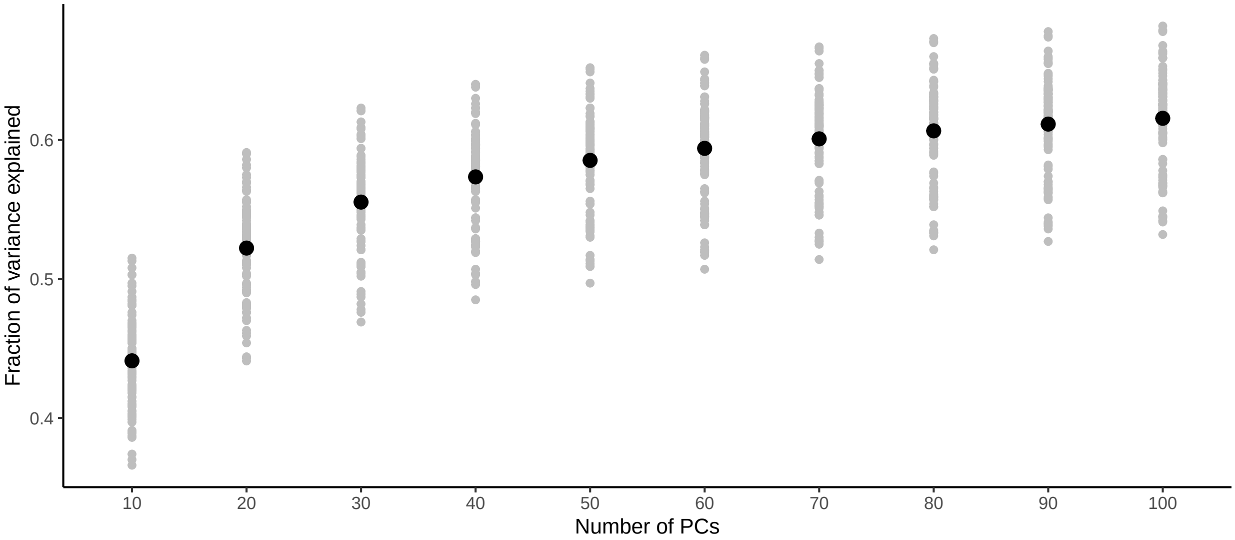

**Supplementary Figure 10. Lengths of haplotypes shared by top "haplotype neighbors."** These data describe the length distribution (across SNPs on the array and the two haplotypes of each UK Biobank participant) of the $n$-th longest haplotype match. Numerical results are available in Supplementary Table 20.

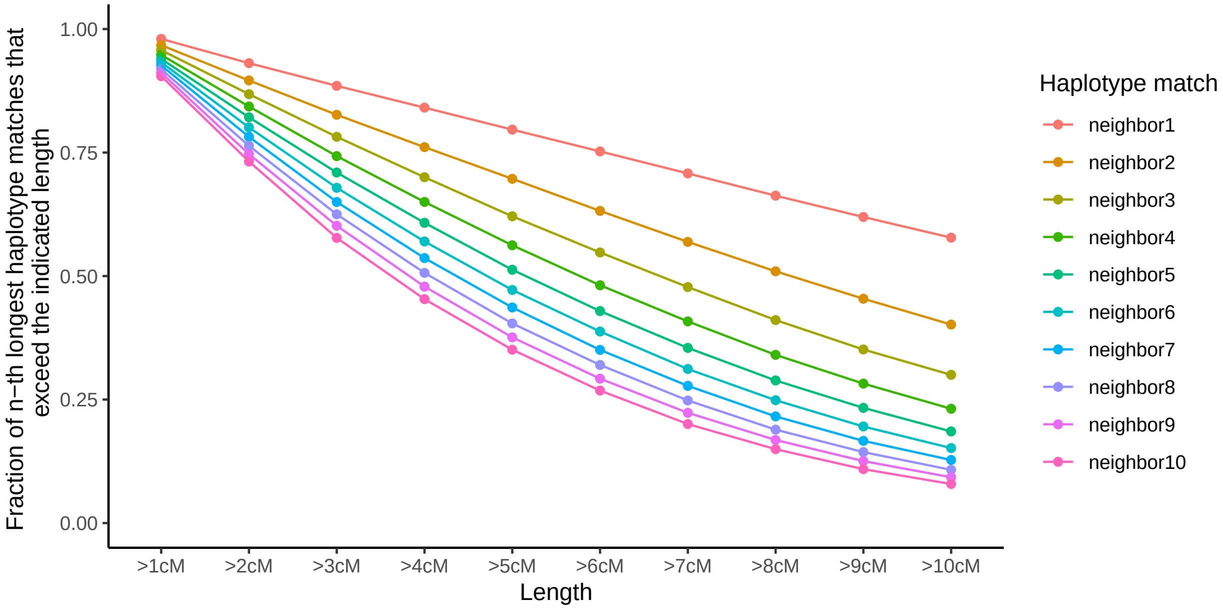

**Supplementary Figure 11. Hidden Markov model transition probabilities as a function of distance between probes.**

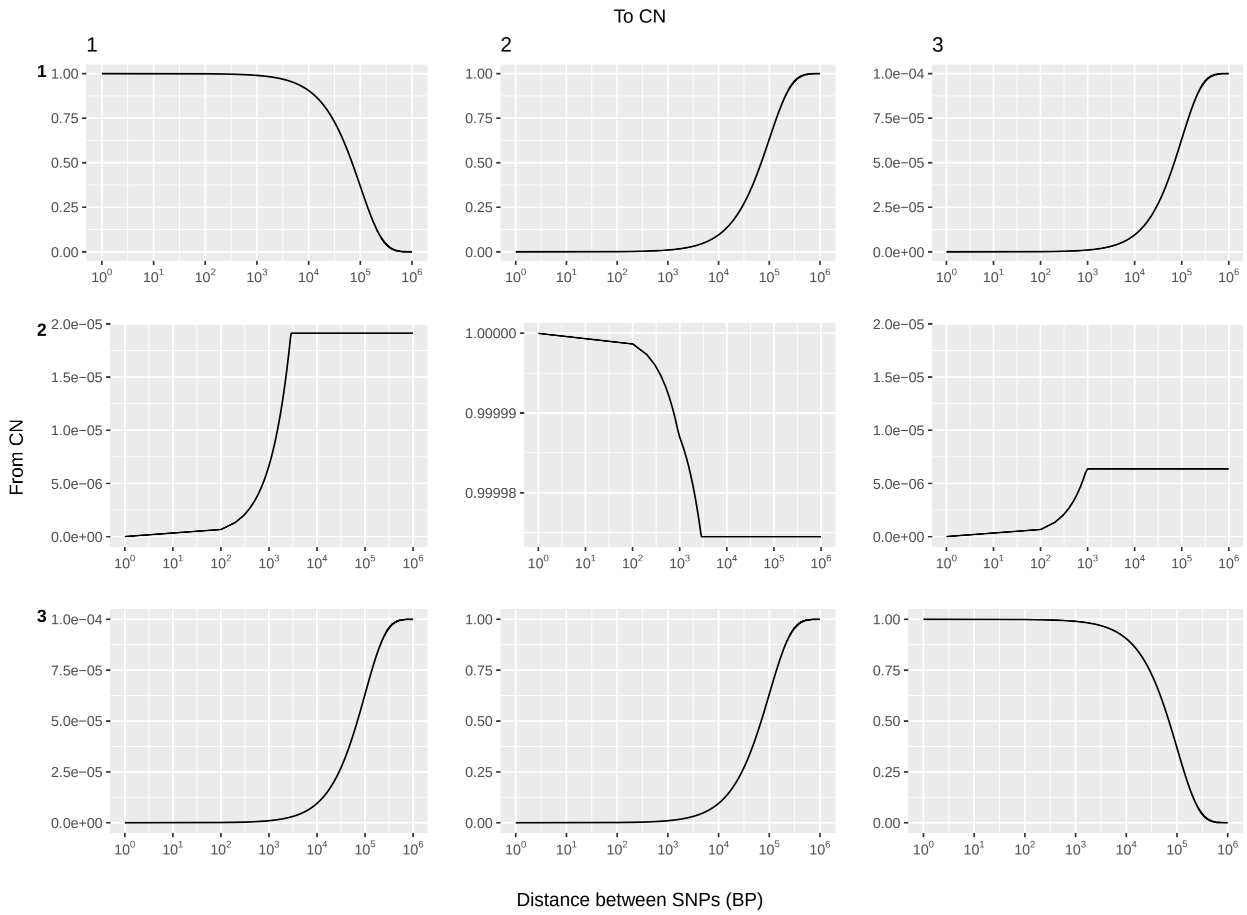

**Supplementary Figure 12. Schematic of non-allelic homologous recombination (NAHR) at the $\alpha$-globin locus.** The *HBA2* and *HBA1* genes lie within $\sim$4 kb regions containing three distinct regions of high homology (X, Y, and Z boxes; Fig. 1 of ref. [36]). The spacing between the X and Y boxes differs in the sequence left of *HBA2* vs. *HBA1*, such that NAHR within the X boxes creates 4.2-kb DEL/DUP events ($-\alpha^{4.2}$ and $\alpha\alpha\alpha^{anti\ 4.2}$) while NAHR within the Z boxes creates 3.7-kb DEL/DUP events ($-\alpha^{3.7}$ and $\alpha\alpha\alpha^{anti\ 3.7}$). The UK Biobank SNP-array contained a probe in unique sequence at 16:221057 between X2 and Y2 (pink star). Because intensity at this probe is affected by 4.2-type events but unaffected by 3.7-type events, we concluded that duplication calls starting at the probe 16:221057 and extending through the three probes in *HBA* exons were probably 4.2-type events that involved full deletion or duplication of *HBA2*.

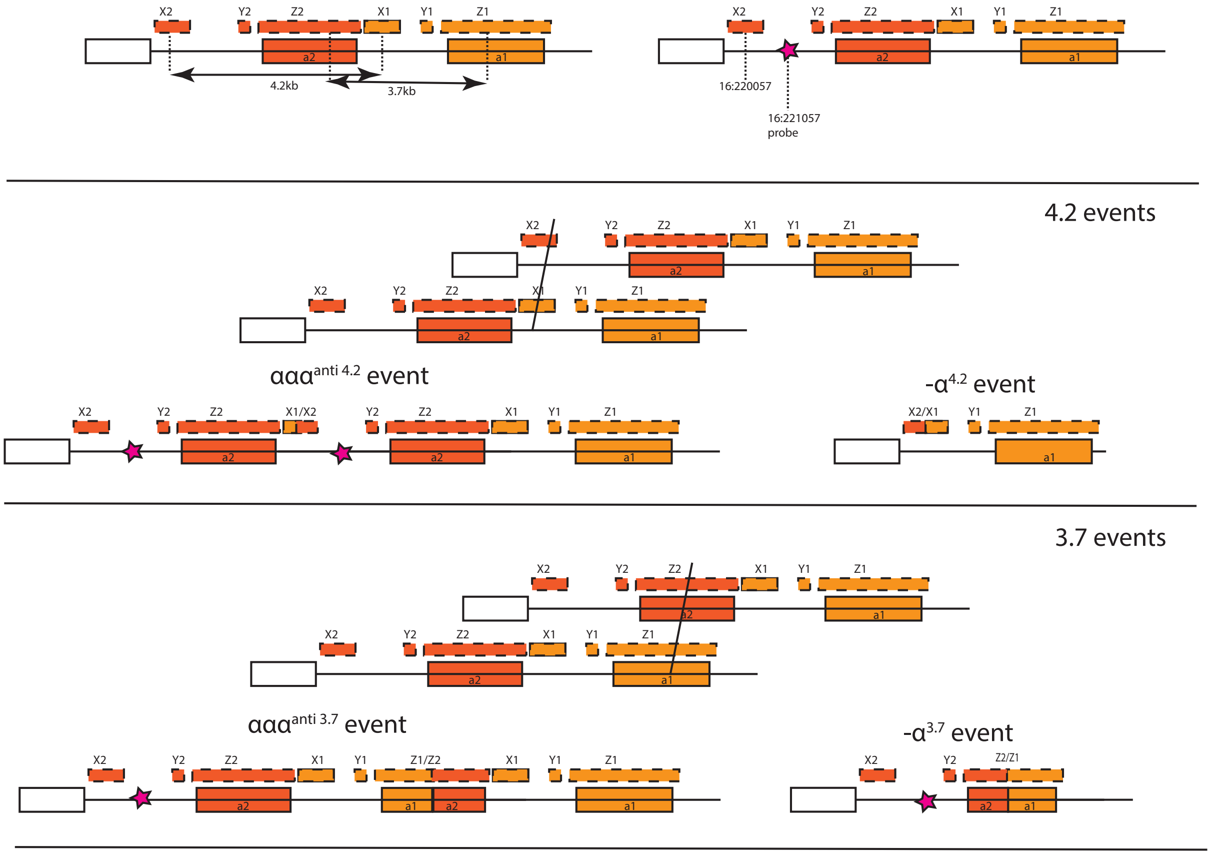

**Supplementary Figure 13. Mean LRR and normalized WES read depth for CNVs at the $\alpha$-globin locus.** Mean LRR across four probes (three within *HBA1* and 16:221057) and mean normalized read depth for exome-sequencing reads mapping uniquely to *HBA2* or *HBA1* are plotted for each category of $\alpha-$globin CNV we considered. Error bars, 95% CIs (computed as mean $\pm 1.96\times$ s.e.m. across carriers). WES read depth restricted to reads aligning uniquely to *HBA2* and *HBA1* (i.e., with nonzero mapping quality) was computed using mosdepth [37] and normalized by exome-wide coverage.

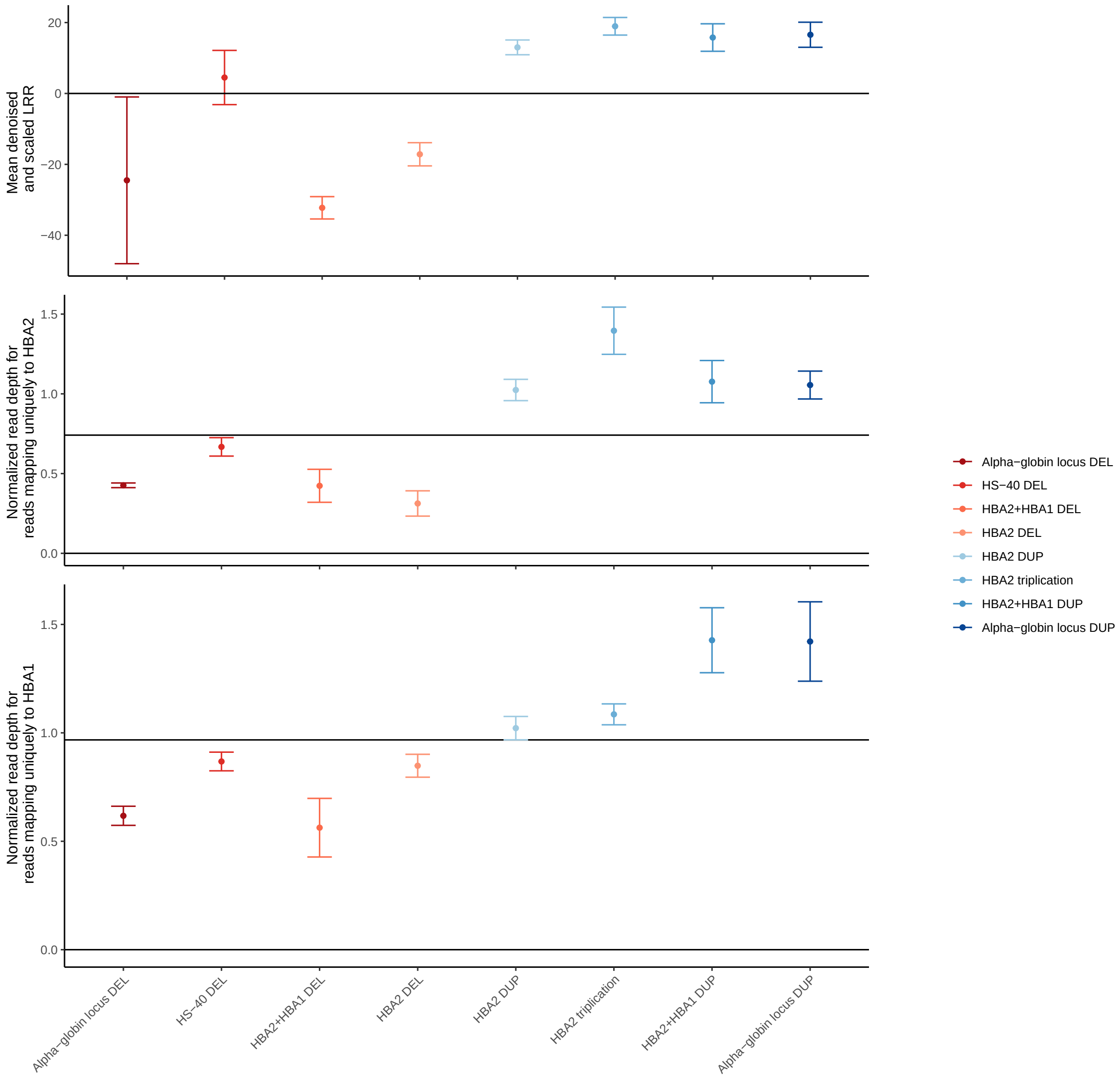

| | DEL | | DUP | |
|---|---|---|---|---|
| | Mean | SD | Mean | SD |
| HI-CNV | | | | |
| Total size (kb) | 429.94 | 308.72 | 899.26 | 641.28 |
| # CNV | 18.42 | 4.37 | 12.69 | 3.67 |
| HI-CNV$_0$ | | | | |
| Total size (kb) | 293.76 | 285.48 | 526.01 | 558.51 |
| # CNV | 6.52 | 2.71 | 3.43 | 2.07 |
| PennCNV | | | | |
| Total size (kb) | 299.02 | 374.88 | 722.51 | 1320.05 |
| # CNV | 2.67 | 1.77 | 2.45 | 1.94 |

**Supplementary Table 1. Summary of CNVs called per individual.** For each method, we report the mean and standard deviation of the number and total length of CNV calls per individual in our analysis set. HI-CNV$_0$ denotes analysis without incorporating information from haplotype neighbors (but still using our SNP-specific probabilistic models of genotyping intensities).

| DEL | | | | DUP | | | |
|---|---|---|---|---|---|---|---|
| Replicated | $RD_{sig}$ | $RD_{excess}$ | Sum | Replicated | $RD_{sig}$ | $RD_{excess}$ | Sum |
| HI-CNV | | | | | | | |
| 0.80 | 0.07 | 0.05 | 0.91 | 0.74 | 0.11 | 0.06 | 0.92 |
| $HI\text{-}CNV_0$ | | | | | | | |
| 0.91 | 0.06 | 0.02 | 0.98 | 0.85 | 0.09 | 0.06 | 1.00 |
| PennCNV | | | | | | | |
| 0.89 | 0.00 | -0.02 | 0.88 | 0.78 | 0.09 | 0.10 | 0.97 |

**Supplementary Table 2. Validation rate of SNP-array-based CNV calls assessed using WGS data available for 43 individuals.** For each method, we report the proportion of calls replicated using CNVnator, the proportion not called by CNVnator but with a significant deviation in read depth in the correct direction (e.g., significantly higher than average for duplications) ($RD_{sig}$), the difference in proportions with non-significant read depth in the correct vs. incorrect direction ($RD_{excess}$), and the sum of the previous three proportions, which we took to be the estimated validation rate. $HI\text{-}CNV_0$ denotes analysis without incorporating information from haplotype neighbors (but still using our SNP-specific probabilistic models of genotyping intensities).

| | DEL | | | DUP | | |
|---|---|---|---|---|---|---|
| | Size (kb) | | N | Size (kb) | | N |
| | Mean | SD | | Mean | SD | |
| HI-CNV | | | | | | |
| All | 23.34 | 68.10 | 8333810 | 70.84 | 167.47 | 5743712 |
| Distinct | 54.54 | 208.35 | 296822 | 148.94 | 481.05 | 247752 |
| $\text{HI-CNV}_0$ | | | | | | |
| All | 45.06 | 101.92 | 2950267 | 153.32 | 257.43 | 1552418 |
| Distinct | 63.66 | 216.96 | 275828 | 180.72 | 513.68 | 211270 |
| PennCNV | | | | | | |
| All | 112.04 | 195.55 | 1209343 | 295.32 | 744.30 | 1108613 |
| Distinct | 142.66 | 328.17 | 146533 | 328.24 | 1070.25 | 140690 |
| HI-CNV (unique CNVs, accounting for breakpoint uncertainty) | | | | | | |
| | 26.93 | 90.83 | 41486 | 80.91 | 182.85 | 23065 |

**Supplementary Table 3. Sizes and numbers of CNV calls.** For each method, we report the mean and standard deviation of CNV size (as well as the number of CNV calls over which these metrics were computed). For HI-CNV we computed metrics on the unioned callset (Supplementary Note 5.2). $\text{HI-CNV}_0$ denotes analysis without incorporating information from haplotype neighbors (but still using our SNP-specific probabilistic models of genotyping intensities). For each method, we computed metrics across all CNV calls (All) as well as distinct CNV calls (Distinct; i.e., counting calls with exactly the same breakpoints made in different individuals only once). For HI-CNV, we also computed metrics on the set of unique CNVs we identified after accounting for breakpoint uncertainty (Supplementary Note 7.3).

| | All CNVs | | | Genic CNVs | | |
|---|---|---|---|---|---|---|
| | Any size | $\geq$ 5kb | $\geq$ 5kb and $\geq$ 2 probes | Any size | $\geq$ 5kb | $\geq$ 5kb and $\geq$ 2 probes |
| HI-CNV | | | | | | |
| All | 0.06 | 0.37 | 0.81 | 0.24 | 0.50 | 0.82 |
| Del | 0.06 | 0.41 | 0.82 | 0.20 | 0.48 | 0.79 |
| Dup | 0.15 | 0.31 | 0.77 | 0.34 | 0.53 | 0.86 |
| HI-CNV$_0$ | | | | | | |
| All | 0.04 | 0.23 | 0.48 | 0.16 | 0.34 | 0.55 |
| Del | 0.04 | 0.26 | 0.51 | 0.14 | 0.33 | 0.53 |
| Dup | 0.09 | 0.17 | 0.42 | 0.23 | 0.36 | 0.58 |
| PennCNV | | | | | | |
| All | 0.01 | 0.07 | 0.16 | 0.05 | 0.12 | 0.21 |
| Del | 0.01 | 0.06 | 0.12 | 0.03 | 0.08 | 0.13 |
| Dup | 0.04 | 0.09 | 0.23 | 0.12 | 0.19 | 0.31 |

**Supplementary Table 4. Detection sensitivity (recall) of SNP-array based methods on a benchmark set of CNVs called from WGS data.** We assessed recall of low-frequency (MAF$\leq$5%) CNVs called by Delly in 43 individuals with whole-genome-sequence data available. Sensitivity increased with CNV size and probe overlap and for gene-overlapping CNVs. HI-CNV$_0$ denotes analysis without incorporating information from haplotype neighbors (but still using our SNP-specific probabilistic models of genotyping intensities).

| | Percentile | | | | | | |
|---|---|---|---|---|---|---|---|
| Type | Min | $5^{th}$ | $25^{th}$ | $50^{th}$ | $75^{th}$ | $95^{th}$ | Max |
| DEL | $6.11 \times 10^{-6}$ | $6.11 \times 10^{-6}$ | $9.78 \times 10^{-6}$ | $1.83 \times 10^{-5}$ | $4.15 \times 10^{-5}$ | 0.000275 | 0.137 |
| DUP | $6.11 \times 10^{-6}$ | $6.11 \times 10^{-6}$ | $9.78 \times 10^{-6}$ | $1.71 \times 10^{-5}$ | $4.03 \times 10^{-5}$ | 0.000426 | 0.115 |

**Supplementary Table 5. Minor allele frequency (MAF) distribution of unique CNV calls.** For deletions and duplications the minimum and maximum as well as the 5th, 25th, 50th, 75th, and 95th percentile of MAF is computed across unique CNV calls; unique CNVs were from the set of independent CNVs constructed on unrelated self-reported Europeans (see Section 7.3). The callset is depleted for common CNVs, probably due to a combination of SNP-array probe placement (avoiding common CNV regions) and less-accurate genotype cluster priors for SNPs within common copy-number polymorphisms.

| Genic context | Type | All unique CNVs $N$ | Prop. | Likely-causal CNVs $N$ | Prop. |
|---|---|---|---|---|---|
| No nearby gene | DEL | 12370 | 0.298 | 6 | 0.045 |
| | DUP | 5095 | 0.221 | 0 | 0.000 |
| Gene(s) within 100kb | DEL | 12372 | 0.298 | 11 | 0.082 |
| | DUP | 5752 | 0.249 | 6 | 0.053 |
| Intronic | DEL | 7596 | 0.183 | 2 | 0.015 |
| | DUP | 2212 | 0.096 | 0 | 0.000 |
| Genic | DEL | 9148 | 0.221 | 115 | 0.858 |
| 1 gene | | 7791 | 0.188 | 47 | 0.351 |
| 2 genes | | 855 | 0.021 | 24 | 0.179 |
| 3 genes | | 250 | 0.006 | 8 | 0.060 |
| 4 genes | | 91 | 0.002 | 5 | 0.037 |
| 5 genes | | 48 | 0.001 | 7 | 0.052 |
| >5 genes | | 113 | 0.003 | 24 | 0.179 |
| Genic | DUP | 10006 | 0.434 | 108 | 0.947 |
| 1 gene | | 6153 | 0.267 | 18 | 0.158 |
| 2 genes | | 1970 | 0.085 | 19 | 0.167 |
| 3 genes | | 829 | 0.036 | 10 | 0.088 |
| 4 genes | | 399 | 0.017 | 6 | 0.053 |
| 5 genes | | 199 | 0.009 | 2 | 0.018 |
| >5 genes | | 456 | 0.020 | 53 | 0.465 |

**Supplementary Table 6. Genic context of likely-causal deletions and duplications.** We compared the genic contexts of likely-causal CNVs to all unique CNVs (after accounting for breakpoint uncertainty; Supplementary Note 7.3), stratifying by CNV type (deletion or duplication).

| Genomic annotation | Proportion overlapping annotation<br>All unique CNVs | <br>Likely-causal CNVs | $P$ |
|---|---|---|---|
| H3K4me3_peaks_Trynka | 0.505 | 0.947 | 7.74e-05 |
| Human_Enhancer_Villar | 0.084 | 0.421 | 7.76e-05 |
| H3K4me3_Trynka | 0.536 | 0.947 | 0.000241 |
| H3K27ac_Hnisz | 0.583 | 0.947 | 0.0007 |
| H3K9ac_Trynka | 0.445 | 0.789 | 0.00415 |
| H3K27ac_PGC2 | 0.523 | 0.842 | 0.00518 |
| Enhancer_Hoffman | 0.240 | 0.526 | 0.00653 |
| H3K9ac_peaks_Trynka | 0.417 | 0.737 | 0.00846 |
| Promoter_UCSC | 0.073 | 0.263 | 0.00996 |
| FetalDHS_Trynka | 0.634 | 0.895 | 0.0169 |
| TSS_Hoffman | 0.090 | 0.263 | 0.0231 |
| PromoterFlanking_Hoffman | 0.139 | 0.316 | 0.0389 |
| Enhancer_Andersson | 0.105 | 0.263 | 0.0419 |
| TFBS_ENCODE | 0.664 | 0.895 | 0.0483 |
| WeakEnhancer_Hoffman | 0.313 | 0.526 | 0.0793 |
| Coding_UCSC | 0.025 | 0.105 | 0.0801 |
| BivFlnk | 0.091 | 0.211 | 0.088 |
| DHS_peaks_Trynka | 0.780 | 0.947 | 0.0963 |
| H3K4me1_peaks_Trynka | 0.783 | 0.947 | 0.0968 |
| CTCF_Hoffman | 0.292 | 0.474 | 0.126 |
| SuperEnhancer_Hnisz | 0.178 | 0.316 | 0.131 |
| DGF_ENCODE | 0.801 | 0.947 | 0.15 |
| DHS_Trynka | 0.795 | 0.947 | 0.151 |
| Conserved_LindbladToh | 0.626 | 0.789 | 0.162 |
| Ancient_Sequence_Age_Human_Enhancer | 0.040 | 0.105 | 0.175 |
| UTR_3_UCSC | 0.041 | 0.105 | 0.18 |
| H3K4me1_Trynka | 0.826 | 0.947 | 0.23 |
| UTR_5_UCSC | 0.048 | 0.105 | 0.234 |
| Vahedi_Tcell_TE | 0.058 | 0.105 | 0.302 |
| Vahedi_Tcell_SE | 0.021 | 0.053 | 0.327 |
| Vahedi_Tcell_SE_500bp | 0.021 | 0.053 | 0.329 |
| Vahedi_Tcell_TE_500bp | 0.062 | 0.105 | 0.329 |
| Conserved_Vertebrate_phastCons46way | 0.701 | 0.789 | 0.465 |
| Intron_UCSC | 0.335 | 0.421 | 0.469 |
| Conserved_Mammal_phastCons46way | 0.648 | 0.737 | 0.481 |
| Transcr_Hoffman.bed | 0.862 | 0.947 | 0.503 |
| Repressed_Hoffman | 0.822 | 0.789 | 0.763 |
| Conserved_Primate_phastCons46way | 0.470 | 0.474 | 1 |

**Supplementary Table 7. Enrichment of enhancer annotations within noncoding likely-causal CNVs.** For a set of binary annotations available from S-LDSC [38, 39], we computed the proportion of likely-causal noncoding deletions that overlapped the annotation, and we compared to the analogous proportion computed among all unique noncoding deletions. The horizontal line in the table indicates Bonferroni significance ($P$<0.05/38 = 0.0013).

| Variant | $n$ | $\beta$ (s.d. units) | s.e. | $P$ |
|---|---|---|---|---|
| pLoF CNVs | 19 | -1.107 | 0.170 | $8.2 \times 10^{-11}$ |
| SNP/indel PTVs | 9 | -1.033 | 0.248 | $3.04 \times 10^{-5}$ |

**Supplementary Table 8. Associations between *UHRF2* variants and height.** $n$, number of carriers. $\beta$, mean height (residualized for polygenic predictions from array-typed SNPs, omitting those within 2Mb) among carriers of *UHRF2* pLoF CNVs or SNP/indel PTVs (called in exome-sequenced individuals). s.e., standard error of $\beta$ (estimated as the standard deviation of the residualized phenotype across all UK Biobank participants divided by the square root of the number of carriers). $P$-value, $z$-test for nonzero $\beta$. (This $P$-value differs slightly from the BOLT-LMM $P$-value reported in Supplementary Data 2.)

| Trait | CNV | MAF | $\beta$ | s.e. | $P$ |
|---|---|---|---|---|---|
| Menarche age (yrs) | DEL | 0.004 | 0.200 | 0.034 | $5.8 \times 10^{-9}$ |
| Menarche age (yrs) | DUP | 0.020 | -0.117 | 0.015 | $1.6 \times 10^{-14}$ |
| Menarche age (sd) | DEL | 0.004 | 0.124 | 0.034 | $5.8 \times 10^{-9}$ |
| Menarche age (sd) | DUP | 0.020 | -0.073 | 0.015 | $1.6 \times 10^{-14}$ |
| Basophil # (sd) | DEL | 0.004 | -0.073 | 0.015 | $5.8 \times 10^{-7}$ |
| Basophil # (sd) | DUP | 0.019 | 0.023 | 0.007 | 0.00054 |
| Lymphocyte # (sd) | DEL | 0.004 | -0.061 | 0.015 | $1.1 \times 10^{-5}$ |
| Lymphocyte # (sd) | DUP | 0.019 | 0.028 | 0.007 | $1 \times 10^{-5}$ |
| Height (sd) | DEL | 0.004 | 0.038 | 0.012 | 0.0011 |
| Height (sd) | DUP | 0.019 | -0.034 | 0.006 | $2.5 \times 10^{-10}$ |

**Supplementary Table 9. Associations between *SLC2A3* CNVs and multiple phenotypes.** Minor allele frequency (MAF), effect size ($\beta$), standard error (s.e.), and $P$-value computed by BOLT-LMM are reported for *SLC2A3* deletions and duplications (specifically, 12:7996675-8123306 DEL/DUP, plus or minus two probes on either side; $\delta = 2$). Slight differences in MAF between phenotypes are due to MAF being computed in subsets of samples with non-missing phenotypes.

| Variant | $n$ | $\beta$ (s.d. units) | s.e. | $P$ |
|---|---|---|---|---|
| Upstream deletion | 19,483 | 0.118 | 0.006 | $9.24 \times 10^{-82}$ |
| pLoF CNVs | 8 | 0.298 | 0.302 | 0.325 |
| SNP/indel PTVs | 25 | 0.478 | 0.171 | 0.00519 |

**Supplementary Table 10. Associations between variants at the *BMP5* locus and bone mineral density.** Results are from linear regression of bone mineral density (residualized for polygenic predictions from array-typed SNPs, omitting those within 2Mb) on genotypes of an upstream deletion (6:55826807-55830155, plus or minus one probe on either side; $\delta = 1$), pLoF CNVs, and SNP/indel PTVs (called in exome-sequenced individuals). $P$-values are from $t$-tests for nonzero $\beta$. (This $P$-value differs slightly from the BOLT-LMM $P$-value reported in Supplementary Data 2.)

| Variant | $n$ | Mean corpuscular hemoglobin $\beta$ (s.d. units) | s.e. | $P$ | Red blood cell count $\beta$ (s.d. units) | s.e. | $P$ |
|---|---|---|---|---|---|---|---|
| $\alpha$–globin locus DEL | 2 | -3.411 | 0.600 | $1.31 \times 10^{-8}$ | 2.747 | 0.622 | $1 \times 10^{-5}$ |
| HS-40 DEL | 4 | -2.870 | 0.424 | $1.35 \times 10^{-11}$ | 2.737 | 0.440 | $4.87 \times 10^{-10}$ |
| *HBA2*+*HBA1* DEL | 34 | -2.645 | 0.146 | $8.2 \times 10^{-74}$ | 2.591 | 0.151 | $3.96 \times 10^{-66}$ |
| *HBA2* DEL | 21 | -1.728 | 0.185 | $1.04 \times 10^{-20}$ | 0.851 | 0.192 | $9.2 \times 10^{-6}$ |
| *HBA2* DUP | 58 | -0.214 | 0.111 | 0.0553 | 0.136 | 0.115 | 0.24 |
| *HBA2* triplication | 70 | -0.411 | 0.101 | $5.17 \times 10^{-5}$ | 0.343 | 0.105 | 0.00109 |
| *HBA2*+*HBA1* DUP | 34 | 0.142 | 0.146 | 0.328 | -0.025 | 0.151 | 0.868 |
| $\alpha$–globin locus DUP | 20 | -1.870 | 0.190 | $6.38 \times 10^{-23}$ | 0.892 | 0.197 | $5.68 \times 10^{-6}$ |

**Supplementary Table 11. Allelic series of CNVs at the $\alpha$-globin locus and effects on mean corpuscular hemoglobin (MCH) and red blood cell count (RBC).** $n$, number of carriers. $\beta$, mean MCH and RBC (residualized for polygenic predictions from array-typed SNPs, omitting those within 2Mb) among CNV carriers. s.e., standard error of $\beta$ (estimated as the standard deviation of the residualized phenotype across all UK Biobank participants divided by the square root of the number of carriers). $P$-value, $z$-test for nonzero $\beta$. (This $P$-value differs slightly from the BOLT-LMM $P$-value reported in Supplementary Data 2.)

| Variant | $n$ | $\beta$ (s.d. units) | s.e. | $P$ |
|---|---|---|---|---|
| Enhancer SNP | 231210 | 0.113 | 0.002 | $17.8 \times 10^{-684}$ |
| Enhancer deletion | 80 | 0.542 | 0.094 | $9.53 \times 10^{-9}$ |
| pLoF CNVs | 32 | 1.156 | 0.149 | $9.9 \times 10^{-15}$ |
| SNP/indel PTVs | 58 | 0.890 | 0.111 | $1.07 \times 10^{-15}$ |

**Supplementary Table 12. Associations between variants at the *JAK2* locus and platelet count.** Results are from linear regression of platelet count (residualized for polygenic predictions from array-typed SNPs, omitting those within 2Mb) on genotypes of the enhancer SNP rs12005199, deletions spanning this SNP, pLoF CNVs, and SNP/indel PTVs (called in exome-sequenced individuals). $n$, number of carriers. $P$-values are from $t$-tests for nonzero $\beta$. (This $P$-value differs slightly from the BOLT-LMM $P$-value reported in Supplementary Data 2.)

| **a** | | | Promoter capture Hi-C interactions | | | | |
|---|---|---|---|---|---|---|---|
| baitChr | baitStart | baitEnd | baitName | oeChr | oeStart | oeEnd | MK |
| 9 | 4977624 | 4986440 | *JAK2* | 9 | 4751287 | 4756519 | 25.353 |
| 9 | 4977624 | 4986440 | *JAK2* | 9 | 4756520 | 4772881 | 15.655 |
| 9 | 4977624 | 4986440 | *JAK2* | 9 | 4772882 | 4779795 | 6.846 |
| **b** | | | | Distal DHS data | | | |
| | Promoter DHS | | | | Distal DHS | | |
| chr | start | end | gene | chr | start | end | $\rho$ |
| 9 | 4727620 | 4727770 | *AK3* | 9 | 4755100 | 4755250 | 0.870 |
| 9 | 4727620 | 4727770 | *AK3* | 9 | 4762200 | 4762350 | 0.786 |
| 9 | 4983680 | 4983830 | *JAK2* | 9 | 4759420 | 4759570 | 0.795 |
| 9 | 4983680 | 4983830 | *JAK2* | 9 | 4760260 | 4760410 | 0.960 |
| 9 | 4983680 | 4983830 | *JAK2* | 9 | 4762200 | 4762350 | 0.701 |
| 9 | 4983680 | 4983830 | *JAK2* | 9 | **4763040** | **4763190** | **0.974** |
| **c** | | PTV analysis | | | | | |
| Gene | $n$ | $\beta$ (s.d.) | s.e. | $P$ | | | |
| *JAK2* | 58 | **0.882** | **0.111** | **2.45e-15** | | | |
| *AK3* | 40 | 0.023 | 0.134 | 0.864 | | | |

**Supplementary Table 13. Evidence for *JAK2* as the target of an enhancer ∼220kb upstream of *JAK2*.** We identified a large-effect platelet count association for deletions of a <4 kb focal region on chromosome 9 (4762652-4766114; Fig. 4b). This region contains an enhancer element containing a common SNP very strongly associated with platelet count that previous work has suggested may regulate *AK3*, the most proximal gene [40]; however, we found three lines of evidence suggesting that this element is actually a distal enhancer of *JAK2*. (**a**) Promoter capture Hi-C interactions suggest an interaction between *JAK2* and the region 9:4751287-4779795. The table reports CHiCAGO scores for all promoter capture Hi-C interactions involving *JAK2* or *AK3* in megakaryocytes (MK) (from PCHiC_peak_matrix_cutoff5.tsv from Supplementary Data of ref. [41]); genomic coordinates (hg19) indicate baited regions (baits) and the respective “other ends” (oe) of interactions (restricted to oe start or end $\geq$ 4.75Mb and $\leq$ 4.79Mb). (**b**) Distal DHS data show that a DHS within the enhancer region is highly correlated with the promoter of *JAK2*. For *AK3* and *JAK2*, the correlation ($\rho$) is given for all distal, non-promoter DHS within 500kb of the promoter DHS with correlation $\geq$0.7 (from Supplementary Table 7 of ref. [42]) that intersect chr9:4.75–4.79Mb. (**c**) PTV analyses show a strong association between *JAK2* loss-of-function and platelet count. $n$, number of carriers. $\beta$, mean platelet count (residualized for polygenic predictions from array-typed SNPs, omitting those within 2Mb) for carriers of SNP/indel PTVs called in exome-sequenced individuals. s.e., standard error of $\beta$ (estimated as the standard deviation of the residualized phenotype across all UK Biobank participants divided by the square root of the number of carriers). $P$-value, $z$-test for nonzero $\beta$.

| Variant | $n$ | $\beta$ (s.d. units) | s.e. | $P$ |
|---|---|---|---|---|
| Regulatory SNP (rs11642657) | 6371 | 0.389 | 0.012 | $2.9 \times 10^{-235}$ |
| Regulatory DEL (16:86011146-86014241) | 433 | 0.281 | 0.043 | $4.7 \times 10^{-11}$ |
| 10bp insertion (rs144009594) | 355714 | 0.102 | 0.002 | $7.8 \times 10^{-587}$ |
| pLoF CNVs | 5 | 1.099 | 0.397 | 0.0056 |
| SNP/indel PTVs | 5 | 0.772 | 0.397 | 0.052 |
| pLoF CNVs and SNP/indel PTVs | 10 | 0.935 | 0.281 | 0.00086 |

**Supplementary Table 14. Associations between variants at the *IRF8* locus and monocyte count.** Results are from linear regression of monocyte count (residualized for polygenic predictions from array-typed SNPs, omitting those within 2Mb) on genotypes of the downstream SNP rs11642657, a putatively regulatory downstream deletion (16:86011146-86014241, plus or minus two probes on either side; $\delta = 2$), a 10bp noncoding insertion in *IRF8*, pLoF CNVs, and SNP/indel PTVs (called in exome-sequenced individuals). The pLoF CNVs and SNP/indel PTV categories were analyzed both separately and merged into a single independent variable. $n$, number of carriers. $P$-values are from $t$-tests for nonzero $\beta$. (This $P$-value differs slightly from the BOLT-LMM $P$-value reported in Supplementary Data 2.)

| Variant | $n$ | $\beta$ (s.d. units) | s.e. | $P$ |
|---|---|---|---|---|
| Regulatory SNP (rs1683587) | 318379 | 0.041 | 0.002 | $6.59 \times 10^{-86}$ |
| Deletions in intron 1 | 79 | 0.449 | 0.101 | $8.74 \times 10^{-6}$ |
| pLoF CNVs | 58 | 0.528 | 0.118 | $7.5 \times 10^{-6}$ |
| SNP/indel PTVs | 79 | 0.520 | 0.101 | $2.73 \times 10^{-7}$ |
| Whole-gene duplications | 23 | 0.031 | 0.187 | 0.869 |

**Supplementary Table 15. Associations between *R3HDM4* variants and reticulocyte count.** Results are from linear regression of reticulocyte count (residualized for polygenic predictions from array-typed SNPs, omitting those within 2Mb) on genotypes of the intronic SNP rs1683587, deletions in intron 1 of *R3HDM4*, pLoF CNVs, SNP/indel PTVs (called in exome-sequenced individuals), and whole-gene duplications. $n$, number of carriers. $P$-values are from $t$-tests for nonzero $\beta$. (A probe-level test of deletions spanning 19:908648—which included both deletions in intron 1 and pLoF CNVs—reached genome-wide significance; Supplementary Data 2.)

**a Associations with platelet distribution width**

| Event | AF | $\beta$ | s.e. | $P$ |
|---|---|---|---|---|
| Top SNP (rs8001925) | 0.653 | 0.060 | 0.002 | 2.5e-172 |
| Top eQTL (rs842368) | 0.564 | 0.043 | 0.002 | 4.1e-98 |
| Insertion (typed by Affx-52351109) | 0.007 | 0.099 | 0.012 | 3.5e-17 |

**b Associations with *LRCH1* expression**

| | Insertion | | | Top SNP | | Top eQTL | |
|---|---|---|---|---|---|---|---|
| Tissue | $P$ | $\beta$ | $n$ | $P$ | $\beta$ | $P$ | $\beta$ |
| Lung | 0.27 | -0.160 | 8 | 0.94 | -0.000 | 0.52 | -0.020 |
| Muscle_Skeletal | 0.012 | -0.470 | 8 | 6.6e-05 | -0.110 | 5.2e-05 | -0.110 |
| Skin_Not_Sun_Exposed_Suprapubic | 0.0014 | -0.340 | 8 | 0.043 | -0.050 | 0.052 | -0.040 |
| Thyroid | 0.0094 | -0.480 | 8 | 0.21 | -0.040 | 0.021 | -0.070 |
| Whole_Blood | 0.33 | -0.100 | 8 | 5.8e-05 | -0.080 | 2.4e-05 | -0.080 |
| Adipose_Subcutaneous | 0.0093 | -0.330 | 7 | 5.9e-11 | -0.150 | 2.1e-05 | -0.100 |
| Artery_Tibial | 0.0033 | -0.320 | 7 | 0.034 | -0.040 | 0.095 | -0.040 |
| Breast_Mammary_Tissue | 0.21 | -0.150 | 7 | 0.00091 | -0.090 | 0.00082 | -0.090 |
| Esophagus_Muscularis | 0.16 | -0.170 | 7 | 0.00019 | -0.080 | 0.00025 | -0.070 |
| Heart_Atrial_Appendage | 0.81 | -0.040 | 7 | 0.046 | -0.070 | 0.053 | -0.070 |
| Nerve_Tibial | 0.21 | -0.200 | 7 | 5.3e-08 | -0.170 | 4.2e-06 | -0.140 |
| Skin_Sun_Exposed_Lower_leg | 0.00097 | -0.390 | 7 | 0.91 | -0.000 | 0.15 | -0.030 |
| Artery_Aorta | 0.067 | -0.280 | 6 | 9e-05 | -0.130 | 5.9e-06 | -0.150 |
| Esophagus_Mucosa | 0.00015 | -0.640 | 6 | 0.24 | -0.030 | 0.28 | -0.030 |
| Heart_Left_Ventricle | 0.44 | -0.140 | 6 | 7.5e-14 | -0.270 | 8.3e-18 | -0.300 |
| Pancreas | 0.027 | -0.600 | 6 | 0.46 | -0.040 | 0.1 | -0.090 |
| Adipose_Visceral_Omentum | 0.0024 | -0.430 | 5 | 0.012 | -0.070 | 0.73 | -0.010 |
| Brain_Cerebellum | 0.24 | -0.250 | 5 | 0.13 | -0.090 | 0.075 | -0.110 |
| Colon_Transverse | 0.77 | 0.040 | 5 | 0.39 | -0.020 | 0.31 | -0.030 |
| Ovary | 0.96 | 0.010 | 5 | 0.19 | -0.090 | 0.86 | -0.010 |
| Uterus | 0.046 | -0.550 | 5 | 0.47 | -0.060 | 0.14 | -0.110 |
| Adrenal_Gland | 0.24 | -0.350 | 4 | 0.8 | 0.010 | 0.28 | -0.060 |
| Cells_Cultured_fibroblasts | 0.00043 | -0.560 | 4 | 0.0026 | -0.060 | 0.00076 | -0.070 |
| Esophagus_Gastroesophageal_Junction | 0.056 | -0.360 | 4 | 0.42 | -0.020 | 0.019 | -0.070 |
| Stomach | 0.4 | 0.190 | 4 | 0.71 | -0.010 | 0.31 | -0.040 |
| Mean across tissues | | -0.283 | | | -0.070 | | -0.078 |

**Supplementary Table 16. Associations of *LRCH1* variants with platelet distribution width and *LRCH1* expression. a** Allele frequencies (AF), platelet distribution width (PDW) effect sizes and standard errors, and association $P$-values for: the SNP most strongly associated with PDW (among $N$=460K UK Biobank participants of self-reported European ancestry), the top eQTL (from GTEx), and the retroposed *MTMR2* insertion into *LRCH1*. ($P$-values from linear regression are reported because mixed model analysis of *LRCH1* SNPs on chromosome 13 inadvertently partially conditioned on the insertion variant typed by Affx-52351109, which was coded as being on chromosome 11 instead of chromosome 13). **b** Associations with *LRCH1* expression in 25 tissues for which GTEx RNA-seq data was available for at least $n \geq 4$ carriers of the insertion. $P$-values and effect sizes ($\beta$) were computed using FastQTL [43].

| Type | Syndromic CNVs included $\beta$ (s.e.) | $P$ | Syndromic CNVs excluded $\beta$(s.e.) | $P$ |
|---|---|---|---|---|
| | | Height (s.d.) | | |
| DEL (Mb) | -0.051 (0.005) | $8.4 \times 10^{-25}$ | -0.042 (0.005) | $4.7 \times 10^{-15}$ |
| DUP (Mb) | -0.014 (0.002) | $5.9 \times 10^{-9}$ | -0.011 (0.003) | $1.5 \times 10^{-5}$ |
| | | BMI (s.d.) | | |
| DEL (Mb) | 0.036 (0.005) | $1.8 \times 10^{-12}$ | 0.031 (0.005) | $1.9 \times 10^{-8}$ |
| DUP (Mb) | 0.008 (0.002) | 0.0013 | 0.004 (0.003) | 0.13 |
| | | Years of education | | |
| DEL (Mb) | -0.28 (0.025) | $1.1 \times 10^{-28}$ | -0.222 (0.027) | $2.5 \times 10^{-16}$ |
| DUP (Mb) | -0.071 (0.012) | $5.1 \times 10^{-9}$ | -0.045 (0.013) | 0.0006 |

**Supplementary Table 17. Deletion and duplication burden on three quantitative traits.** We restricted analysis to unrelated UK Biobank participants of self-reported European ancestry; within this subset we mean-standardized height and BMI (both in units of standard deviations). We then ran linear regression between the DEL or DUP burden (in Mb) and quantitative traits of interest (height, BMI, and years of education), controlling for age, sex, and 20 PCs. We ran two sets of analyses, one allowing syndromic CNVs ($N$=409252) and one excluding carriers of any syndromic CNV ($N$=392643).

| CNV type | Estimation method | Relative effect size estimate | L | U |
|---|---|---|---|---|
| DUP | MLE | -0.212 | -0.248 | -0.176 |
| DEL | MLE | 0.899 | 0.798 | 1.000 |
| DUP | MLE (absolute value) | 0.295 | 0.259 | 0.331 |
| DEL | MLE (absolute value) | 1.005 | 0.904 | 1.106 |
| DUP | PoiBinom | 0.364 | 0.350 | 0.379 |
| DEL | PoiBinom | 0.864 | 0.814 | 0.915 |

**Supplementary Table 18. Estimation of effect sizes of duplications and deletions relative to PTVs.** For MLE approaches, the estimate reported is the maximum likelihood estimate along with the 95% confidence interval. For the Poisson-Binomial approach the value reported is the one which results in an expected number of significant associations closest to what is observed; the upper and lower values are the ones which result in an expected number of significant associations closest to what is observed $\pm$ 1.

| # of PCs used | Fraction of LRR variance explained |
|---|---:|
| 10 | 0.441 |
| 20 | 0.522 |
| 30 | 0.555 |
| 40 | 0.573 |
| 50 | 0.585 |
| 60 | 0.594 |
| 70 | 0.601 |
| 80 | 0.607 |
| 90 | 0.611 |
| 100 | 0.616 |

**Supplementary Table 19. Average fraction of LRR variance explained by top 10, 20, . . ., 100 principal components.** For each of the 106 genotyping batches in turn, principal components were computed on LRR values for all autosomal variants, restricting to white British samples. The average fraction (across all batches) of LRR variance explained by the top 10, 20, . . ., 100 PCs was computed.

| | Fraction of $n$-th longest haplotype matches exceeding the indicated length | | | | | | | | | |
|---|---|---|---|---|---|---|---|---|---|---|
| Neighbor | >1cM | >2cM | >3cM | >4cM | >5cM | >6cM | >7cM | >8cM | >9cM | >10cM |
| 1 | 0.980 | 0.931 | 0.885 | 0.841 | 0.796 | 0.752 | 0.708 | 0.663 | 0.620 | 0.578 |
| 2 | 0.967 | 0.896 | 0.826 | 0.761 | 0.697 | 0.632 | 0.569 | 0.510 | 0.454 | 0.402 |
| 3 | 0.957 | 0.868 | 0.782 | 0.700 | 0.621 | 0.548 | 0.478 | 0.411 | 0.351 | 0.300 |
| 4 | 0.948 | 0.843 | 0.743 | 0.650 | 0.562 | 0.481 | 0.408 | 0.340 | 0.282 | 0.231 |
| 5 | 0.940 | 0.821 | 0.710 | 0.608 | 0.513 | 0.429 | 0.355 | 0.289 | 0.233 | 0.185 |
| 6 | 0.933 | 0.801 | 0.679 | 0.570 | 0.472 | 0.388 | 0.312 | 0.249 | 0.195 | 0.152 |
| 7 | 0.926 | 0.782 | 0.650 | 0.536 | 0.436 | 0.350 | 0.278 | 0.216 | 0.166 | 0.128 |
| 8 | 0.919 | 0.764 | 0.625 | 0.506 | 0.404 | 0.320 | 0.248 | 0.189 | 0.144 | 0.108 |
| 9 | 0.911 | 0.747 | 0.602 | 0.479 | 0.376 | 0.292 | 0.223 | 0.168 | 0.125 | 0.093 |
| 10 | 0.905 | 0.732 | 0.577 | 0.453 | 0.351 | 0.268 | 0.200 | 0.150 | 0.109 | 0.079 |

**Supplementary Table 20. Lengths of haplotypes shared by top "haplotype neighbors."** These data describe the length distribution (across SNPs on the array and the two haplotypes of each UK Biobank participant) of the $n$-th longest haplotype match. For example, the longest match ("closest haplotype neighbor") is $>$1cM 98% of the time, $>$5cM 80% of the time, and $>$10cM 58% of the time.
